# Transposon-mediated telomere destabilization: a driver of genome evolution in the blast fungus

**DOI:** 10.1101/845669

**Authors:** Mostafa Rahnama, Olga Novikova, John Starnes, Li Chen, Shouan Zhang, Mark Farman

## Abstract

*Magnaporthe oryzae* is a filamentous ascomycete fungus that causes devastating diseases of crops that include rice and wheat, and a variety of turf, forage and wild grasses. Strains from ryegrasses possess highly stable chromosome ends that undergo frequent rearrangements during vegetative growth in culture and *in planta*. Instability is associated with the presence of two related retrotransposons (*<u>M</u>agnaporthe <u>o</u>ryzae* <u>Te</u>lomeric <u>R</u>etrotransposons - MoTeRs) inserted within the telomere repeat tracts. The objective of the present study was to determine the mechanisms by which MoTeRs promote telomere instability. Targeted cloning, restriction mapping, and sequencing of both parental and novel telomeric restriction fragments, along with MinION sequencing of DNA from three single-spore cultures, allowed us to document the molecular alterations for 109 newly-formed telomeres. Rearrangement events included truncations of subterminal rDNA sequences; acquisition of MoTeR insertions by “plain” telomeres; insertion of the MAGGY retrotransposons into MoTeR arrays; expansion and contraction of subtelomeric tandem repeats; MoTeR truncations; duplication and terminalization of internal sequences; and breakage at long, interstitial telomeres generated during MoTeR insertion. Together, our data show that when MoTeRs invade the telomeres, they can dramatically perturb the integrity of chromosome ends, leading to the generation of unprotected DNA termini whose repair has the potential to generate chromosome alterations that extend well into the genome interior.

## INTRODUCTION

*Magnaporthe oryzae* is an ascomycete fungus that causes blast disease in rice, wheat and other crops; and is also responsible for leaf spot diseases of a variety of turf and pasture grasses, including annual ryegrass [1, 2], perennial ryegrass [3], tall fescue and St. Augustinegrass [4]. Most *M. oryzae* strains exhibit host specificity, being capable of infecting only a very small number of host genera and/or species [5, 6]. Additional specificity exists at the sub-species level, with rice, wheat and foxtail pathogens being compatible with some cultivars of their respective host species, and not others [7–9]. Cultivar specificity has been intensively study because it is the main foundation upon which plants are bred for blast resistance. Unfortunately, cultivar specificity often breaks down due to a high degree of pathogenic variability within the fungus [10, 11]. Studies at the molecular level have shown that *M. oryzae* escapes host and cultivar recognition through the mutation (or loss) of genes that code for proteins that are secreted during infection [12, 13]. These proteins would normally trigger resistance in host plants that contain the corresponding resistance receptors and, for this reason, they are termed “avirulence” effectors. Interestingly, many avirulence genes exhibit a high degree of genetic instability and a large proportion of them (∼ 50%) map very close to telomeres [14].

Telomeres are the sequences that constitute the ends of linear chromosomes and in most organisms comprise short, tandem repeats - (TTAGGG)_n_ in most fungi (incl. *M. oryzae*) and animals [15]; and (TTTAGGG)_n_ in plants [16]. In many organisms, the regions immediately adjacent to the telomeres contain sequences that are duplicated at different chromosome ends - effectively creating a defined subtelomeric domain [17]. The distal subtelomere regions usually contain a variety of short, tandem repeat motifs [18–23], and the proximal portions often harbor families of genes which are associated with niche adaptation [24–30]. Because the chromosome ends are the most dynamic regions of the genome - often undergoing spontaneous rearrangements [29, 31–36], and experiencing accelerated mutation [37–39], and epigenetic silencing [29, 32, 40–43] - they tend to exhibit much higher levels of polymorphism than the genome interior [37, 38, 44–50].Accordingly, the genes that reside there benefit from the enhanced evolutionary and adaptive potential afforded by these behaviors [31, 35, 51–53].

Previously, we reported that telomeric restriction fragments (TRFs) in *M. oryzae* strains from perennial ryegrass are unusually polymorphic when compared to internal chromosomal regions [45]. This contrasts with the telomeres of strains from rice, which are remarkably stable by comparison [54]. Characterization of unstable telomeres revealed evidence that this polymorphism is due to frequent, spontaneous rearrangements at the chromosome ends and that this instability is associated with the presence of non-LTR retrotransposons embedded within the telomere repeats [54]. Two related retroelements were identified. The first, MoTeR1, is 5 kb in length and codes for a predicted reverse transcriptase (RT) enzyme that exhibits similarity to RTs encoded by the SLACS retrotransposon from *Giardia lambliae* [55], and CRE1 from *Crithidia faciculata* [56, 57]. These latter elements are site-specific transposons that insert specifically into splice leader sequence genes, using a restriction enzyme-like endonuclease (REL-ENDO). The MoTeR1 RT contains a putative REL-ENDO domain and possesses an extensive run of telomere-like sequence, TTCGGG(TTTGGG)n, at its 5’ terminus [54], which leads us to suspect that MoTeR1 is a site-specific transposon that targets telomere repeats in a similar manner to the TRAS and SART retrotransposons in *Bombyx mori* [58].

MoTeR2 shares the same 5’ (860 bp) and 3’ (77 bp) terminal sequences with MoTeR1. However, in MoTeR2, the RT coding region is replaced with an unrelated 786 bp sequence with no obvious function [54]. Thus, MoTeR2 is likely a defective element and, if mobile, probably uses the MoTeR1 RT for its transposition. MoTeR1 and MoTeR2 sequences can exist in single copy in a given telomere, or in homogeneous, or heterogeneous arrays. When they occur in tandem, adjacent elements are always arranged in a head to tail orientation, and are separated by a (TTAGGG)_n_ tract ranging in length from one half of a repeat unit to more than 20 [54].

Given that MoTeR1 bears all the hallmarks of a site-specific transposon, we hypothesized that the abundant rearrangements observed during vegetative growth *in vitro* and *in planta* [54] might reflect frequent transposition events. To test this idea, we first employed shotgun and targeted cloning to characterize a number of newly-formed TRFs, along with the respective chromosome ends in the progenitor strain. Next, we used MinION sequencing to uncover a large number of cryptic telomere alterations. Together, these efforts allowed us to document a wide range of molecular mechanisms giving rise to more than 100 telomere rearrangements in *M. oryzae*.

## METHODS

### Passaging the fungus through plants

Two experiments were performed in which the fungus was serially passaged through plants two times, with no artificial culture in between (Fig. S1). For experiment 1, strain LpKY97 was reactivated from a stock culture by placing on oatmeal agar. Spores were harvested by flooding the plate with a 0.2% gelatin solution and spayed on leaves of the annual ryegrass cultivar Gulf. For experiment 2, a single spore was isolated from the original LpKY97 culture and used to establish a single spore (SS) culture on oatmeal agar. A spore suspension was made and sprayed on leaves of the perennial ryegrass cultivar Linn. Seven days later, after disease symptoms had appeared, leaves with visible lesions were removed, placed in a moist chamber and incubated overnight at room temperature for sporulation to occur (∼ 25°). The spores from this second generation were harvested by flooding the plate with a 0.2% gelatin solution and a small aliquot was streaked on water agar, to establish second generation SS cultures. The reminder was used to re-inoculate plants and the process was repeated to generate a third SS generation. Finally, select third generation SS cultures were re-cultured on oatmeal agar and up to 20 single spores were collected and used to generate fourth generation cultures. The various cultures were named according to the following scheme: <experiment#>SS<single-spore-IDs#> (e.g. 2G4SS1-10 indicates experiment 2, 4^th^ generation spore culture #10, derived from 3^rd^ generation spore culture #1).

### Generation of single spore isolates from fungus grown on oatmeal agar plates

For experiment #3, a single spore of strain LpKY97-1 was inoculated at the very edge of a Petri dish containing complete medium agar (CMA). This medium allows rapid growth but suppresses sporulation which ensures that the fungus undergoes a maximal number of nuclear divisions in reaching the other side of the plate - instead of jumping via spores. The mycelium was allowed to grow to the opposite side of the plate, at which point a plug containing the mycelial front was excised and used to inoculate a second CMA plate. After the mycelium had grown to the other side of the second plate, a plug containing the mycelial front was placed on oatmeal agar to induce sporulation. Spores were then harvested by gently brushing a sealed Pasteur pipette over the culture and the pipette tip was then streaked across water agar plates. After overnight incubation at room temperature, over 250 germinated spores were individually isolated and used to start a set of “3G3” SS cultures.

### Shotgun cloning of telomeric restriction fragments

Genomic DNA was purified using a standard procedure [54] and polysaccharides were subsequently removed using differential ethanol precipitation [59]. Approximately 5 µg of polysaccharide-reduced, undigested genomic DNA was end-repaired using the End-It^TM^ kit (Epicentre Technologies, Madison, WI). The enzyme was removed by phenol:chloroform:isoamyl alcohol (PCI, 25:24:1) extraction (2x), followed by chloroform:IAA (CI, 24:1) extraction (1x) and the DNA was then ethanol-precipitated, rinsed with 70% ethanol, dried and re-dissolved in 9 µl of 1x ligation buffer (New England Biolabs, Ipswich, MA). To this was added: 1 µl of *Eco*RV-digested pBlueScript KS II^+^ (∼100 ng) that had been treated with calf intestinal phosphatase (Promega Corp., Madison, WI), and 0.1 µl of T4 DNA ligase (NEB). Ligation was performed overnight at 12°C. The ligase was killed by heating at 65°C for 10 min and then 5 µl of 10x reaction buffer 3 (NEB), 34 µl of H_2_O and 1 µl of *Pst*I (NEB) were added and restriction digestion was performed at 37°C for 4 h. After digestion, the restriction enzyme was removed by PCI and CI extraction and the DNA was ethanol-precipitated. The DNA pellet was re-dissolved in 50 µl of 1x ligation buffer, 0.1 µl (40 u) of ligase (NEB) was added and the reaction was allowed to proceed at 12°C overnight. Finally, the DNA was ethanol-precipitated and re-dissolved in 20 µl of TE. The ligated DNA sample was used to transform electrocompetent *E.coli* cells (EPI300, Epicentre), using 1 µl of ligated DNA solution per transformation. Telomere-positive clones were identified by colony blotting using a published procedure [60, 61].

### Targeted cloning of individual telomeric restriction fragments

The method used to clone specific TRFs through enrichment of terminal sequences has been described in detail elsewhere [60]. Briefly, polysaccharide-depleted, genomic DNA samples (∼2 µg) were end-repaired using the End-it kit from Epicentre Technologies (Madison, WI). The enzymes were inactivated and removed using PCI (2x) and CI (1x) extraction and the DNA was ethanol-precipitated and rinsed in 70% ethanol. The DNA was then dissolved in 1x restriction enzyme buffer and digested with *Pst*I for 4 h before being fractionated by gel electrophoresis. Gel slices containing DNA molecules in size ranges spanning the target fragments were excised and the DNA was extracted using the QIAquick kit (Qiagen, Valencia, CA) and eluted in 25 µl elution buffer. The purified fragments (3.5 µl, ∼ 100 ng) were ligated to *Pst*I+*Eco*RV-digested, CIP-treated pBlueScript KS II^+^ (∼100 ng) in a reaction volume of 5 µl. One microliter of the reaction mix was used to transform ultra-competent EPI300 cells (Epicentre). Telomere-positive clones were identified by colony blotting.

### Southern blotting and hybridization

The methods used for Southern blotting and generation of telomere probes have been described previously [54]. Individual telomeres and their corresponding telomeric restriction fragments (TRFs) were named according to their correspondence with the telomeres in the reference genome for strain B71 [62]. Telomeres on supernumary (mini-) chromosomes were labeled using A, B, C and D identifiers, with the two mini-chromosomes being labeled 1 and 2 according to their sizes (largest first). Newly-formed telomeres/TRFs that could not be associated with specific telomeres retained the lowercase a, b, c, etc. identifiers that were initially assigned to the novel TRFs.

Telomere-linked probes (TLPs) were developed by sequencing the cloned TRFs using T3 and T7 primers, and a primer reading out from the MoTeR 3’ end (MoTeRnR). Where possible, sequences adjacent to multi-MoTeR arrays were acquired using 3’ end-proximal primers that were specific to either MoTeR1 or MoTeR2. Chromosomal sequences adjacent to the MoTeR and the non-telomeric end of the TRF insert were compared with known *M. oryzae* repeats, and primers were designed to amplify single-copy (or low-copy) probes. Primer sequences are listed in Table S1.

### MinION sequencing and analysis

Between 2 - 5 µg of DNA, extracted as described above, was further purified using the MagAttract kit (Qiagen). Sequence-ready libraries were then prepared from 500 ng to 1 µg of intact (non-sheared) DNA using the SQK-LSK109 Ligation Sequencing Kit (Oxford MinION Technologies, Oxford, UK) according to the manufacturer’s instructions. Sequence data were acquired on MinION flow cells for between 16 and 24 h. Raw data in fast5 format were converted to fastq using Guppy [63] and individual strain assemblies were generated using canu [64], with default settings. To generate a high quality, base reference for strain LpKY97, a hybrid assembly was created with data from three descendant progeny, using only reads that were 8 kb in length, or longer and single nucleotide errors were corrected using an Illumina assembly. Telomere rearrangements were identified by aligning individual reads to the individual and merged reference genomes using minimap2 [65], followed by manual inspection of the alignments in the Integrative Genomics Viewer [66]. Specifically, we examined only those alignments where the reads extended from the unique, telomere adjacent region all the way to the telomere. In this case, we required that the reads contained 10 to 30 TTAGGG repeats that extended to an end of the sequenced molecule (with an allowance for up to approximately 30 nt of sequencing adaptor that was present at the ends of some reads).

## RESULTS

### Terminology

#### Telomeres

The sequences that comprise the chromosome ends - in this case, repeats of the hexanucleotide motif, CCCTAA/TTAGGG.

#### Telomere-adjacent sequences

The sequences immediately subtending the telomeric repeats. *Subtelomeres.* A term reserved for specific domains that comprise sequences duplicated at multiple chromosome ends, and not usually found elsewhere in the genome.

#### Subterminal sequences/regions

Sequences/regions near the chromosome ends that do not qualify as “subtelomeres,” either because they are unique, or they comprise sequences duplicated throughout the genome. Arbitrarily, we define the subterminal regions as being within 100 kb from the chromosome ends.

#### Chromosome-unique sequence

Sequence from a subterminal region that uniquely identifies it as coming from a specific chromosome end.

### Identification of *M. oryzae* cultures with altered telomere “fingerprints.”

To explore the molecular events responsible for the frequent telomere rearrangements in MoTeR-containing strains, we established a large collection of isolates containing one or more newly-formed telomeric restriction fragments (TRFs) (Fig. S1). The first set of 20 single-spore isolates came from an experiment in which the perennial ryegrass pathogen, LpKY97 was passaged through annual ryegrass plants for two successive generations. Six of 10 isolates collected after a single infection cycle and 10 of 10 collected after two cycles exhibited novel telomere profiles compared with the original parental strain (Fig. 1). Most of the fragments less than 10 kb in length showed rearrangements in at least one single spore culture. However, there was a notable exception in the 5.7 kb TRF (“TEL5,” Fig. 1). This fragment was relatively stable and exhibited rearrangement in only two out of more than 250 single spore isolates that were analyzed (Fig. 1, Fig. S2; and data not shown). Isolates containing novel TRFs were selected for further investigation.

**Fig. 1.**
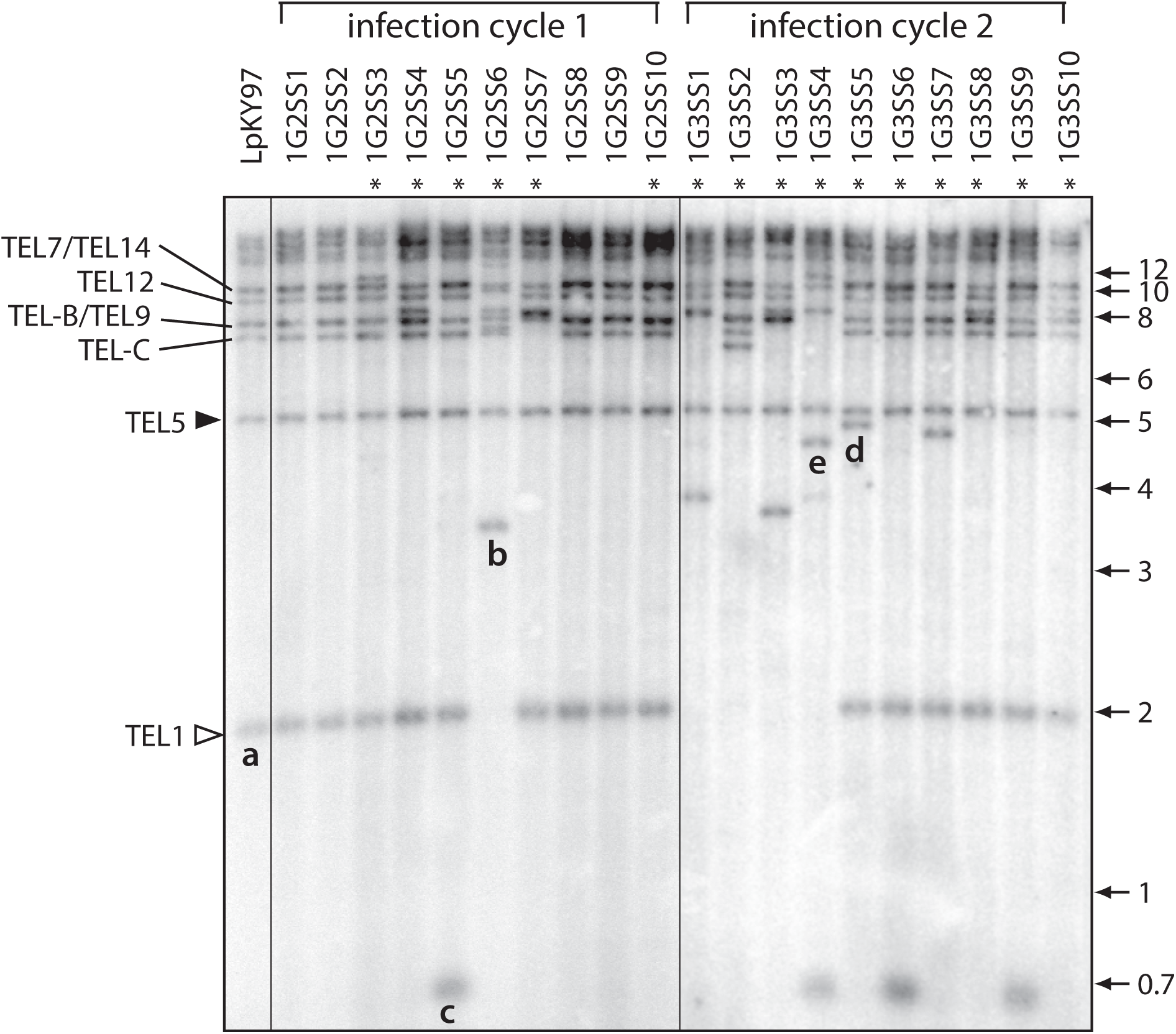
Characterization of rearranged telomeric restriction fragments (TRFs) formed during growth *in planta*. A) Telomeric restriction profiles of single spore isolates from successive rounds of plant infection. DNA samples were digested with *Pst*I, fractionated by gel electrophoresis, blotted to a membrane and hybridized with a telomere probe. TRFs are labeled based on their chromosome locations in the *M. oryzae* B71 reference genome. Asterisks mark lanes with telomere profiles that are different from the parent culture. An open arrowhead marks the highly unstable “rDNA” telomere (TEL1). A closed arrowhead marks a relatively stable telomere that rarely exhibited rearrangement (TEL5). Telomeres on mini-chromosomes 1 and 2 are labeled TEL-A and TEL-C, respectively. Novel fragments that were successfully cloned and characterized are labeled “a” through “e.”

First, we used a shotgun strategy to clone and sequence *Pst*I TRFs from the single spore cultures exhibiting telomere alterations. In parallel, we attempted to clone *Pst*I TRFs from the original parent strain, LpKY97. Consistent with their terminal position on the chromosome, all of the cloned TRFs contained telomere repeats ligated directly to the *Eco*RV half-site in the vector, with the C-rich strand reading in the 5’ to 3’ direction.

### “rDNA telomere” truncations

One of the first telomeres to be cloned was contained on a ∼2 kb *Pst*I fragment (Fig. 1, band “a”) and corresponded to telomere 1 (TEL1). The cloned fragment consisted of ribosomal DNA (rDNA) sequence with telomere repeats attached at a position 1,567 bp downstream of the 26S rRNA gene (Fig. 2A), or nucleotide position 7,581 in the *M. oryzae* rDNA reference sequence (NCBI #AB026818). Among 39 single-spore progeny isolated following plant infection, 13 lacked the 2 kb TEL1 fragment (1G2SS6, 1G3SS1, 2, 3 & 4 [Fig. 1], 2G3SS4, 5, 6, 8 & 9 [Fig. S2], and 3G1SS56 [data not shown]). Altered versions of the rDNA telomere were successfully captured from shotgun and targeted libraries for seven of these cultures. In five cases, the telomere repeats were attached to different positions within the rRNA genes, causing the size of the TEL1 fragment to increase (Fig. 2A; see band “b” in Fig. 1 & 3A). These rearrangements are attributable to simple truncations of the rDNA array (note that while array extensions might be involved, the initiating events for extensions most likely will be telomere loss followed by array truncations). One culture (1G3SS4) experienced more a complex TEL1 rearrangement (see below).

**Fig. 2.**
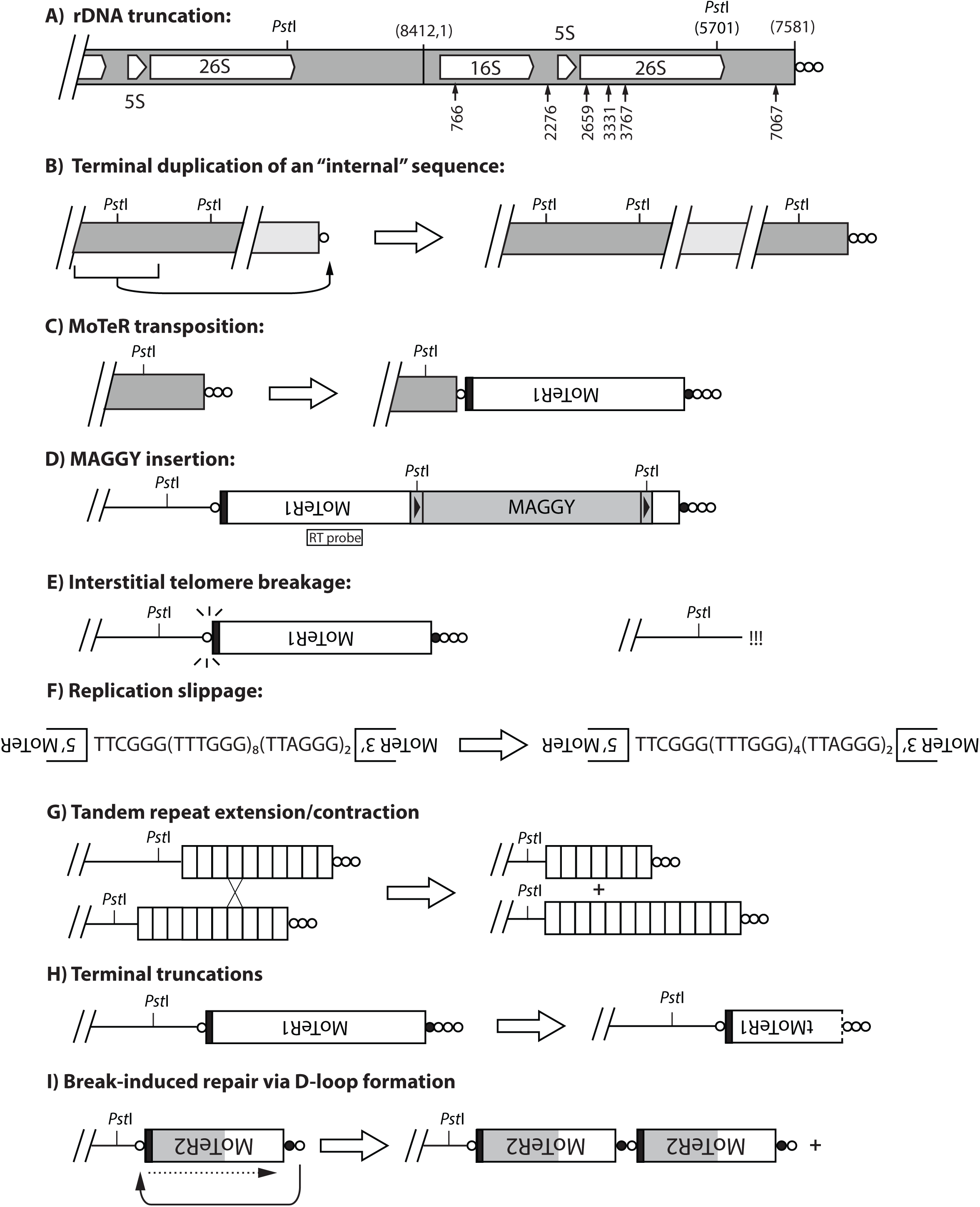
Mechanisms giving rise to rearranged TRFs. A) *Truncations of the ribosomal DNA array* (TEL1). The positions of 18S, 5S and 26S rRNA genes are shown, along with the positions of the two most distal *Pst*I restriction sites. The telomere repeats are denoted as circles with each circle representing 1 to 10 copies of TTAGGG - (TTAGGG)_1-10_. Relevant nucleotide coordinates are provided in parentheses above the chromosome (based on the rDNA reference sequence in NCBI, accession #AB026819.1). The most distal *Pst*I site is at position 5,701, which lies ∼2.05 kb from the chromosome end in LpKY97. rDNA TRF truncations were identified by targeted cloning of novel restriction fragments, and by inspection of Illumina and MinION sequence reads. Arrows show positions of telomere addition in truncated variants that were successfully cloned. B) *Duplication of an internal sequence at a chromosome end* - presumably initiated when a chromosome end is effectively de-capped (<10 TTAGGG repeats). C) *MoTeR acquisition by plain telomeres*. Regardless, of whether it occurs through recombination, BIR, or transposition, MoTeR acquisition results in the internalization of telomere repeats, creating interstitial telomeres. Note: the black shading at the MoTeR1 3’ end represents 3’ sequence shared with MoTeR2. The MoTeR1 label is inverted to illustrate that telomeric elements are always oriented with their 5’ ends nearest to the chromosome end. White circles = (TTAGGG)_1-10_ repeats; black = TTCGGG(TTTGGG)_5-8_. D) *MAGGY insertion into MoTeR sequences*. Shown is a full-length MAGGY element inserted in an inverted orientation relative to MoTeR1. This diagram shows MAGGY inserted distal to the MoTeR1 RT probe. Restriction sites in the MAGGY LTRs causes the probe to hybridize with non-telomeric (i.e., internal fragments) in *Pst*I digests. E) *Interstitial telomere breakage*. Internal TTAGGG tracts > 3 repeat units are prone to breakage, generating recombinogenic ends that can be repaired by various processes operating on double-stranded breaks. The pathway utilized likely depends on the nature of the sequences at the exposed ends. F) *Replication slippage*. Polymerase pausing and dissociation leads to skipping forward/backward in a short tandem repeat array, resulting in contraction/expansion of the array. G) *Subtelomeric tandem repeat (STR) contraction/expansion*. The characteristic feature of subtelomeric repeat contraction/expansion is preservation of the telomere junction. Array length changes can occur via unequal chromatid exchange (shown here) or gene conversion, with contractions also being possible as a result of intrachromatid recombination. H) *MoTeR truncations*. These relatively rare occurrences involve degradation of MoTeR sequences, followed by *de novo* telomere formation (likely through telomerase action). MoTeRs with a dotted line at the 5’ borders represent truncated elements (tMoTeRs), and the truncation position is noted in parentheses. I) *D-loop formation followed by BIR*. Uncapped telomere/MoTeR sequences in a multi-element array potentially can invade homologous sequences in proximal elements and initiate break-induced replication. The “+” symbol indicates that the process can occur in an iterative fashion, generating multi-element extensions. The black shading represents the 3’ border sequences common to MoTeR1 and 2, while the gray shading represents sequence unique to MoTeR2.

**Fig. 3.**
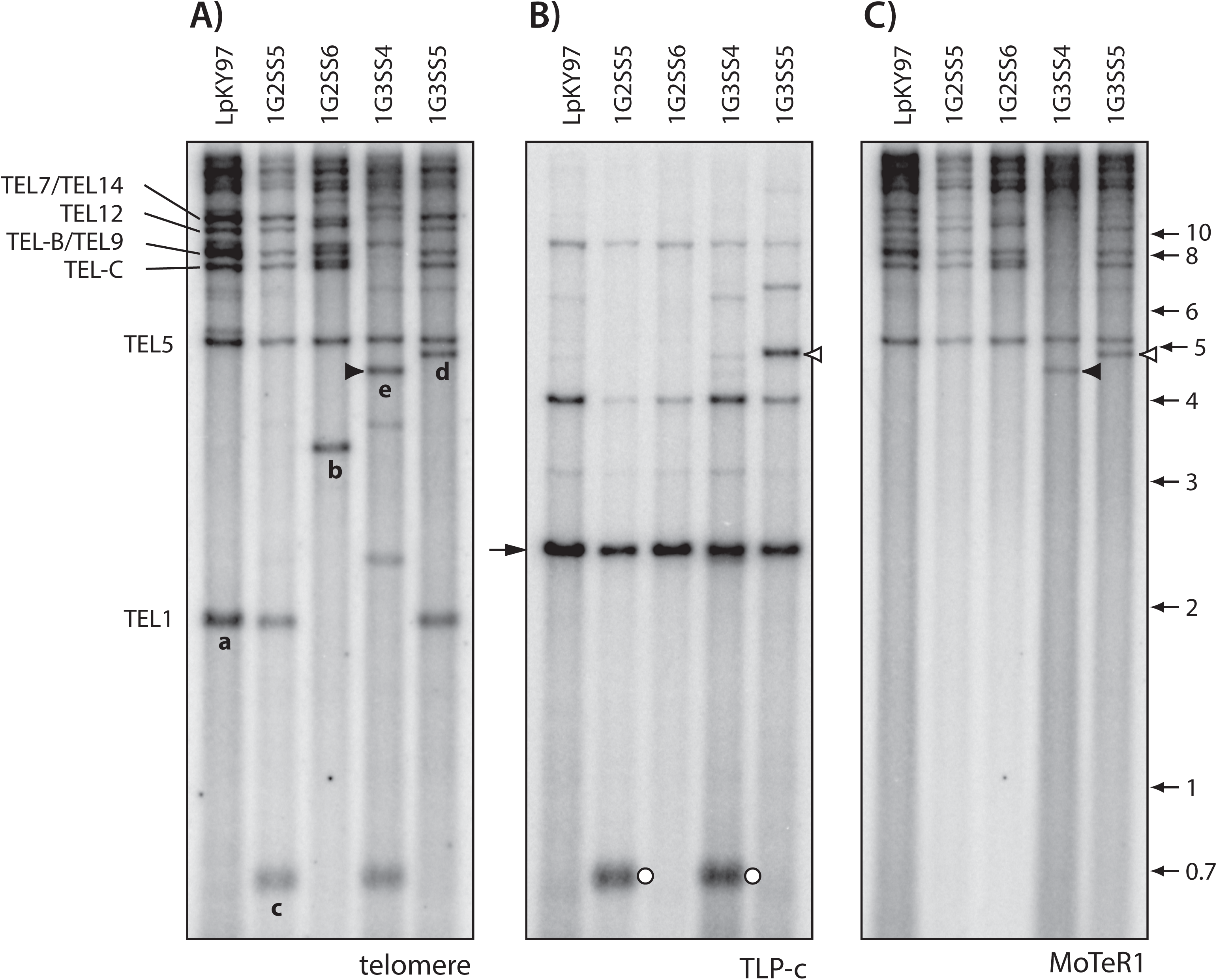
Identification of putative MoTeR1 transposition events. The three panels show selected single spore DNA samples from the population shown in Fig. 1. Panels A, B and C were hybridized with the telomere, TLP-b and MoTeR1 probes, respectively. A) Novel TRFs that have been characterized are labeled “a” through “e.” B) Re-probing of the blot with TLP-b produced a major signal in all lanes at a position corresponding to a molecular size of ∼2.5 kb. As expected, the probe hybridized to novel TRF-b, but also appeared to hybridize with novel TRF “d” (open arrowhead). C) The same blot was re-probed with the MoTeR1 probe. This also produced a hybridization signal at a position corresponding to TRF-d (open arrow), implying that TRF-d had acquired a MoTeR1 insertion. Note that the MoTeR1 probe also hybridized to TRF-e - signaling the occurrence of a second putative transposition event.

### Terminal duplication of “internal” sequences

Four single-spore progeny (1G2SS5, 1G3SS4, 1G3SS6 and 1G3SS9) exhibited a 700 bp TRF that was not present in the original parent isolate (Figs. 1 and 3A). A representative of this TRF (band “c” in Figs. 1 and 3A) was recovered from genomic DNA of strain 1G2SS5 and sequenced. To gain insight into the disposition of this sequence in the original LpKY97 parent culture, we sequenced TRF-c and generated a telomere-linked probe (“TLP-c”; note: this fragment is unrelated to TEL-C) based on the sequence adjacent to the telomere and re-probed the Southern blot shown in Fig. 3A. As expected, the probe hybridized to TRF-c in DNA from the 1G2SS5 and 1G3SS4 cultures (Fig. 3B, white circles). In LpKY97, the TLP-c probe hybridized with varying intensities to a number of fragments, with the strongest signal coming from a 2.5 kb band (Fig. 3B, black arrow), that was not telomeric (see Fig. 3A). A BLAST search of a MinION-based, chromosome level, reference genome for LpKY97 revealed that the TLP-c sequence is located ∼ 157 kb from TEL5 in the parent strain; and that the weaker signals correspond to highly diverged sequences elsewhere in the genome. The conservation of the parental band in the cultures with TRF-c, along with densitometric assessment of the parental band intensities, indicated that the novel TRF arose via duplication of the internal sequence at a chromosome end (Fig. 2B).

### Acquisition of MoTeR1 by “plain” telomeres

In culture 1G3SS5, the TLP-c probe hybridized to another telomeric fragment that was not present in the parental culture (see white arrowhead in Fig. 3B). This fragment corresponded to the novel TRF designated “d” in Figs. 1 and 3A. Considering it unlikely that a second terminal duplication of the same internal locus had arisen, we surmised that this TRF might have arisen through MoTeR transposition into the newly-formed TEL-c (see Fig. 2C). To explore this possibility, we re-hybridized the membrane sequentially with probes derived from MoTeR1 and MoTeR2. Consistent with our prediction, TRF-d co-hybridized with the MoTeR1 probe (Fig. 3C, white arrowhead). The presence of MoTeR1 within the telomere was subsequently confirmed by amplifying and sequencing across the predicted TLP-c-MoTeR1 junction, which revealed a MoTeR1 3’ terminus just 5 telomere repeat units from the telomere-adjacent sequence (Fig. S3A)). Notably, a variant (TTAGG) repeat unit in the TRF-c telomere was retained in the MoTeR1-containing version (Fig. S3Aii, sequence highlighted with double underlining). No amplification product was obtained when using DNA from the original parental culture as template, making it unlikely that this telomere-MoTeR1 junction was acquired from another chromosome location via recombination. Finally, we used a targeted cloning approach (Farman, 2011) to isolate the novel TRF in its entirety. Sequencing showed the telomere had acquired a truncated MoTeR1 (“tMoTeR1”) insertion that was missing 783 bp from its 5’ end (Fig. S3A.i).

As seen in Fig. 3C, the MoTeR1 RT probe also bound to a novel TRF in isolate 1G3SS4 (black arrowhead). Considering that MoTeR1 lacks *Pst*I restriction sites and insertion into a telomere should therefore cause the corresponding TRF size to increase, we reasoned that the novel band was likely derived from TEL1 - the only parental TRF that is smaller; and which was conspicuously absent in the 1G3SS4 culture (Figs. 1 and 3A). Furthermore, because a full-length MoTeR1 is ∼5 kb, we surmised that the insertion must have involved another truncated MoTeR1 of ∼2.5 kb in length. A PCR screen for the predicted rDNA-MoTeR1 junction in the single-spore culture yielded an amplification product whose size was consistent with the insertion of MoTeR1 in the rDNA telomere, and sequencing of the PCR amplicon revealed the MoTeR1 3’ border sequence positioned a single TTAGGG repeat unit away from the junction between the telomere and the rDNA sequences (Fig. S3B.ii). The novel TRF was subsequently isolated via targeted cloning, and end-sequencing revealed that the newly-acquired MoTeR1 insertion was truncated at nucleotide position 2252 and capped with a canonical telomere (Fig. S3B). No amplification products were obtained when using the LpKY97 parental DNA as template, which ruled out the possibility that the band arose through alteration of a MoTeR-containing rDNA telomere already present in the parent culture.

TEL5, which was relatively stable, resided on a 5.7 kb *Pst*I fragment and harbored a single full-length MoTeR1 insertion in the telomere (Fig. S3Ci). This fragment was lacking in 2G3SS4 (Fig. S2) and sequential re-probing with TLP5 and MoTeR sequences revealed that the fragment had increased in size by ∼ 1 kb (Figure S2, band “f”), and now hybridized with both the MoTeR1 and MoTeR2 probes (results not shown). Targeted cloning and sequencing of the new fragment, as well as MinION sequencing of a related SS culture (2G4SS4-20, see below), revealed two truncated MoTeR2 (tMoTeR2) insertions inserted distally to the original MoTeR1 (Figure S3C.i). Again, identically truncated elements were not found among Illumina sequencing reads acquired from the original parent, or MinION reads for two other SS isolates (see below), making it unlikely that these elements were simply acquired from other chromosome ends via recombination, or gene conversion.

### Transposition of the MAGGY LTR retrotransposon into MoTeR sequences

Shotgun cloning of telomeric *Pst*I fragments in DNA of the original starting culture and two single-spore isolates resulted in the repeated recovery of small hybridizing fragments, while recovery of larger ones was rare (data not shown). Therefore, we switched to a targeted cloning strategy to isolate novel TRFs in a more efficient manner [60]. To avoid repeated recovery of variably truncated rDNA telomeres (TEL1), we focused our cloning efforts on DNA samples where the parental TEL1 band was still present.

Among the first novel telomere clones isolated using the targeted approach were three that contained telomere repeats linked to the MoTeR 5’ terminus at one end of the insert and sequences from the long terminal repeat (LTR) of the MAGGY retrotransposon at the other (Fig. S4). MAGGY contains one *Pst*I site in each LTR and, therefore, insertion of this element into a MoTeR array (which lacks *Pst*I sites) is usually expected to cause the size of the terminal *Pst*I fragment to decrease (see Fig. 2D).

In single-spore cultures 3G1SS50 and 3G1SS76, MAGGY insertions occurred in MoTeR1 sequences, and were located at nucleotide positions 2799 and 4205; and were in sense and antisense orientations (relative to the MoTeR target), respectively (Fig. S4). In the case of 3G1SS50, the insertion position was between the telomere and the sequences used as the MoTeR1 probe, which meant that the novel TRF failed to hybridize with the MoTeR probe (data not shown), thereby obscuring the association between MoTeR presence and telomere instability. Cultures 3G1SS56 and 3G1SS145 inherited the same MAGGY insertion, which occurred in the penultimate MoTeR element, relative to the telomere. Because the integration site was at position 854 (sense orientation) in the 5’ region that is conserved between MoTeR1 and MoTeR2, the specific identity of the target MoTeR (1 vs. 2) was undetermined. Interrogation of Illumina MiSeq sequence data for strain LpKY97 identified 23 reads that spanned MAGGY-MoTeR boundaries and in each case, the insertion position was at nucleotide position 3510, which corresponds to an insertion in an extended MoTeR array in TEL-D on mini-chromosome 2 (Fig. S5 and Supplemental Results). This confirmed that the three cloned TRFs represent new MAGGY integration events.

As several of the first telomere rearrangements characterized were due to MAGGY insertions, we surmised that such occurrences, along with rDNA truncations, might predominate the alterations in MoTeR-containing telomeres. Considering that our goal was to gain a comprehensive insight into mechanisms underlying MoTeR-associated instability, we devised a Southern hybridization-based strategy to pre-screen for MAGGY insertion events. The genomic DNA samples were digested (separately) with *Afl*II and *Sac*II and then probed with internal MAGGY sequences, mA and mS (see Fig. 4A). *Afl*II and *Sac*II cut within MAGGY but not in MoTeR1 or MoTeR2 so that digestion should release a TRF that co-hybridizes with either mA or mS depending on the orientation of the MAGGY insertion. When MAGGY is inserted in a MoTeR array, it will produce predictable size shifts when comparing *Afl*II and *Sac*II restriction digests, with the shifts corresponding to the distance between the *Afl*II and *Sac*II sites: ∼ 1.2 kb for MAGGY insertions in the sense orientation, and ∼1.4 kb for antisense (Fig. 4A). The orientation is then easily established based on which probe (mA or mS) hybridizes with the TRF.

**Fig. 4.**
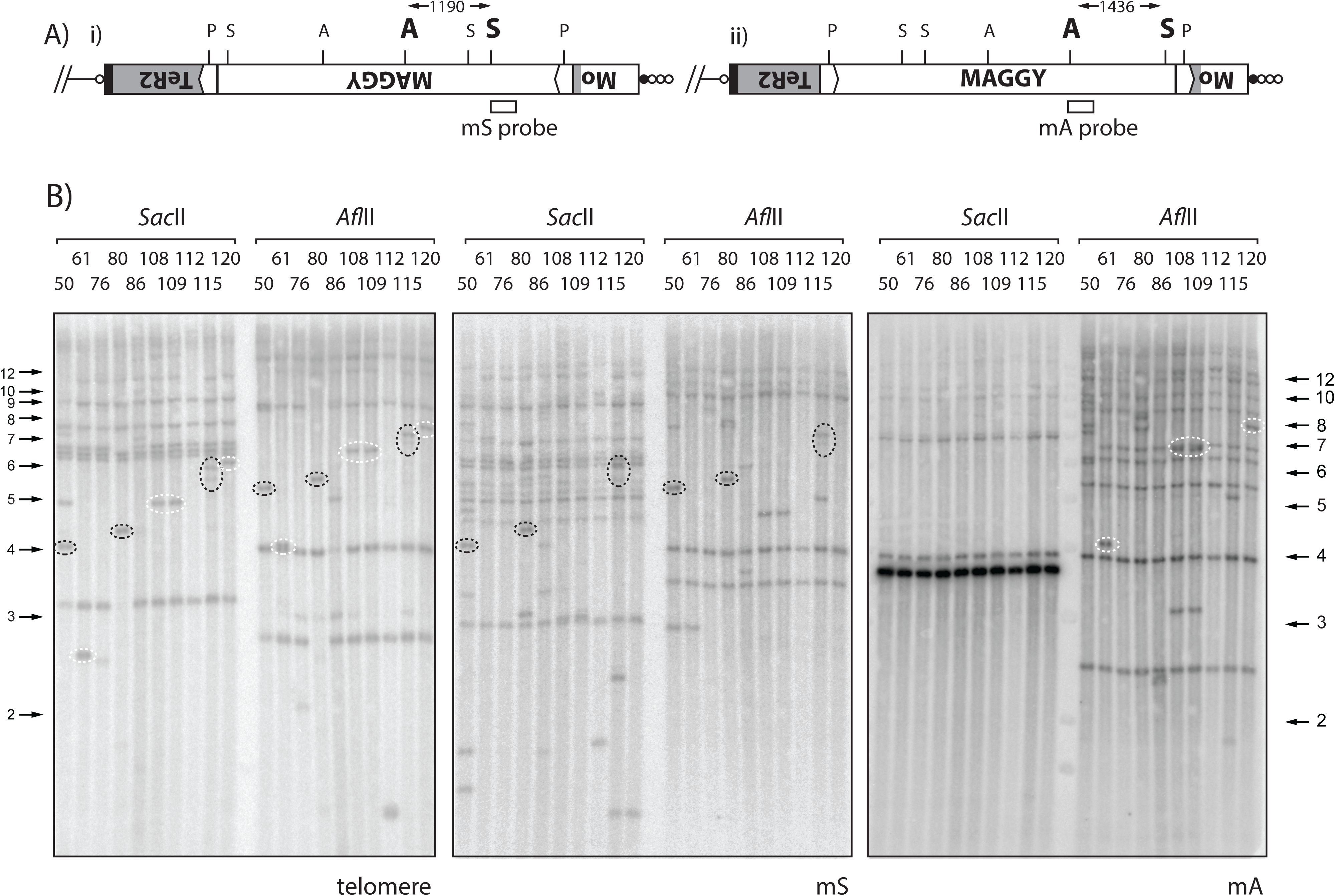
Detection of new MAGGY insertions. A) MAGGY can insert into MoTeR sequences in two orientations relative to the telomere: i) sense and ii) antisense. The gray shaded boxes represents a MoTeR1 element that has been disrupted by the MAGGY insertion. Hybridization probes were developed to detect each orientation: mS (sense; *Sac*II digest) and mA (antisense; *Afl*II digest). B). Detection of new MAGGY insertions. Genomic DNAs from the single spore isolates were digested with *Sac*II or *Afl*II, size-fractionated by electrophoresis and probed sequentially with the telomere, mS and mA probes. The resulting phosphorimages are shown. Novel telomeric fragments that contain MAGGY sequences are highlighted. Left-hand panel: telomere probe; middle panel: mS probe; and right-hand panel: mA probe. With the telomere probe, note that certain TRFs have predictable size shifts between the *Afl*II and, *Sac*II digests, which represents the distance between the *Sac*II and *Afl*II sites in MAGGY. The orientation of each MAGGY insertion is easily determined based on whether the mS probe (dotted black ovals) or the mA probe (dotted white ovals) co-hybridized with the telomeric fragment.

Examples of MAGGY screening results are shown in Fig. 4B. Among the nine single-spore cultures shown in the figure, the telomere probe identified at least 11 novel TRFs migrating at positions where the hybridization signals were clearly visible (see Fig. 4B, left-hand panel, *Sac*II digest). Eight exhibited 1.2 or 1.4 kb size increases in the *Afl*II digest, consistent with the restriction site having been contributed via integration of MAGGY near the telomere. The existence of these insertions was confirmed by sequential probing with the mS and mA sequences: four of the novel TRFs hybridized with the mS probe (black dotted rings), indicating four, independent MAGGY insertions orientated away from the telomere (i.e. in the direction shown in Fig. 4A(i)). The remaining four novel TRFs hybridized with mA (white dotted rings), consistent with three independent insertions oriented toward the telomere (ii) (note that cultures 108 and 109 appear to be clones of one another). Two additional *de novo* MAGGY integrations were identified among MinION reads from fourth generation SS isolates. Isolate 2G4SS6-1 (hereafter, SS6-1) had a MAGGY insertion in a MoTeR1 element that formed part of an array in TEL-A (Fig. S6A), while 2G4SS15-11 (hereafter, SS15-11) had a MAGGY inserted in a MoTeR1 in TEL11. Interestingly, this latter element already contained an insertion of a copia-like retroelement, RETRO8 (Fig. S6B)

Based on the sizes of new *Pst*I TRFs, we were able to estimate the insertion positions for the remaining MAGGY insertions (Table S3) which revealed that none of them corresponded to the “MoTeR-inserted” MAGGY in the starting culture. This confirmed that all of the novel TRFs arose through new MAGGY insertion events, as opposed to recombination with the existing MAGGY-MoTeR array. Altogether, we identified twelve *de novo* MAGGY insertion events among 34 single-spore isolates that were screened and/or sequenced. Also, note that with the exception of 3G1SS76, all of the insertion positions were 5’ to the MoTeR1 RT and MoTeR2 probe regions, which explained why MoTeR probes often failed to hybridize to novel TRFs (data not shown).

### Break-induced expansion and contraction of MoTeR arrays

Shotgun cloning of TRFs from single-spore culture 1G2SS2 resulted in the recovery of an 8.7 kb fragment which contains TEL14. Restriction analysis and strategic sequencing revealed the telomere contains two MoTeR insertions, one a severely truncated MoTeR1 in the proximal position, and an intact MoTeR2 inserted distally (Fig. 5A). All single-spore cultures, except for 2G3SS12, 13, 17 and 18, showed hybridization at the expected position (Fig. 1, Fig. S2 and Fig. 5B). However, when compared with the TEL5 bands, the varying signal strengths for TEL14 suggested that some cultures might possess additional TRFs with a similar size. Therefore, to monitor TEL14 instability more accurately, we used the telomere-linked probe, TLP14, to re-probe the telomere blot. In addition to expected signals from bands of 8.7 kb (band “g”) that corresponded to the cloned TRF, three additional signals were detected (bands “h”, “i”, “j”) among the other SS isolates, with the molecular sizes of the bands differing by increments of approximately 1.7 kb - the length of MoTeR2 (Fig. 5B). Accordingly, we surmised that TEL14 is unusually unstable and undergoes repeated rearrangements that involve simple additions and losses of MoTeR2 copies. This supposition was confirmed through targeted cloning and sequencing of TEL14-h, -i, and -j variants from representative 3rd generation SS cultures, and by inspecting MinION reads for representative 4th generation SS cultures (see below). The deduced MoTeR arrays structures are shown in Figure 5C.

**Fig. 5.**
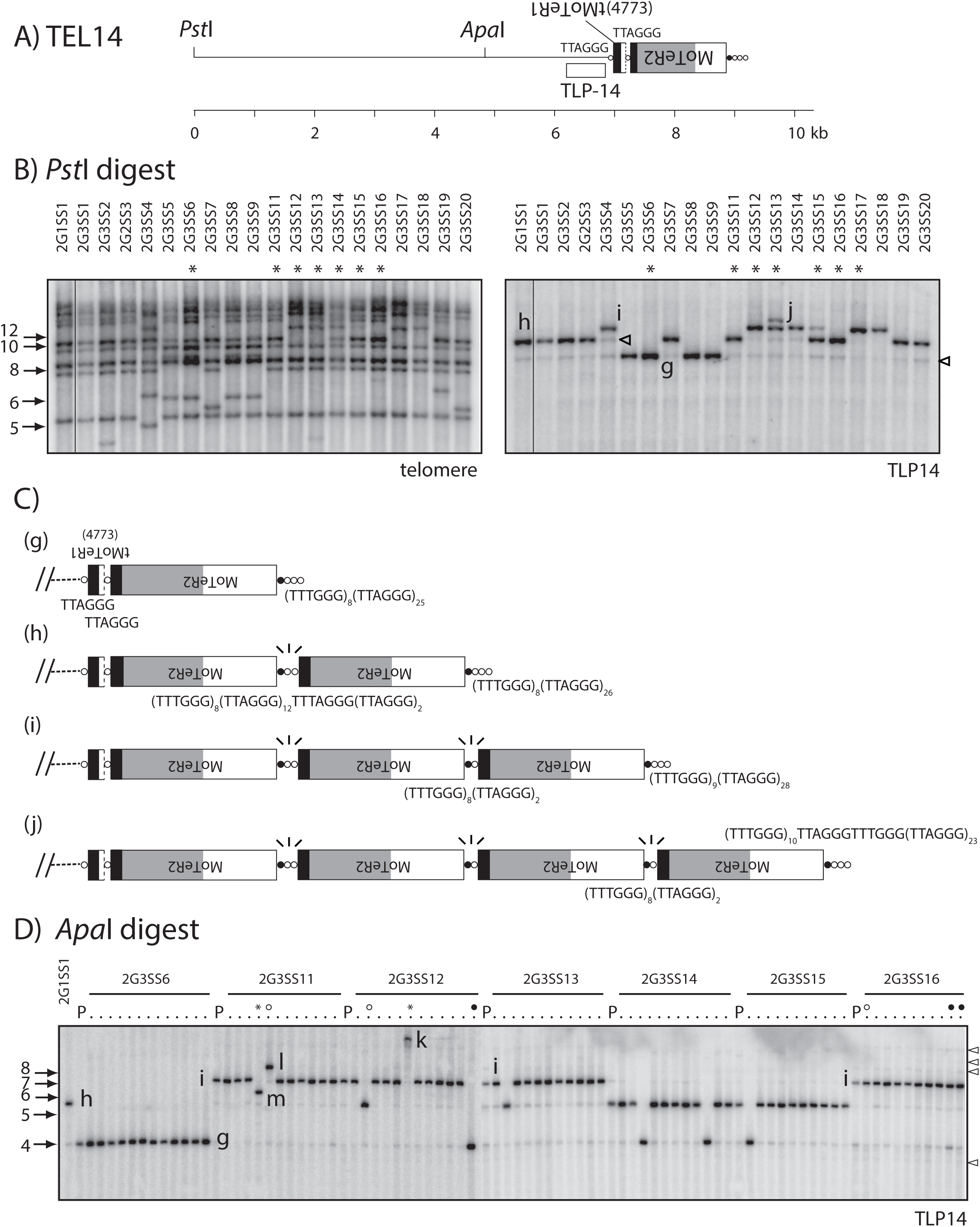
Recurrent rearrangements of TEL14 promoted by MoTeR-generated, interstitial telomeres. A) Organization of the TRF containing TEL14. MoTeR2 segments are shaded to illustrate relevant portions: white = 5’ sequence shared with MoTeR1; light gray = sequence unique to MoTeR2; black = 3’ sequence shared by both elements. The TLP14 probe was derived from chromosome-unique sequence immediately adjacent to the tMoTeR1 insertion. Telomeric circles are not drawn to scale so the ruler is approximate. B) TLP14 identified multiple forms of the TRF in single-spore progeny. The left-hand panel shows a segment from a telomere-probed blot of single-spore cultures derived after two successive rounds of plant infection (Starnes et al. 2012). The blot was stripped and hybridized with the TLP14 probe. The different TRF forms are labeled g, h, i and j. Cultures with the longer TRF variants were invariably heterokaryotic and contained some nuclei with shorter forms of the telomere (closed arrowheads). Cultures used to generate the fourth generation single-spore families shown in C are labeled with asterisks. C) Structural maps of TRFs g, h, i and j. Sequences of interstitial telomeres and telomere-like motifs are positioned below the relevant MoTeR-MoTeR junctions. Junctions without labels have the same sequence as the equivalent, labeled junction above. The fragile MoTeR2-MoTeR2 junctions experiencing breaks are indicated with 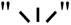 motifs. Band-j is denoted with an asterisk to indicate that its structure is inferred from MinION reads of another SS isolate (2G4SS4-20 and not 2G3SS12). D) TRF14 profiles for 4^th^ generation single-spore families (10 to 12 spores/generation) isolated from the 3rd generation SS cultures (see Fig. S1). DNA samples from the G4 cultures, their respective parents, and the original parent were digested with *Apa*I, fractionated by agarose gel electrophoresis, blotted to membranes and hybridized with the TLP14 probe. The image shows two separate blots that have been placed side-by-side. In the left-most lane of each blot is DNA from strain 2G1SS1. Then, for each family, the left-hand lane has DNA from the 3^rd^ generation parent (labeled “P”, with its identifier provided above the respective block) and DNA from the corresponding single-spore progeny to the right. TRF variants are labeled “g” through “m.” Lanes marked with open and closed circles, and asterisks are discussed in the text.

Many single-spore isolates yielded multiple signals with the TLP14 probe, with most producing one strong signal and a number of fainter ones. In almost every case, the faint signals came from fragments whose sizes matched those of fragments showing strong hybridization in other samples (see Fig. 5B, right-hand panel). That is to say, the cultures with the longer “h,” “i” and “j” TRFs invariably had weak hybridization signals at molecular sizes corresponding to one or more of the shorter forms. To explain this pattern, we surmised that the fungal mycelium was heterokaryotic for variant TEL14 forms because longer MoTeR2 arrays were experiencing frequent breakage at the interstitial telomere sequences that are usually found at MoTeR-MoTeR junctions. Furthermore, based on signal intensities, we estimated that approximately 10 % of nuclei contained a truncated MoTeR array. To test this idea, we generated fourth generation SS families from seven of the 2G3SS isolates (# 6, 11, 12, 13, 14, 15 and 16) and examined their TLP14 restriction profiles. If the faint hybridization signals come from a fraction of nuclei carrying truncated TEL14 MoTeR arrays, these nuclei should segregate during sporogenesis, resulting in SS cultures in which the formerly “minor” variant is now the dominant form. Moreover, the proportion of fourth generation SS cultures containing these shorter variants should approximate the estimated proportion of variant TRFs within the presumed heterokaryotic mycelium (∼10%). For these experiments, the DNA samples were digested with *Apa*I which does not cut in MoTeR2 (see Fig. 5A) and generates a smaller TRF that is better resolved during electrophoresis. The blotted DNAs were hybridized with the TLP14 probe, which revealed that five of the seven 4th generation spore families contained at least one culture in which a shorter MoTeR array was now the dominant form. For example, starting cultures 2G3SS14 and 2G3SS15 contained variant “h” as the major fragment but also exhibited weaker hybridization to a fragment migrating at the “g” variant position (Fig. 5D). Two of the SS cultures from 2G3SS14 exhibited hybridization exclusively at the position of variant “g”, while the other eight cultures had a TEL14 profile similar to that of their parent. Likewise, the spore family from 2G3SS15 had one individual with variant “g” and nine with the parental pattern (“h”). Again, a majority of the cultures having band “h” as the predominant form also showed weak hybridization at a position corresponding to “g.”

The 2G3SS12 culture had the majority variant, “i,” and also showed faint hybridization at positions corresponding to “g” and “h.” Consistent with these faint signals, one of the ten 4th generation SS cultures lacked hybridization to “i” and contained variant “h” as the predominant form instead (open circle). Another culture showed exclusive hybridization to variant “g” (closed circled). Interestingly, a third SS isolate from 2G3SS12 (asterisked) exhibited hybridization to very high molecular weight band (“k”) that was estimated to be much more than 10 kb longer than fragment “j” (Fig. 5D). Here, it is important to note that MoTeR1 has an *Apa*I site, so these large additions are unlikely to represent MoTeR1 acquisitions.

One of the SS cultures from 2G3SS11 yielded a new TRF (band “l”) that was ∼1.7 kb greater than the “i” variant (open circle). This likely resulted from the addition of a fourth MoTeR2 copy to the i-type array. The 2G3SS11 SS family also contained an individual, in which TEL14 was contained on a fragment (“m”) that was intermediate in size between “h” and “i” (asterisked, Fig. 5D). Based on the retained hybridization to the telomere probe (data not shown), this fragment likely consists of an i-type array in which the distal MoTeR2 copy experienced truncation, followed by *de novo* telomere addition. Two of the original cultures did not segregate variants in the single-spore families.

As was the case with the 2G3SS14 and 2G3SS15 spore families, several of the SS cultures from 2G3SS11, 2G3SS12 and 2G3SS16 showed faint hybridization signals above the “i” band, with estimated size increments of 1.7, 3.4 and 5.1 kb (Fig. 5D, open arrowheads on right-hand side). We suspect these come from TRFs with additional insertions of one, two and three MoTeR2 copies, respectively. Critically, none of the SS cultures having band “g” as the predominant form showed hybridization to any *Apa*I fragments that are larger than the “g” band. Although, some samples do have possible weak signals below (bottom arrowhead, Fig. 5D).

Further evidence that the faint signals are due to heterokaryosis for variously truncated MoTeR arrays came from the observation that when “i”, “h” and/or “g” fragments co-occurred, their relative signal intensities varied. Clear examples of this are seen in the spore cultures from 2G3SS16, which showed strong hybridization to “i,” clear signals in the “g” position but little or no hybridization to “h” (Fig. 5D, “parental” and open circle lanes). Likewise, cultures showing similar hybridization intensities for the major TRF often had varying signal strengths to the minor bands (e.g. the rightmost progeny from 2G3SS16 – closed circles).

Work in other systems has shown that interstitial telomeres undergo frequent, spontaneous breakage [67–72] and, in fact, we and others have utilized this property as a facile way to engineer breaks in chromosomes [67, 73]. Because the MoTeR arrays were truncated precisely at MoTeR-MoTeR junctions, we reasoned that the interstitial repeats between elements promote breakage (Fig. 2E) and then serve as seeds for end-repair via telomerase-mediated telomere addition (“telomere healing”). Furthermore, we predicted that the rearrangement-prone “h,” “i” and “j” variants of TEL14 contain elements separated by multiple TTAGGG and TTAGGG-like motifs. This would be in contrast to the “g” form which has single TTAGGG motifs separating elements, and which is not expected to be unstable - considering the existence of ∼7,000 instances of TTAGGG/CCCTAA genome-wide. To test this prediction, we sub-cloned a number of TEL14 variants and sequenced the MoTeR-MoTeR junctions. In addition, we inspected MinION reads from three SS cultures. This confirmed that all junctions experiencing breakage comprised long, interstitial, telomere-like/telomere tracts - the shortest examples being TTCGGG(TTTGGG)_8_(TTAGGG)_2_ and the longest, (TTTGGG)_5_TTAGGGTTTGGG(TTAGGG)_18_ (e.g., Figs. 5C.h & S12.Bi). Critically, the MinION reads for TEL14 revealed variable (TTTGGG)n tract lengths between interstitial telomeres that occupied equivalent positions in TRFs with the same overall MoTeR array configuration (Fig. S7). This is most easily explained by replication slippage (Figure 2F) - a known mechanism underlying length variation in tandem repeat sequences [74]. Given, the highly dynamic nature of TEL14, however, it is also possible that the observed length polymorphisms could have arisen through iterative cycles of break-induced replication (BIR). One apparent example of this is shown in Fig. S7 (variant iii), where it appears that an interstitial break between elements 2 and 3 in variant i or ii was repaired by invading one of the more distal interstitial telomeres.

Telomeres 6, 12, 14, A, B, and D also underwent rearrangements that also appeared to have been caused by simple breaks within interstitial telomeres (see Supplemental results). On the other hand, some telomeres exhibited alterations that resembled simple interstitial breaks, but closer inspection of the variant telomeres sequences subtending the new telomeres (and sometimes separating other elements in the array) pointed to the occurrence of more complex repair mechanisms, including D-loop formation (e.g. TEL3, see below), or invasion of sister chromatids/other telomeres (Supplemental results).

### “Contraction/expansion” of subterminal tandem repeats

Single-spore cultures 2G2SS12, 2G2SS13, 2G2SS17 and 2G2SS18 exhibited a novel TRF at a molecular size corresponding to ∼ 3.7 kb (band “n”, Fig. S2 & Fig. 6A). The TRF in the 2G2SS17 culture was cloned and subjected to end-sequencing, which revealed a simple telomere at one end of the insert (Fig. 6B, SS17). Comparison with the B71 reference genome identified this as TEL4. To determine the molecular events giving rise to the novel TRF, telomere-linked probe, TLP4 (Fig. 6B), was used to re-probe the stripped blot. As expected, TLP4 hybridized with the 3.7 kb fragments but produced very faint, high molecular weight bands with smeared tails in the lanes containing DNAs from the other SS cultures (Fig. 6A, right-hand panel). We suspected that the parental TRF was large and was sheared during DNA isolation, causing the hybridization signal to be distributed down the entire lane. This was confirmed using spot blots, which showed that all DNA samples hybridized to TLP4 with approximately equal intensities (Fig. S8A). Next, we generated agarose-embedded, intact chromosome preparations [15, 75] which were then digested with *Pst*I, fractionated using contour-clamped homogeneous electric field (CHEF) electrophoresis, blotted to a membrane, and sequentially hybridized with the TLP4, telomere and MoTeR probes. TLP4 hybridized to large fragments that were more than 45 kb in length (Fig. S8B) and whose sizes appeared to vary slightly among different chromosome ends. Additionally, cultures 2G1SS1 and 2G3SS1 exhibited a number of faint signals to shorter fragments (indicated with arrowheads). Interestingly, in cultures 2G1SS1, 2G3SS14 and 2G3SS20, the TLP4-hybridizing fragments did not hybridize with the telomere probe and are, therefore, not telomeric (Fig. S8B). They also failed to hybridize with either MoTeR probe.

**Fig. 6.**
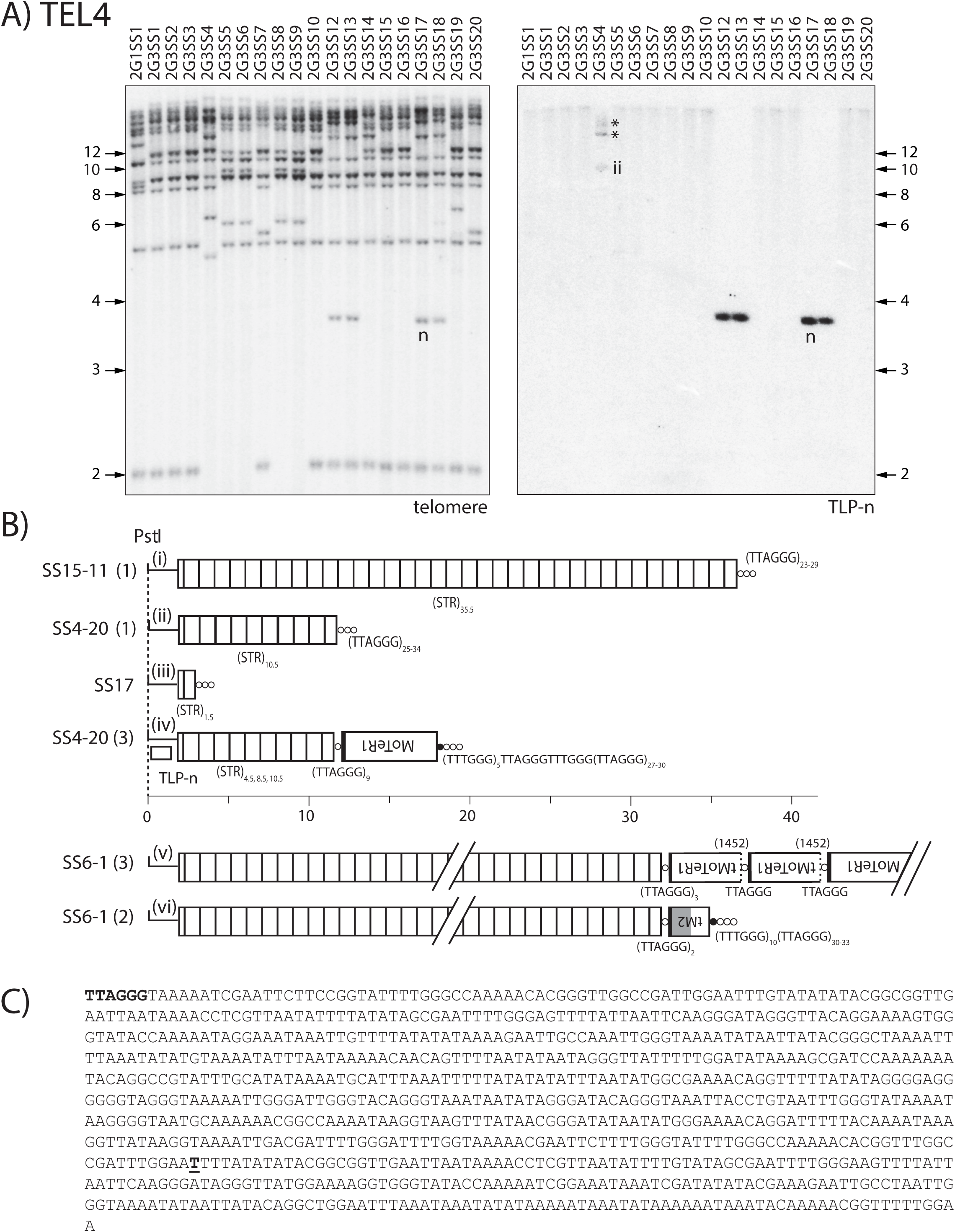
Subtelomeric repeat (STR) expansions/contractions, interstitial telomere breaks and MoTeR acquisitions at TEL4. A) Southern hybridization analysis of SS progeny showing novel TRF-n (left-hand panel) and monitoring instability at the corresponding telomere (TEL4) using TLP-n (right-hand panel). B) TRF structures elucidated via MinION sequencing of SS4-20, 6-1 and 15-11 and Sanger sequencing of TRF-m. MoTeR arrays are annotated according to the standard schema. Individual units of a tandem repeat array found in the subterminal region of TEL4 are represented as open boxes. MinION sequencing identified distinct TRF structural variants in SS4-20 and SS6-1, while SS15-11 contained a single TRF form. The TRF structure for SS17 was inferred from Sanger sequence data. All structures shown as fully contiguous entities were inferred from individual MinION reads that spanned the entire distance from the chromosome unique sequence to the telomere. Inferred rearrangement mechanisms: i -> ii -> iii: intrachromatid recombination, unequal sister chromatid exchange or gene conversion; ii -> v) MoTeR acquisition via ectopic recombination with TEL8; array shortening via intrachromatid recombination, unequal sister chromatid exchange or gene conversion. C) Nucleotide sequence of an individual tandem repeat unit. The single CCCTAA repeat unit is shown in bold and the position at which the telomere starts in all of the SS cultures is marked with bold, underlined text.

Inspection of MinION reads for the fourth generation SS cultures SS6-1 and SS15-11 indicated that the TEL4 *Pst*I fragments were telomeric in these strains, and contained many copies of a ∼0.9 kb subtelomeric tandem repeat (STR) between the *Pst*I site and the telomere (Fig. 6B and Fig. 6C). For SS15-11, the complete *Pst*I TRF was captured in a single MinION read, which contained 35.5 copies of the STR which was capped with a canonical telomere (Fig. 6B). Southern blot hybridization suggested that the TRF was mostly stable in this particular strain, although a faint signal was observed at a molecular size of ∼17 kb (Fig. S8C).

MinION reads from 2G4SS4-20 (hereafter, SS4-20) showed an identical STR-telomere junction but the subtelomeric array contained varying numbers of repeat units ranging from 4.5 to 10.5 repeat units (variants ii & iv). In addition, a number of reads from this strain had a MoTeR1 inserted in the telomere, with varying STR lengths (Fig. 6B.iv). These reads likely correspond to the various bands seen in 2G3SS4 lane in the Southern blot (Fig. 6, asterisks). Although long, STR-containing reads of up to 34.9 kb, with up to 22.5 repeat units, were found for SS6-1, none of them spanned the entire STR array. Southern blot hybridization using TLP4 with *Pst*I-digested DNA from SS6-1 produced an indistinct smear (Fig. S8C), which showed that TRF4 experienced recurrent rearrangement despite the recent genetic purification of this strain via single-spore isolation. The original 3.7 kb TEL4 restriction fragment from SS17 contained just 1.5 repeats units (Fig. 6B). An overwhelming majority of the MinION reads containing TEL4 exhibited identical boundaries between the telomere repeats and adjacent sequence. From this, we conclude that most TRF4 rearrangements were unrelated to the telomere *per se* and, instead, involved simple contractions/expansions of the STR array - likely via unequal intra-chromatid and/or unequal sister chromatid exchanges (Fig. 2G).

Interestingly, the same STRs were also present in the TEL8 subtelomere (Fig. S9), and the corresponding TRF experienced identical alterations to TEL4 - namely subtelomeric array contraction/expansions, as well as MoTeR1 acquisition by a plain telomere (Fig S9A.ii). Intriguingly, in both cases, the interstitial telomeres that separated the MoTeR element from the telomere-adjacent sequence were nine TTAGGG repeat units in length (see Figs. 6B.iv and S9A.ii). This provides a strong indication that the MoTeR sequence was exchanged between the two chromosome ends, with the most likely route being unequal, inter-chromosomal exchange, as this would also explain the reciprocal extensions and contractions.

Another example of a tandem repeat array experiencing contraction and expansion, in the absence of terminal alterations, involved a MoTeR array in TEL10 (Supplemental results).

### Simple terminal truncations

Another type of alteration in MoTeR-containing telomeres was identified in a separate study involving analysis of progeny from a cross between a MoTeR-lacking strain of *M. oryzae*, Guy11, and strain 2539, which has three MoTeR-containing telomeres. 2539 has a single, intact MoTeR2 copy inserted distal to a truncated MoTeR1 in TEL10 (Fig. 7A). This telomere is relatively stable and in most cultures has remained unaltered through repeated subculturing over several years. In addition, in most cases, it is faithfully transmitted through meiosis (Fig. 7B lower panel; and data not shown). Southern hybridization analysis using MoTeR2 to probe *Pst*I-digested DNAs from 52 progeny of a cross between 2539 and Guy11 revealed a single alteration among 25 progeny that inherited this telomere (representative isolates are shown Fig. 7B). Progeny isolate 6077 contained two MoTeR2-hybridizing bands, with the second one hybridizing at a slightly weaker intensity. Cloning of the new TRF from strain 6077 revealed that it is identical in structure to TRF10, except that the MoTeR2 had suffered a terminal truncation (Fig. 2H) that eliminated 856 nucleotides from the 5’ end (Fig. 7C). The truncation happened within a region that was highly rich in thymine residues (57 out of 85 nucleotides, 67%), with the new telomere having been extended off a “T” nucleotide seed (underlined) (Fig. 7D).

**Fig. 7.**
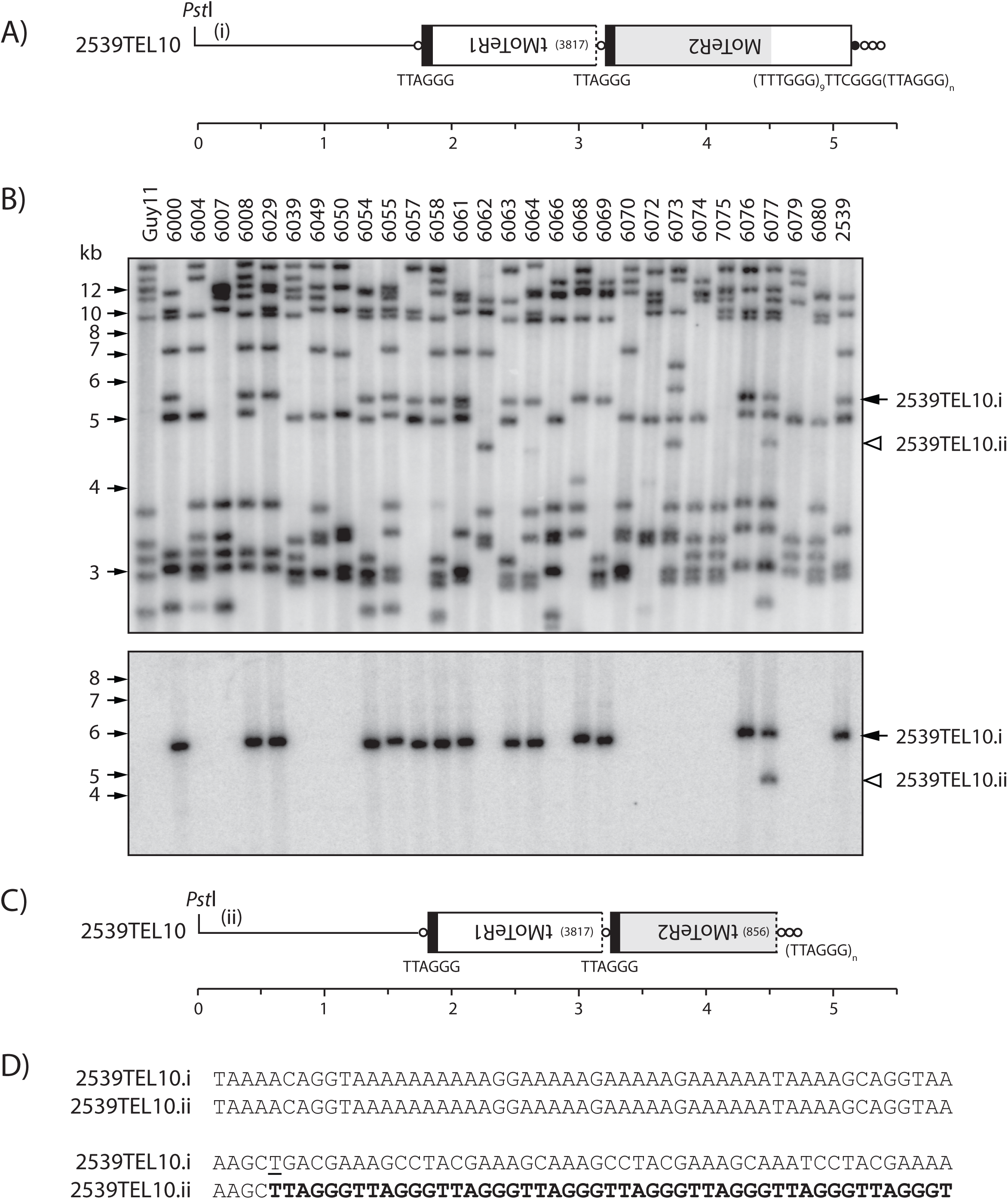
Simple terminal truncations. A) Telomeric *Pst*I fragment containing 2539TEL10 showing MoTeR sequences embedded in the telomere. B) TRF segregation among progeny of a cross between 2539 (MoTeR^+^) and Guy11 (MoTeR^-^). The top panel shows the phosphorimage of a Southern hybridization analysis of *Pst*I-digested DNAs probed with telomere repeats. Parental strain DNAs are loaded in the outermost lanes, with progeny DNAs in between. 2539TEL10.i is marked with a black arrowhead. A white arrowhead marks a novel TRF that corresponds to a TEL10.ii variant. Bottom panel: the membrane used in A was stripped and hybridized with the MoTeR2 probe. C) Organization of 2539TEL10.ii showing 5’-truncation of the terminal MoTeR2. D) 106 bp of sequence surrounding the site of *de novo* telomere addition at the MoTeR2 truncation boundary. Telomere repeats are highlighted with bold text. A “T” nucleotide that seeded the *de novo* telomere addition is underlined. inferred rearrangement mechanism: i -> ii) MoTeR truncation, *de novo* telomere formation.

Another instance of terminal truncation, this time at TEL14, was detected in one of the 4th generation SS cultures (SS11-4) (Fig. 5D). We did not characterize this novel TRF, but, based on the progenitor telomere organization, and the Southern hybridization patterns, we predict that the terminal MoTeR2 copy was truncated by ∼1 kb, with the free 5’ end having been subsequently capped by *de novo* telomere addition. Finally, a third rearrangement consistent with a MoTeR1 truncation was detected at TEL2 - in this case possibly following an interstitial telomere break (Supplemental results).

### D-loop formation and tandem amplification

All of the MinION reads containing TEL3 contained an intact MoTeR1 inserted at a proximal position in the telomere (Fig. 8). In SS4-20, there were between one and five intact MoTeR2 distal copies. Structurally, this organization resembled that of TEL14. However, unlike the latter telomere, and virtually all other multi-MoTeR arrays, the lengths of interstitial telomeres and variant repeats separating element copies were identical, making it unlikely that array extensions occur via sequential transposition, or through recurrent interstitial telomere breakage. Instead, it appears that these variants represent the healed products from sequential cycles of array extension occurring via BIR with D-loop formation (Fig. 2I and Fig. 8B). Note that free ends comprising telomere repeats are expected to invade interstitial repeats at random positions and, thus, should result in alterations of interstitial repeat compositions as the array is extended. For this reason, we propose that the free end comprises internal MoTeR sequence that is shared betwen MoTeR1 and MoTeR2 (Fig. 8B) as this would constrain the invasion position and guarantee that all newly synthesized elements are identical.

**Fig. 8.**
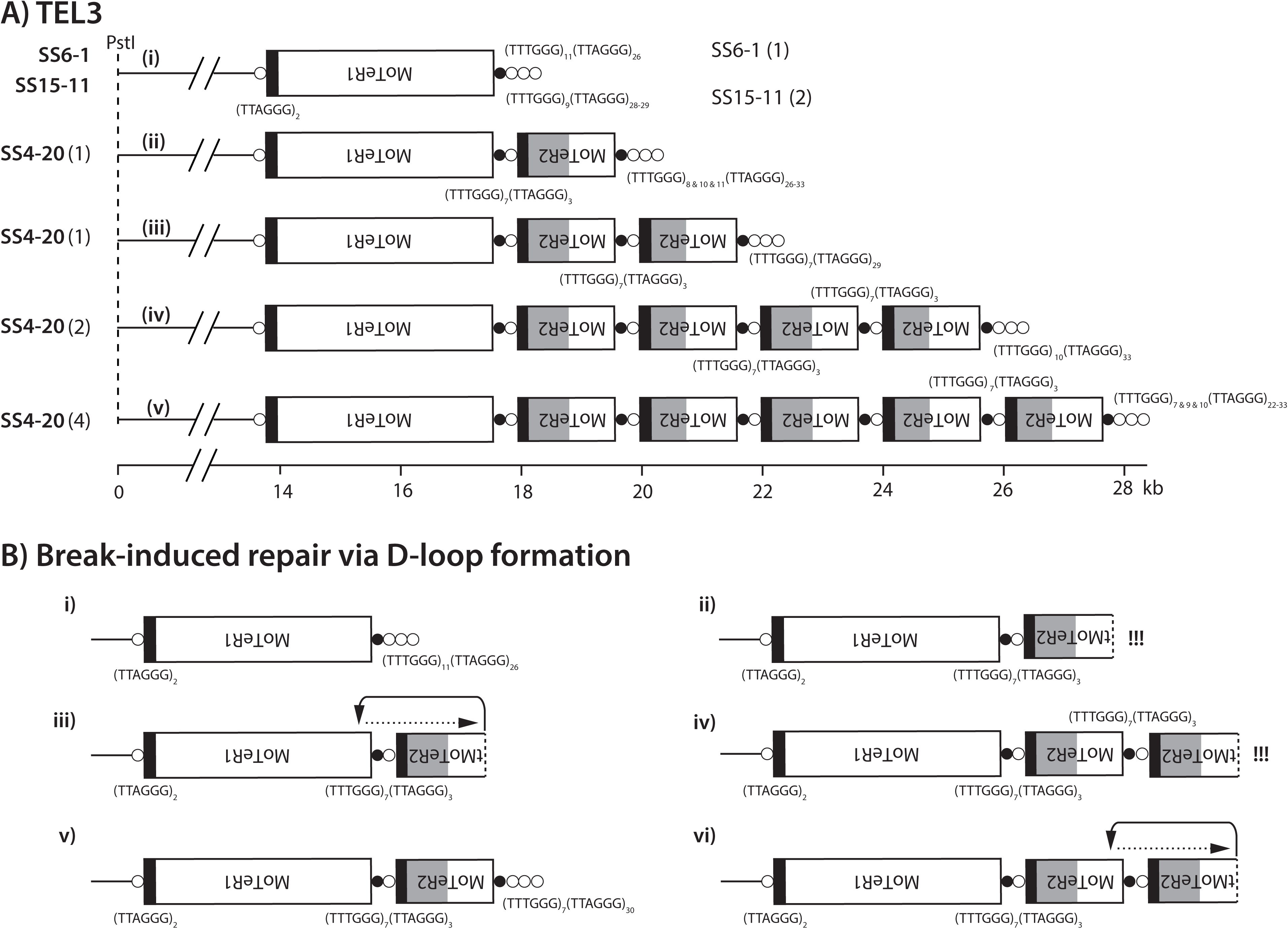
D-loop mediated MoTeR array expansions in TEL3. A) Structures of TEL3 variants identified among MinION reads. All structures were inferred from individual MinION reads that spanned the entire distance from the chromosome unique sequence to the telomere. Numbers of reads supporting each structure are indicated in parentheses. The inferred rearrangement mechanisms giving rise to the various structures are as follows: i -> ii) MoTeR2 transposition (or interstitial telomere break) followed by telomere healing; ii -> iii -> iv -> v) BIR involving D-loop formation by free end; followed by strand extension (see B). B) Schematic showing MoTeR array extensions via break-induced replication with D-loop formation. i) MoTeR2 transposition (or an interstitial telomere break at MoTeR2 proximal junction) generates a free, variant telomere-terminated DNA end (ii); iii) free end invades interstitial telomere; DNA synthesis replicates MoTeR2 sequence (iv); end of newly synthesized strand is either healed with a telomere (v), or initiates a second D-loop (vi); etc.

Further evidence that D-loop formation and extension operates on telomeric MoTeR arrays comes from the organization of an array comprising truncated MoTeR1 and MoTeR2 copies in TEL-D. This telomere contains 18 different MoTeR copies and exhibits a repeating tMoTeR1-TMoTeR2-tMoTeR2 pattern, in which MoTeR truncation positions, and the interstitial telomeres defining the tMoTeR-tMoTeR junctions, are identical throughout the array (Fig. S5). Again, precise preservation of the repeating tMoTeR1-TMoTeR2-tMoTeR2 pattern would require that the invading free end comprise internal MoTeR sequence and, more specifically, MoTeR1 sequence that is not shared with the truncated MoTeR2 copies. The variation in repeating pattern for MoTeR2 in the proximal portion of the array could be explained by intra-/unequal sister chromatid exchange.

### Stable telomeres

In addition to TELs 5 and 9, two additional telomeres (TEL11 and TEL13) were relatively stable and rarely appeared to undergo rearrangement, as evidenced by inspection of Southern blot data and MinION reads. TEL9 was visible in the Southern blots (Figs. 1 and S2) and was missing in only four out of 143 SS isolates (see Fig. 1, 1G3SS1 & 1G3SS4 as examples). In the case of TEL11, one of 37 MinION reads identified a MAGGY insertion in a MoTeR element (Fig. S6B) but, otherwise, there no rearrangements that were directly attributable to MoTeR sequences *per se*. The general organization of these “stable” chromosome ends comprised a telomere with a single intact, or truncated, MoTeR1; the interstitial telomere tract contained no more than two repeats; and the canonical telomeres were attached to truncated MoTeR sequences, or TTTGGG tracts of various lengths (Fig. S9B). Together these data suggest that the TTCGGG(TTTGGG)_n_ tracts at the 5’ ends of intact MoTeR insertions do not compromise telomere integrity by themselves, and that two consecutive TTAGGG repeats are insufficient to promote breaks. Although the general structure of the “stable” telomeres remained largely altered, varying numbers of TTTGGG repeats were sometimes found at the MoTeR 5’ termini (see Fig. 6B). These changes are not easily explained through recurrent cycles of break/repair as this ought to result in collateral alteration of the adjacent, and rather unusual, TTAGGGTTTGGG motif. Instead, we propose that alteration in the short tandem repeat tracts occur via replication slippage [74].

### The LpKY97 genome has a history of MoTeR-associated chromosome rearrangements

Many *M. oryzae* strains lack full-length MoTeR copies but, when intact elements are present, they are only found embedded in telomeres (Starnes et al. 2012; this study). On the other hand, all strains possess short relics of MoTeR sequences residing at internal chromosome locations. These relics comprise MoTeR 3’ termini attached to a short stretch of telomere sequence (Table S4) and, as such, they serve as tags for sequences that were once telomeric (Starnes et al. 2012; Farman et al. 2014). In turn, they can provide insights into MoTeR impacts on genome structure. To this end, we examined the chromosomal neighborhoods of 19 internal MoTeR relics present in the LpKY97 genome (Fig. 9A). Among these loci, 12 had 5’ flanking sequences that were duplicated, with the duplications starting at the MoTeR 5’ boundary and extended into neighboring DNA (see Fig. 9B). Nine of these structures are consistent with a scenario in which telomeric MoTeRs were resected and underwent non-homologous end-joining (NHEJ) with internal sequences that became duplicated in the process. The remaining three formed a tandem repeat in the middle of mini-chromosome 1, where the relics were interspersed with non-MoTeR DNA (Fig. 9A). This array was probably generated through iterative cycles of BIR via D-loop formation, and likely occurred at a time when the 3’ MoTeR copy was still positioned at a chromosome end, or the end of a break. All MoTeR relics have telomere repeats (or at least a partial repeat) at their 3’ termini. In LpKY97, two relics also had TTAGGG tracts at their 5’ truncation boundaries (see Fig. 9F). This arrangement is consistent with a scenario in which breaks occurred in interstitial telomeres at 5’-truncated MoTeR borders and underwent NHEJ, resulting in the internalization of the relic. One relic resides on a segmental duplication, with similarity extending into sequences flanking both sides of the MoTeR (Fig. 9D). In this case, the duplication event was probably unrelated to the element’s presence. Finally, five relics were flanked by single copy sequences, signaling their possible immigration to internal chromosome locations via translocations. Alternatively, it may be that duplications were originally involved but the copies were subsequently lost. In this regard, it is perhaps significant that strain LpKY97 comes from a population with a history of genome shuffling through admixture, which has caused a number of segmental duplications to be purged from some population members (M. Farman, unpublished data).

**Fig. 9.**
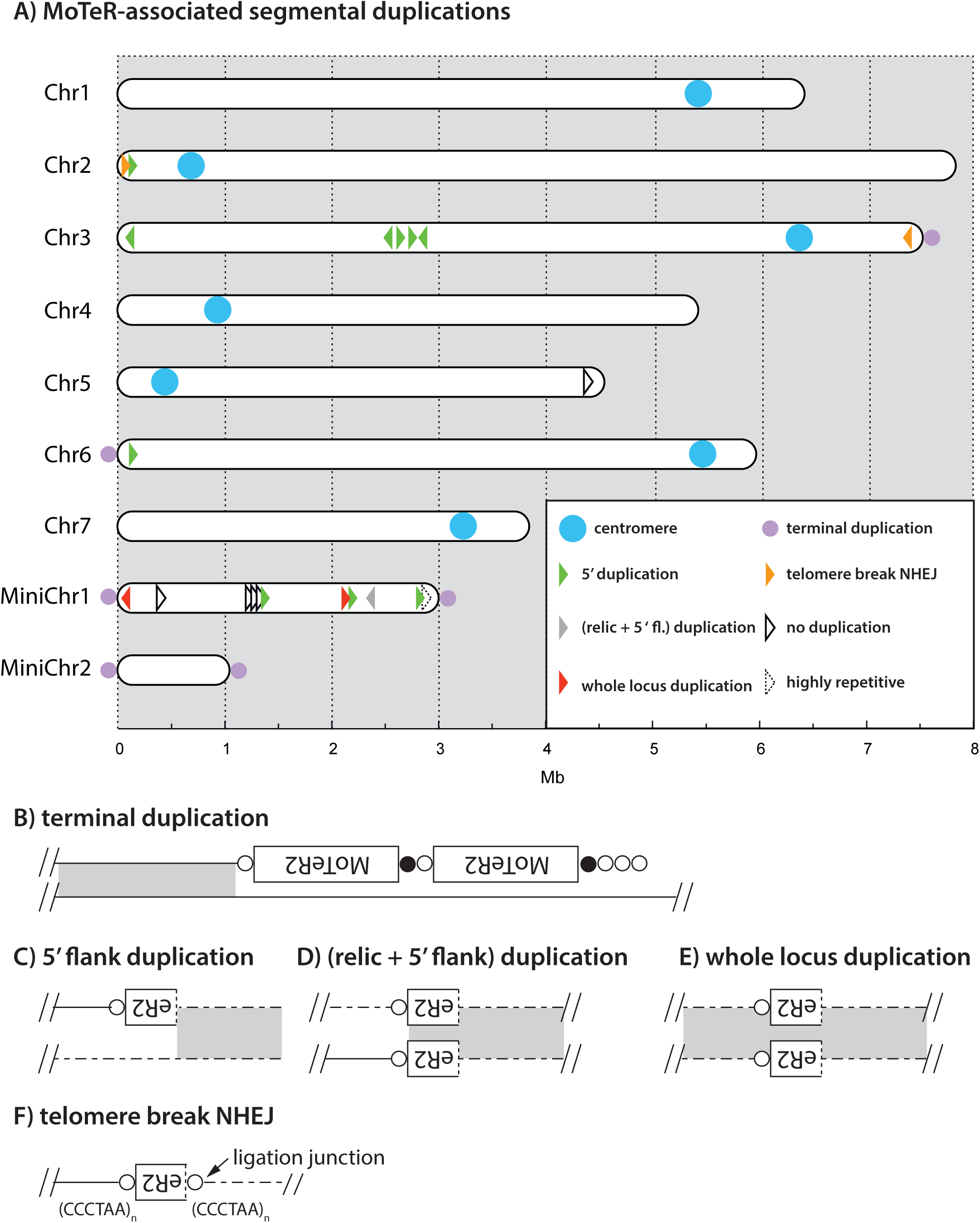
MoTeR-associated, segmental duplications in chromosomes of LpKY97. A) Locations of MoTeR relics in the LpKY97 genome. Each element is represented with an arrowhead representing the relic’s orientation (5’ to 3’). The colors show the nature of the duplication as shown in the legend. The types of duplications observed were: (B) sequences at chromosome ends, immediately adjacent to terminal MoTeR arrays - indicated with lilac dots on the respective chromosome ends; C) 5’-flank duplications beginning at, and extending out from, a relic’s 5’ truncation boundary, (D) duplication of the relic and the 5’-flanking sequence; (E), duplication of the entire locus encompassing a relic. Also shown are examples of putative NHEJ involving broken interstitial telomeres (F). Relics embedded in single copy DNA are shown in black. One insertion was flanked by a highly repeated transposon sequence.

## CONCLUSIONS

With this study, we gained a comprehensive insight into the molecular basis for frequent TRF rearrangements associated with the presence of MoTeRs inserted in the telomeres of *M. oryzae*. Altogether, 109 telomere/subterminal rearrangements were identified and characterized at the molecular level (Table S5). Previously, we showed that rearrangements were largely restricted to TRFs in which the telomeres contained MoTeR insertions - TRFs lacking MoTeRs were generally quite stable, with the lone exception of the rDNA telomere [54]. Here, we show that MoTeR presence is neither necessary nor sufficient to promote extremely high-frequencies of terminal rearrangements. The 28S rRNA gene array is telomeric in several organisms ranging from fungi to plants [18, 19, 68, 76] and, in fungi, the array undergoes frequent terminal truncations ([18, 54, 68, 77] and this study). Consistent with our previous observations, 27 independent rDNA telomere rearrangements were detected through characterization of altered TRFs and analysis of MinION reads. Recently, it has been shown that silent rRNA copies are replicated in mid to late S-phase and have a compact heterochromatin structure [78], which possibly promotes replication stress and subsequent rDNA breakage [79]. It seems reasonable to suppose that *M. oryzae* experiences replication stress in the telomere-adjacent rDNA copies due to a heterochromatin structure resulting from a telomere position effect, and the resulting breaks would then be repaired via *de novo* telomere formation.

Here, we also demonstrated instability in a newly-discovered tandem repeat present in subtelomeres 4 and 8. Contrary to the rDNA, however, this STR, which comprises a much shorter repeat unit, and whose function is unknown, showed TRF length variation in the absence of terminal truncations. This implicates a novel mechanism that does not appear to involve the telomere becoming compromised. In yeast, expansion and contractions of the rDNA repeat are initiated following breakage at a replication fork barrier (Kobayashi et al. 1998), and the resulting breaks are usually repaired via gene conversion, although unequal sister chromatid exchanges are also seen at a not insignificant frequency [80]. The high frequency of STR contractions/expansions in *M. oryzae* implies the existence of a strong replication fork block in these sequences. Why STR breaks should be repaired via gene conversion/unequal crossover, as opposed to *de novo* telomere addition, as was seen in the rDNA, is unclear. However, in humans, there is evidence that the telomere-associated Bloom helicase also interacts with rDNA sequences [81], which possibly indicates a specific role for helicases in the healing of rDNA breaks.

Just as MoTeR presence is not necessary for instability to occur, their presence also does not guarantee it. One MoTeR-containing telomeres in LpKY97 showed no evidence of alterations (TEL 13), while three others (TELs 5, 9 and 11) showed low frequency changes. This provides important insights into the mechanisms by which MoTeRs cause rearrangements and their relative frequencies. First, it shows that the MoTeR sequences *per se* do not compromise the stability or the integrity of the chromosome termini. This is key because both elements contain several unusual sequence motifs, including highly T-rich tracts, blocks of tandem repeats and, most significantly, telomere-like repeats at their 5’ ends. Initially, we suspected that these variant repeats might destabilize the protective telosome, causing the telomere proper to become compromised. However, the rare alterations in TELs 5, 9, 11 and 13 indicates that this is not a major cause of MoTeR-driven rearrangements.

Instead, the key property that distinguished a stable, MoTeR-containing telomere from an unstable one was the absence of an extended interstitial telomere tract between elements. Stable interstitial telomeres had a maximal tract length of (TTAGGG)_2_ (minimal: AGGG), while most of the highly unstable telomeres had much longer inter-MoTeR tracts, with the longest comprising 15 TTAGGG/TTTAGGG repeats attached to the variant repeats found at the MoTeR 5’ ends (consensus: TTCGGG[TTTGGG]_8_). From this we conclude that interstitial telomeres with three or more TTAGGGs are sufficient to promote replication stalling and subsequent breakage. Additional, albeit indirect, evidence that three repeat units are sufficient to promote breaks is that (TTAGGG)_3+_ tracts are entirely absent from internal locations in the majority of *M. oryzae* genomes (data not shown). The lone exceptions are *M. oryzae* strains from foxtail grasses (*Setaria* spp.) which often have numerous long, interstitial telomeres in their subtelomere regions, which likely arise from bouts of telomerase failure (A. Yackzan, M. Rahnama and M. Farman, unpublished data). Such strains exhibit extremely frequent TRF alterations, despite the fact that they lack MoTeRs.

MoTeR1 codes for a reverse transcriptase with a restriction endonuclease-like (REL-ENDO) domain, which suggests that it is a site-specific transposon that targets telomeres [54]. Several of the rearrangements described here involved MoTeR acquisition by a plain telomere, or the addition of new MoTeR copies at the end of an existing array. While both types of occurrences can be explained via transposition, we cannot rule out the possibility that new MoTeRs were copied from other chromosome ends, or from sister chromatids, via BIR (see below).

On the other hand, we would not expect “plain” telomeres to acquire *de novo* MoTeR insertions by BIR because, with the exception of the rDNA, such telomeres are very stable, indicating that recombinogenic end formation through de-protection rarely occurs [45, 54, 82]. As an example, when surveying for telomere alterations in *M. oryzae* rice pathogens that lack MoTeRs, only 14 novel TRFs were identified among more than 1,200 telomeres surveyed [45, 54, 82], and most, if not all, of these were almost certainly due to rDNA truncations. In a similar vein, we do not expect MoTeR-containing telomeres with short interstitial TTAGGG/TTTGGG tracts (e.g. TEL5 & TEL-C) to experience spontaneous breakage. For these reasons, we suspect that MoTeR acquisitions by otherwise stable telomeres possibly originated via transposition. In this regard, we hypothesize that the MoTeR RT protein has the ability to access and cleave telomere repeats that would otherwise be highly protected - an ability that might be further enhanced through interaction with the (TTTGGG)_n_ motifs at MoTeR 5’ ends. Ultimately, however, formal proof that specific rearrangements are due to MoTeR transposition, and an estimation of the frequency of such events, will require the use of a definitive retrotransposition assay [83].

If the MoTeR RT or, more specifically, the predicted endonuclease does have access to an otherwise protected telomere (as is implied by the MoTeRs’ residence therein), then there is also the possibility that it could promote recombination-mediated re-arrangements by generating free DNA ends through telomere cleavage without attendant reverse transcription.

While we were unable to provide conclusive demonstration of *de novo* MoTeR transposition events, several TRF alterations were attributable to new insertions of the MAGGY retrotransposon [84] into MoTeR arrays. The MAGGY reverse transcriptase contains a chromodomain which targets insertions to heterochromatic regions [85]. This suggests that MoTeR arrays might inherit a heterochromatic state from the telomeric regions into which they insert. Given that MoTeR sequences were such attractive MAGGY targets, it is perhaps surprising that we never identified MAGGYs in any of the many MoTeR copies that we characterized previously [54] (and data not shown). Similarly, other *M. oryzae* strains had remarkably stable telomeres even when their genomes contained multiple MAGGY copies (unpublished data). It appears that the element may have been uniquely activated in the particular LpKY97 cultures under study. MAGGY is known to be more active under stress conditions [86] so it is possible that the particular culture of LpKY97 under study suffered an unintended stress.

Telomere fingerprinting of SS cultures from *M. oryzae* strains lacking MoTeRs generally reveals a high degree of telomere stability [45, 54, 82]. However, when a MoTeR inserts into a plain telomere, they inevitably cause a portion of the target site to become internalized, producing an interstitial telomere (Fig. 10A) whose lengths, here, ranged from 0.5 to at least 18 repeat units. Critically, it appears that interstitial telomeres with as few as three TTAGGG repeats (or just two when attached to variant repeats at MoTeR 5’ ends) can promote breakage, with the frequencies of such events reaching ∼10% or more (Fig. 5). As has been shown for Saccharomyces cerevisiae [87], this collateral effect of MoTeR insertion has the potential for genome re-organization on a massive scale, depending on the nature of the break and the mechanism of its repair: If the proximal side of the break contains a telomere vestige, this can probably be extended by telomerase resulting in a “healed” telomere (Fig. 10A.i). Indeed, from the TEL14 signal intensities in Fig. 5D, which show very clearly that arrays shorten much more often than they lengthen, we conclude that most interstitial telomere breaks are simply healed. Nevertheless, the occasional appearance of longer arrays in some cases, points to the operation of other repair processes or, again, possibly transposition.

**Fig. 10.**
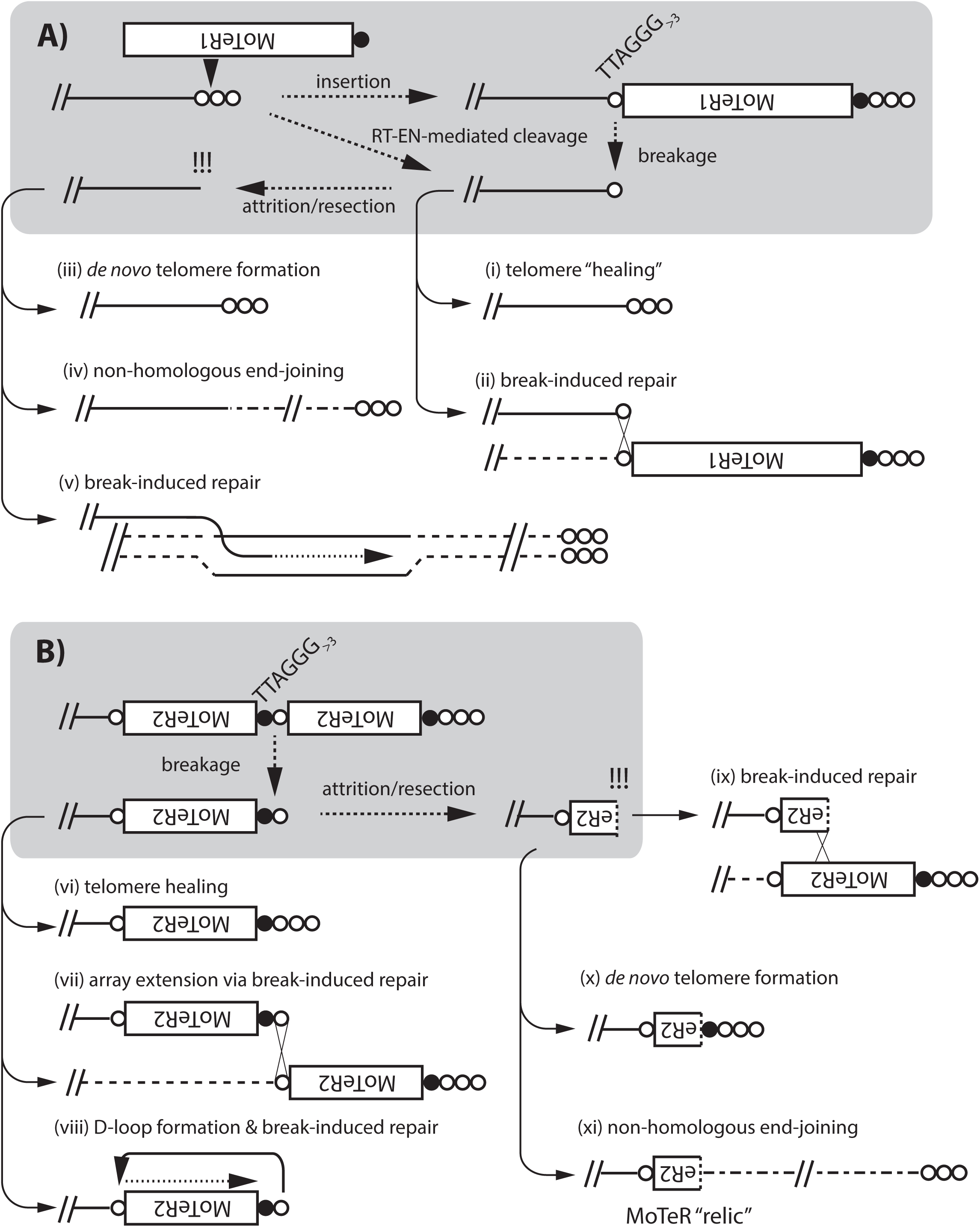
Schematic showing how MoTeR insertions can promote re-organization of the genome interior. A) *End-deprotection.* MoTeR insertion in a plain telomere can generate a fragile site if the insertion position is more than three TTAGGG repeats distal to the telomere boundary. Subsequent breakage would result in the creation of a de-protected end. Alternatively, end de-protection could occur directly via telomere cleavage by the site-specific endonuclease in the MoTeR reverse transcriptase, without a subsequent reverse transcription step. If repair is initiated before the remaining telomere repeats are lost through replicative attrition, or enzymatic resection, the ends could be repaired by telomere “healing” (i), or by ectopic invasion of homologous sequences on a sister chromatid or another chromosome end. Furthermore, if the donor telomere possesses a MoTeR insertion, invasion of an interstitial telomere can result in MoTeR addition (ii). Delayed repair could result in loss of subterminal sequences through attrition/resection, resulting in the generation of a “naked,” recombinogenic end and, possibly, extensive terminal sequence loss. These naked ends can be repaired by *de novo* telomere formation (iii), non homologous end-joining (NHEJ) (iv), or break-induced replication (BIR) - if the free end has homology to another chromosomal sequence (v). B) *MoTeR array “activation.”* Breaks at interstitial telomeres constituting MoTeR-MoTeR junctions can be repaired via healing (vi); BIR using other MoTeR arrays as templates, resulting in array extensions (or contractions) (vii). Alternatively, the free end can invade the same DNA strand, forming a D-loop, with the end being extended by BIR (viii). Finally, attrition/resection of the new terminus will produce a free end comprising internal MoTeR sequences (a truncated MoTeR). This can be repaired via *de novo* telomere addition (ix), BIR (x), or NHEJ (xi). Note that NHEJ will result in the internalization of the truncated version of a formerly telomeric MoTeR. We refer to these internalized elements as MoTeR “relics.”

Break-induced replication (BIR) was evidently involved in some instances of MoTeR array elongation because the newly acquired elements possessed features characteristic of specific MoTeR copies in other telomeres. Examples included MoTeRs that were truncated at specific nucleotide positions, or copies flanked by variant repeat tracts with unusual (TTTGGG)_n_TTAGGG configurations. The operation of BIR was not all that surprising because broken interstitial telomeres should be structurally equivalent to the shortened telomeres that result from experimental deletion of the telomerase (TERT) and, in yeasts and humans, BIR is the default repair pathway for telomeres thus compromised [88–90]. Moreover, *M. oryzae* TERT knockout strains preferentially repair their telomeres via BIR, wherein critically shortened telomeric 3’ ends invade their own subtelomeres at a very short TTAGGG motif (with as few as 1.5 repeats) to form a D-loop. The telomere “tail” then primes replication which extends to the chromosome end. In TERT KO strains, this process is re-iterated many times (or occurs via a rolling circle mechanism, [91]) to generate long tandem arrays at the chromosome ends (Peppers, Rahnama and Farman, unpublished data). For the interstitial telomere breaks observed here, however, it is easy to imagine that once BIR extends to the telomere on the invaded, template strand, the new end thus acquired is then easily maintained through normal telomerase action.

If breaks occur in a very short interstitial telomere, or at the proximal end of an extended one, this would produce a very short telomere vestige that may fail to engage telomerase, causing the adjacent sequences to be lost through replicative attrition, or enzymatic resection. The fate of such ends, and the possibility of wider chromosomal alterations, likely depends on the nature of the terminal sequences. If breaks occur at the boundary of an interstitial telomere and chromosome-unique sequence, attrition/resection will probably result in loss of subterminal sequence. Eventual exposure of a short telomere seed (minimally, T or A) may allow for end- repair via *de novo* telomere formation (iii) [92]. Alternatively, if the ends occur in single copy DNA, they may participate in NHEJ (iv) [93]; or, if they comprise repeated sequences, they potentially could invade a homologous locus and undergo BIR (v). Both of the latter events have the potential to result in the capture and duplication of internal sequence at the chromosome end.

Intuitively, it would seem that breaks occurring at fragile sites at MoTeR-MoTeR junctions would be less likely to produce major chromosomal alterations. Not only do proximal MoTeR sequences act as buffers against loss/exposure of chromosome-unique sequences (Fig. 10B) but they should also provide additional opportunities for end-repair. Exposed telomere repeats can be simply healed (vi), or they could initiate BIR by invading interstitial telomeres between other MoTeRs (vii). Finally, they have the potential to form D-loops by invading proximal interstitial telomeres on the same DNA strand. In this case, the resulting BIR would lead to a precise duplication of the proximal element (viii). Should the telomere repeats be lost through attrition/resection, the MoTeR sequences at the ends should still be eligible for BIR (ix), or *de novo* telomere formation (x). Given the multiple routes by which MoTeR-terminated free ends could be repaired, it might seem unlikely that they would undergo NHEJ (xi). However, the presence of multiple MoTeR relics at internal chromosome locations (Fig. 9A) indicates that such occurrences are, in fact, far from rare.

MoTeR-induced breaks that undergo NHEJ, or utilize internal loci as BIR templates, have the potential to cause internal sequences to be duplicated at, or translocated to, chromosome ends. In a prior study we were able to pin-point the boundary of one such duplication to a transposon present in both the terminal and interior locations, directly implicating BIR as the driving mechanism [18]. Unfortunately, in the present study, we were unable to characterize the boundary of the duplication that gave rise to the novel TRF-C, but it seems reasonable to suppose that a MoTeR-induced break might have been the initiating event. Evidence supporting direct MoTeR involvement in the creation of segmental duplications in the LpKY97 genome came from 12 internal relics that defined duplication boundaries and, therefore, serve as historical markers of terminally-directed duplication events.

Segmental duplications play important roles in gene and genome evolution, by providing raw materials for functional diversification through the creation and expansion of gene families [94, 95]. By generating fragile sites at chromosome ends, MoTeRs not only promote repair-driven duplications/translocation events, but they direct these processes to the chromosome regions with the greatest evolutionary potential. Thus, one can imagine a dynamic in which genes are duplicated in the subterminal regions where they experience accelerated evolution and, as result, have an enhanced potential for neo-functionalization. Moreover, newly-evolved genes with useful functions then have a means of being recruited back to the more stable genome interior, via subsequent terminalization events. Thus, the impressive telomere dynamicism generated through MoTeR invasion of telomeres can be leveraged for maximal evolutionary benefit.

## Acknowledgments

The merged, SNP-corrected, MinION genome assembly for LpKY97 has been deposited at NCBI and has the accession number XXXXX. The SS4-20, SS6-1 and SS15-11 raw read data are available in the Sequence Read Archives under BioProject PRJNA579424. The authors would like to acknowledge technical support from David Thornbury, Melanie Heist, Rebekah Ellsworth, and Jennifer Webb; and contributions from Katelyn Richard and Alex Stewart. This work was supported by the following grants from the National Science Foundation: MCB-0135462, MCB-0653930 and MCB-1716491.

## Declarations

None of the authors of this manuscript have any competing interests.

## SUPPLEMENTAL RESULTS AND DISCUSSION

### Additional terminal rearrangements identified by using telomere-linked probes to monitor instability at individual chromosome ends

The restriction enzyme, *Pst*I, does not cut in either MoTeR element which means that each telomeric restriction fragment (TRF) contains a short segment of chromosome-unique sequence adjacent to the MoTeR insertion. Sequences adjacent to the proximal MoTeR, or distal to the terminal *Pst*I site were used as probes to detect alterations of individual TRFs among SS isolates. **TEL7**. Targeted cloning yielded a number of plasmids with insert sizes of approximately 10.5 kb, and which shared the same telomere adjacent sequence mapping to the top of chromosome 4 (=TEL7). In most SS cultures, the TLP7 probe hybridized to fragments with a molecular size of ∼10.5 kb fragment (Figure S10A, bands i and ii). Four isolates (SSs 12, 13, 17 & 18) had signals at a position corresponding to ∼ 16 kb (band iv), one (SS4) had one slightly larger than 16 kb (band iii), and the 2G1SS1 parent culture had a signal at ∼13 kb.

MinION reads for 2G4 SSs 6-1 and 15-11 revealed that the 10.5 kb TRF - which was also present in the original LpKY97 culture (Fig. 1) but not 2G1SS1 (Fig. S10A) - had a single, full-length MoTeR1 insertion in the telomere with a short interstitial telomere repeat (TTAGGG)_2_ (Fig. S10B, i & ii). In contrast, most reads from SS4-20 identified a TEL7 variant with four truncated MoTeR copies distal to the intact element (iii) (Fig. 10A, band iii). These four elements were duplicates of truncated variants found at the end of TEL-D (Fig. S5). Considering that the parental telomere lacks obvious break-inducing features, we hypothesize that the first three elements were acquired through a multi-step process. Step 1 likely involved transposition of a composite MoTeR cassette transcribed from TEL-D; in step 2, a terminal duplicate of these last three elements was generated by D-loop formation & extension; step 3 involved terminal resection that eliminated two distal tMoTeR copies; and, finally, step 4 was healing using the TTAGGG seed at the tMoTeR2:tMoTeR1 junction. Under this scenario, the slightly shorter variant with just two truncated elements distal to the intact MoTeR1 could be the product of more extensive resection prior to healing (iv). The low abundance of variant (iv) in SS4-20 (and its similar size) is probably why a corresponding band is not visible in the SS4-20 lane but it likely corresponds to band “iv” seen in SS 12, 13, 17 and 18. Finally, we were unable to characterize the ∼13 kb TRF in LpKY97 but given that it is slightly smaller than variant “iv,” it possibly comprises an array with a telomere-healed break at the (TTAGGG)_3_ repeat.

#### TEL10

Targeted cloning yielded a number of plasmids with inserts of approximately 11.5 kb and 15 kb which shared the same chromosome unique sequence and, therefore, represented parental and rearranged versions of the same chromosome end (TEL10). This was confirmed by using the unique sequence adjacent to the *Pst*I site to probe *Pst*I digested DNAs from a number of SS cultures. As expected, strain 2G3SS4 which yielded the clones with the 11.5 kb inserts exhibited a hybridization signal at a position corresponding to 11.5 kb, while the remaining cultures - including 2G3SS7, the strain that produced the larger insert - all had signals at a position corresponding to ∼16 kb (Fig. S11A). MinION sequencing revealed that the larger, parental TRF (variant 10.i) had four truncated MoTeR1s inserted tandemly in the telomere, two of which were missing the same amount of 5’ sequence (264 nt). The TRF was unusual in that it also contained a second, drastically truncated MoTeR1 (MoTeR1 relic) at an “internal” location approximately 2 kb away from the telomere but whose orientation was inverted with respect to the telomeric copies. Also present was an inverted, interstitial telomere just 300 bp away from the start of the TTAGGG repeats at the MoTeR array border. It is clear that the internal relic was once telomeric because it had 1.5 telomere repeat units at its 3’ end. In SS SS4-20, TRF10-1 had an identical organization, except that it possessed just one copy of the larger truncated MoTeR1 (Fig. S11B, variant ii). As was seen with TEL14 (Fig. 5D), SS cultures possessing TRF10.i often exhibited signals at the same position as TRF10.ii (Fig. S11A), which suggests that 10.i also undergoes spontaneous truncations at a high frequency, that gives rise to structure 10.ii in some nuclei. However, this conclusion was not supported by the sequence data because, while two variants were identified among reads for SS15-11, their sizes were ∼15 kb and > 20 kb.

Furthermore, because the MoTeR array contained no long interstitial telomeres (Fig. S11B), there was no reason to suspect it would undergo recurrent breakage. Instead, the identification (and structure) of the longer (20 kb+) variant, and its presence - albeit as another faint band - in several SS cultures that possessed the short form, presented an alternative mechanism that fully explained the data. Specifically, the large and short variants are the expected products of an unequal sister chromatid exchange between the two larger tMoTeR1 insertions. It is not clear what would promote such a high level of recombination in this array.

#### TEL-B *&* TEL-D

Targeted cloning of TRFs from 2G3SSs 2, 4, 5, 6, and 7 resulted in the isolation of an 8.3 kb terminal fragment that contains a single MoTeR1 insertion in TEL-B on minichromosome 1 (Fig. S12A, variant B.iv). The corresponding TLP-B produced three main hybridization signals at 8.3 kb, ∼11 kb and >12 kb (Fig. S12C, right-hand panel). The 11 kb band remained unaltered in all SS cultures consistent with its origination from an internal chromosome location that lacks MoTeR sequences, interstitial telomeres, or other obvious de-stabilizing features (data not shown). In contrast, both the upper and lower bands (>12 kb & 8.3 kb) showed rearrangements in certain SS isolates. The lower band exhibited varying hybridization intensities among isolates. It was very faint in SS4, and was largely replaced by a strong signal at ∼16 kb, and a weaker one at ∼14 kb (Fig. S12C, B.i and B.iii, respectively); and it was completely absent in 2G1SS1, which instead possessed novel bands of ∼11.7 kb, 10 kb, and faint signals at ∼9.5 kb, and a high molecular weight fragment (Fig. S12C, right-hand panel). The hybridization signal from the >12 kb fragment (D.i), which contains TEL-D, was absent in SSs 5, 6, 8 and 9 which, instead, showed doubly intense bands at ∼8.3 kb (D.ii*, Fig. S12C).

Inspection of the MinION assembly of SS4-20 revealed that the original 8.3 kb TRF, which contained a single MoTeR1 insertion (Fig. S12A, variant B.iv), was a truncated version of a longer TRF that contained a multi-MoTeR array with an extended interstitial telomere distal to the aforementioned MoTeR1 (Fig. S12A, variants B.i, ii and iii). Normally, the fact that the majority of SS isolates possessed variant B.iv would imply that this short form is the original parental structure. However, given the absence of obvious destabilizing features, it is more likely that the shorter TRF-B forms were derived from longer variants via breakage and healing of the interstitial telomeres at the MoTeR junctions. The absence of variant B.iv in SSG1SS1 is consistent with this interpretation, and it appears that the most of the SS isolates might have inherited a truncated variant that arose in the original colony while it was being grown to generate spores for the first round of plant infection. The structure of variant B.v, is most easily explained as a result of MoTeR2 insertion at the internal TTAGGG motif in the B.iv telomere [i.e., TTCGGG(TTTGGG)_5_<u>TTAGGG</u>TTTGGG(TTAGGG)_28-33_]. In this case, the additional TTAGGG repeats found in the newly-created interstitial telomere would have been contributed by the invading MoTeR. Alternatively, the interstitial telomere could have become altered through replication slippage.

The MinION assemblies revealed that TEL-D contains a complex MoTeR array spanning a total of 48 kb, and comprises 18 separate tMoTeR1 and tMoTeR2 insertions (Fig. S5). One of the tMoTeR1 copies is interrupted by a MAGGY insertion (Fig. S12B, variant D.i) which explains why the 12kb+ hybridizing band is not telomeric (see Fig. S12C, left-hand panel). Inspection of MinION reads for the SS culture 6-1, which lacked the 12kb+ signal, revealed a D.ii variant that is very similar in size to the B.iv fragment. D.ii likely arose following a break at the interstitial telomere tract, (TTAGGG)_3_, which is distal to the most proximal tMoTeR1 copy, with the break having been subsequently healed by telomere addition (Fig. S12B.ii). The original tMoTeR1 truncation boundary (position 362) was conserved, indicating that at least a portion of a TTAGGG repeat at the breakpoint was retained and served as a seed for telomere healing. Interestingly, the complete absence of reads spanning the MoTeR1/MAGGY junctions indicates that the sequences beyond the truncation point were eradicated from the SS6-1 and SS15-11 genomes.

#### TEL-C

Targeted cloning using DNA from 2G3SSs 4 and 6 resulted in the recovery of plasmids with inserts of ∼ 7.6 and 9.3 kb, respectively. The cloned had the same sequence adjacent to the *Pst*I cloning site, consistent with their representing variants of the same TRF which was derived from the start of mini-chromosome 2 (i.e., TEL-C). This sequence was duplicated at an internal position on mini-chromosome 1. The shorter TRF contained three truncated MoTeR copies with an intact MoTeR1 in the distal position, while the longer fragment contained an intact MoTeR2 insertion distally to the MoTeR1 (Fig. S13A).

As expected, the TLP-C probe bound primarily to two loci in genomic DNA from the SS isolates. One, a 7 kb fragment corresponding to the internal locus, showed no rearrangement, while the other displayed two alternate forms with molecular sizes approximating (∼7.5 and 9.5 kb). These latter fragments clearly corresponded to the two cloned variants, with the shorter TRF (i) being the predominant form. Cultures 5, 6, 7 and 8 all exhibited the larger (ii) form (Fig. S13A, right-hand panel), which was consistent with their clonal relationship, as evidenced by telomere profile identity (Fig. S13A, left-hand panel). The observed alterations in TTTGGG tracts lengths at the presumed insertion site (Fig. S15B.ii) could be explained by the utilization of a retrotransposition template carrying these repeats at its 5’ and 3’ termini, or through replication stalling/slippage without breakage.

All of the MinION reads from SS isolates 4-20 and 15-11 were consistent with the 7.6 kb TRF (Fig. S13B.i) and there was no evidence for the emergence of other variants within the SS cultures that possessed this form. Consequently, aside from the one alteration recovered in SS6 (and visible in its sister cultures), this TRF appears to be generally quite stable. No further rearrangements were detected among the small number of TEL-C reads from SS6-1. However, based on the presence of an extended interstitial telomere between the two intact MoTeR copies, we predict that breakage would promote further alterations at an appreciable frequency.

### Telomere rearrangements identified among MinION reads from single spore cultures 2G4SS4-20, 2G4SS6-1 and 2G4SS15-11

Unless otherwise stated, telomere instability was documented by interrogating only those reads that started in the chromosome-unique region proximal to the telomere and terminated in telomere repeats (or *vice versa*). In other words, they spanned entire MoTeR/subtelomeric repeat arrays (note: the C-rich strand of the chromosome end was rarely captured via MinION sequencing).

#### TEL2

Inspection of MinION reads showed that TEL2 was contained on a *Pst*I fragment of between ∼46 kb and 48 kb (Fig. S13A) and the subterminal sequences comprise another tandem repeat array. Note that this array is disqualified from being classified as a “subtelomeric” tandem repeat due to its presence at a single chromosome end (see “Terminology”). In SS4-20 and SS6-1, the telomere contained a truncated MoTeR2 and an intact MoTeR1, with short interstitial repeats (≤3 TTAGGGs) (Fig. S13A, i & ii). Differences in the sequence of variant repeats subtending the telomere suggests that the TEL2.ii form was derived from TEL2.i via replication slippage. In SS15-11, a truncated MoTeR1 was found inserted between the tMoTeR2 and MoTeR1 copies and two different terminal organizations were identified (Fig. S13A. iii & iv). The newly acquired tMoTeR1 was truncated at the same position as two copies found in TEL4, suggesting that it was acquired via gene conversion, or that TEL2 and TEL4 underwent an exchange of sequence. These scenarios also account for the acquisition of the drastically truncated MoTeR1 that is found in a terminal position in TEL4. Variant TEL2.iv presumably arose from TEL2.iii through an interstitial telomere break, followed by resection and, eventually, *de novo* telomere formation.

#### TEL6

The MinION reads containing TEL6 revealed the presence of multiple MoTeR1 and MoTeR2 insertions, with most of the molecules harboring between five and 10 element copies (Fig. S14A). Several of these MoTeR arrays were unusual in having abnormally long and complex interstitial telomeres comprising alternating runs of the various TTAGGG-like motifs (e.g., 6.iii - vi); and in two cases, the telomere proper was subtended by variant repeats with even more complex organizations (6.iv & 6.vi). The 6.i and 6.ii variants had a truncated tMoTeR1 as the most proximal array element, which was replaced by an intact MoTeR1 in all other forms.

The 6.ii arrangement probably arose from 6.i via intrachromatid or unequal sister chromatid exchanges involving MoTeR1 copies 2 and 4 (Fig. S14B), while variant iii is the likely product of intrachromatid or unequal sister chromatid exchange between the tMoTeR1 and MoTeR1 copy 2 Fig. S14C). Variant 6.iv almost certainly arose from 6.iii via interstitial telomere breakage and healing; and it seems likely that 6.vi was generated from 6.iii, 6.iv or 6.v by the same mechanism. On the other hand, 6.v was probably derived from 6.iii through an intrachromatid or unequal sister chromatid exchange between MoTeR1 copies 3 and 4.

#### TEL12

Most reads containing TEL2 supported a telomere structure comprising insertions of three truncated MoTeR1s in a proximal position and intact, distal copies of MoTeR1 and MoTeR2, with long interstitial telomere/variant repeats between them (Fig. S13B.i & ii). Some reads revealed a shorter TTAGGG tract subtending the telomere (7 TTTGGG repeats vs. 10), consistent with replication slippage. Variant 12.iii likely arose from 12.ii via breakage and healing, while variant 12.iv appeared to have originated via D-loop formation and extension - which best explains the identical structure of the two distal interstitial telomeres.

#### TEL-A

The TRF terminating in TEL-A had a highly variable structure as six different variants were identified among the MinION reads. The majority of reads had a truncated MoTeR1 in the most proximal array position with an intact MoTeR1 adjacent to it, but then there was variation in the distal composition (Fig S15A). TEL-A.i had a MoTeR2 in the distal position, A.v had a second intact MoTeR1, while A.ii was simply capped with a telomere. The A.ii form was likely derived from A.i simply by breakage and healing. On the other hand the A.v variant appears to have originated through recombination with another telomeric array, causing acquisition of the characteristic (TTTGGG)_10_TTAGGGTTTGGG(TTAGGG)_2_ interstitial telomere tract that is found in other telomeres (TEL6, TEL14, TEL-C). All of the TEL-A reads from SS6-1 had a MAGGY inserted in the proximal tMOTER1 (Fig. S15A, 6.vi). However, none of the reads extended to the chromosome thereby precluding identification of other possible alterations.

TEL-A.iii had its probable origin in an intrachromosomal, or unequal sister chromatid exchange that paired the truncated MoTeR1 with the intact copy, while A.iv has a structure consistent with an interstitial telomere break and healing of A.iii.

**Table S1.**
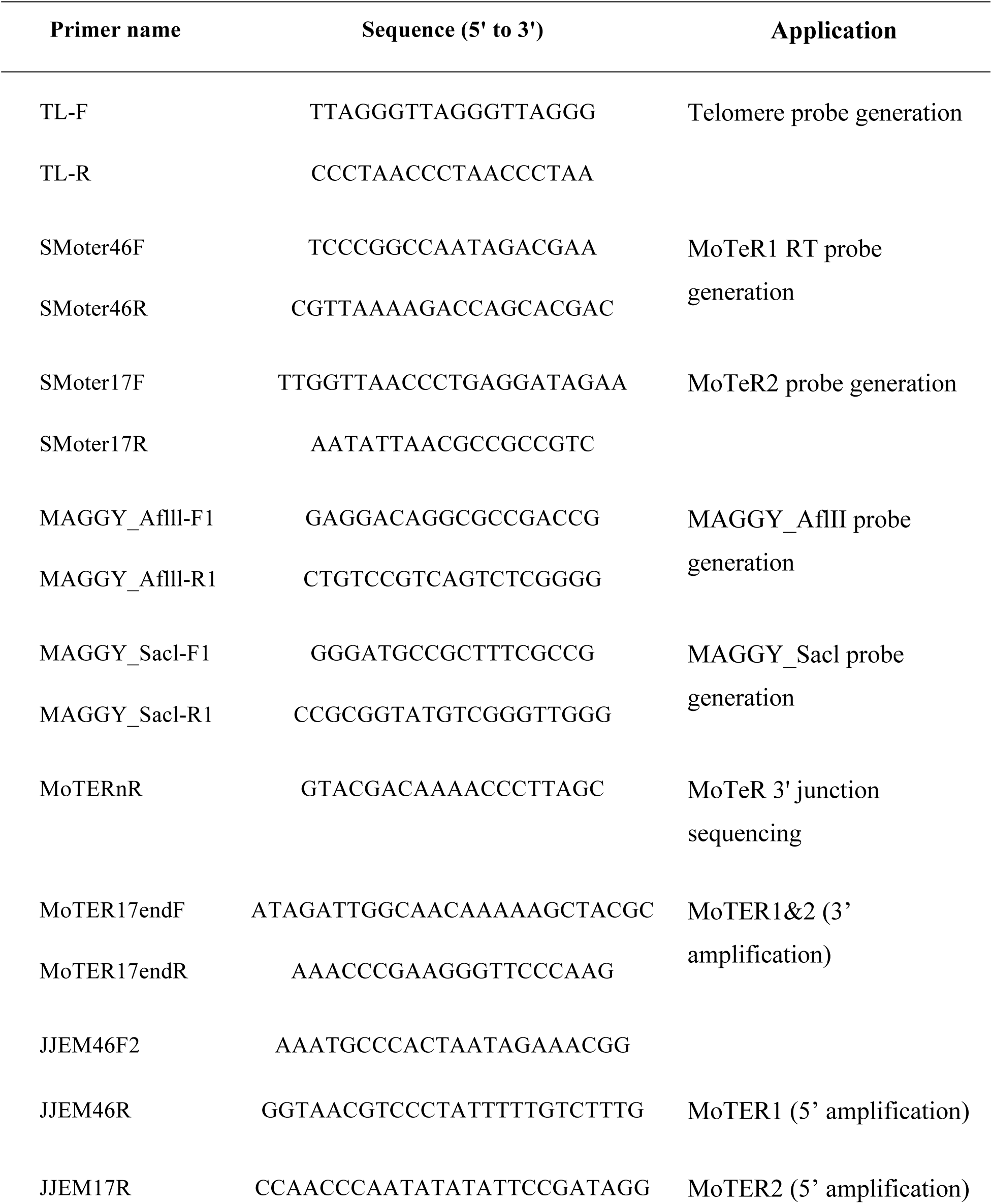
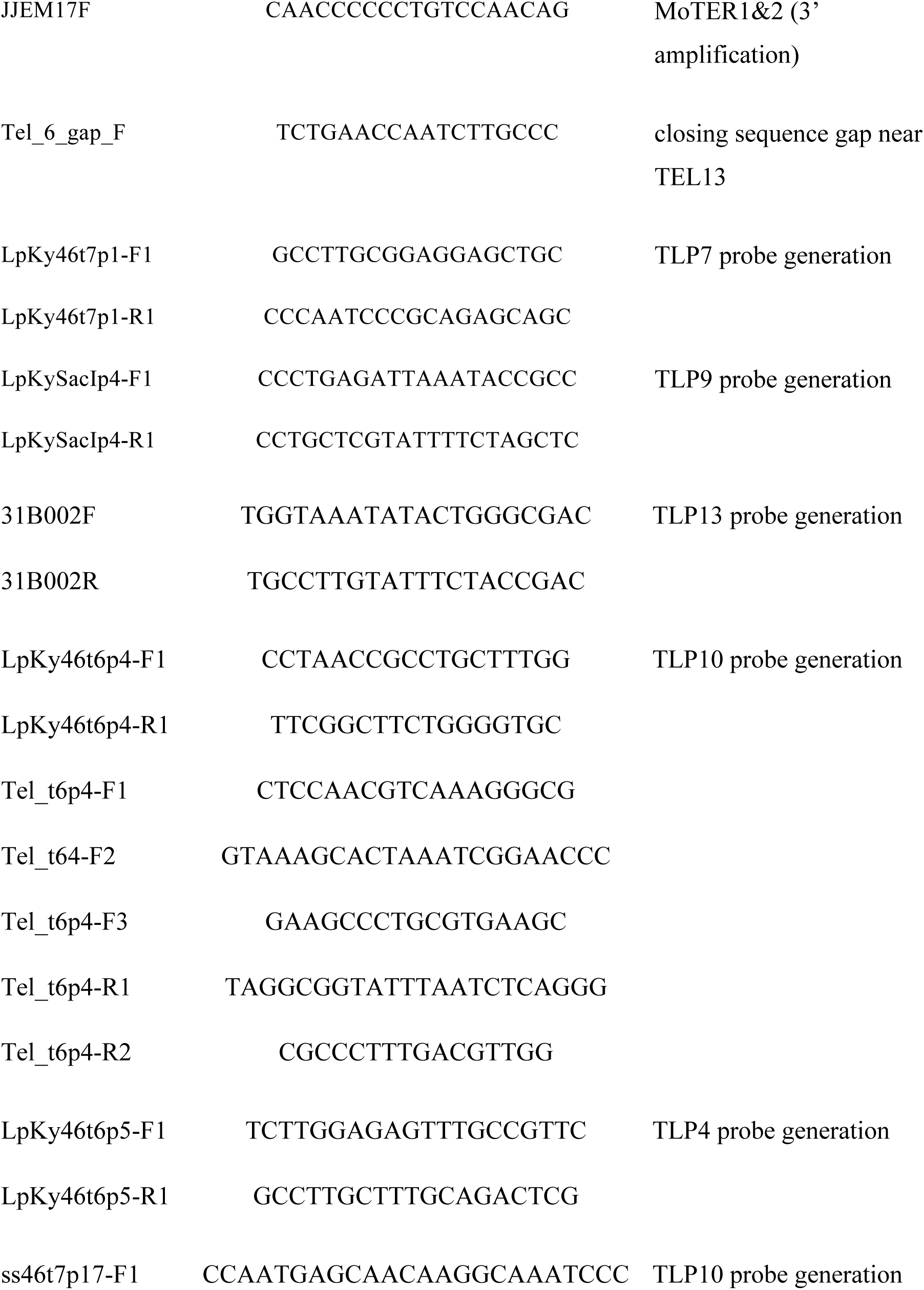
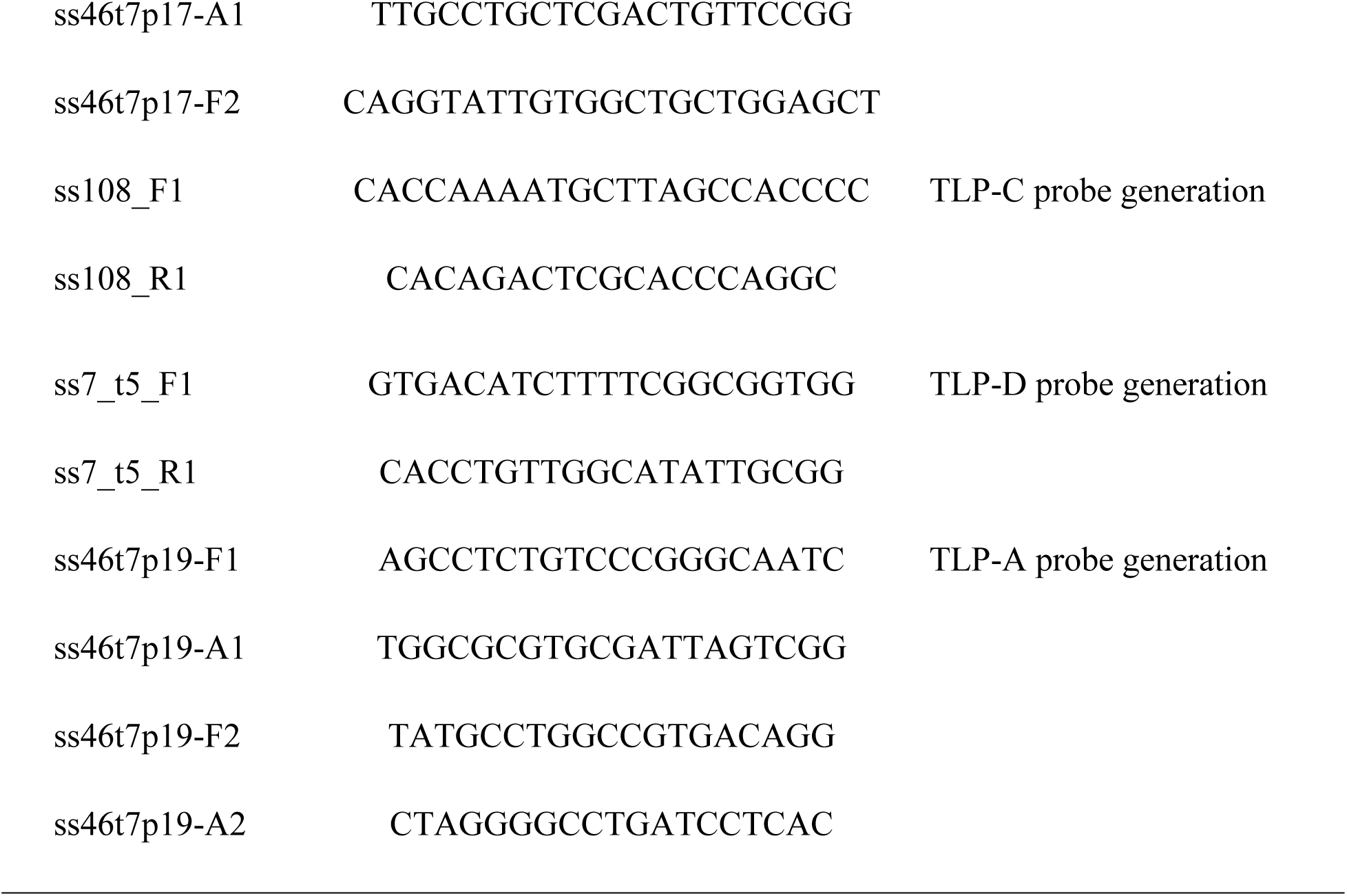
Sequences of primers used in this study.

**Table S2.**
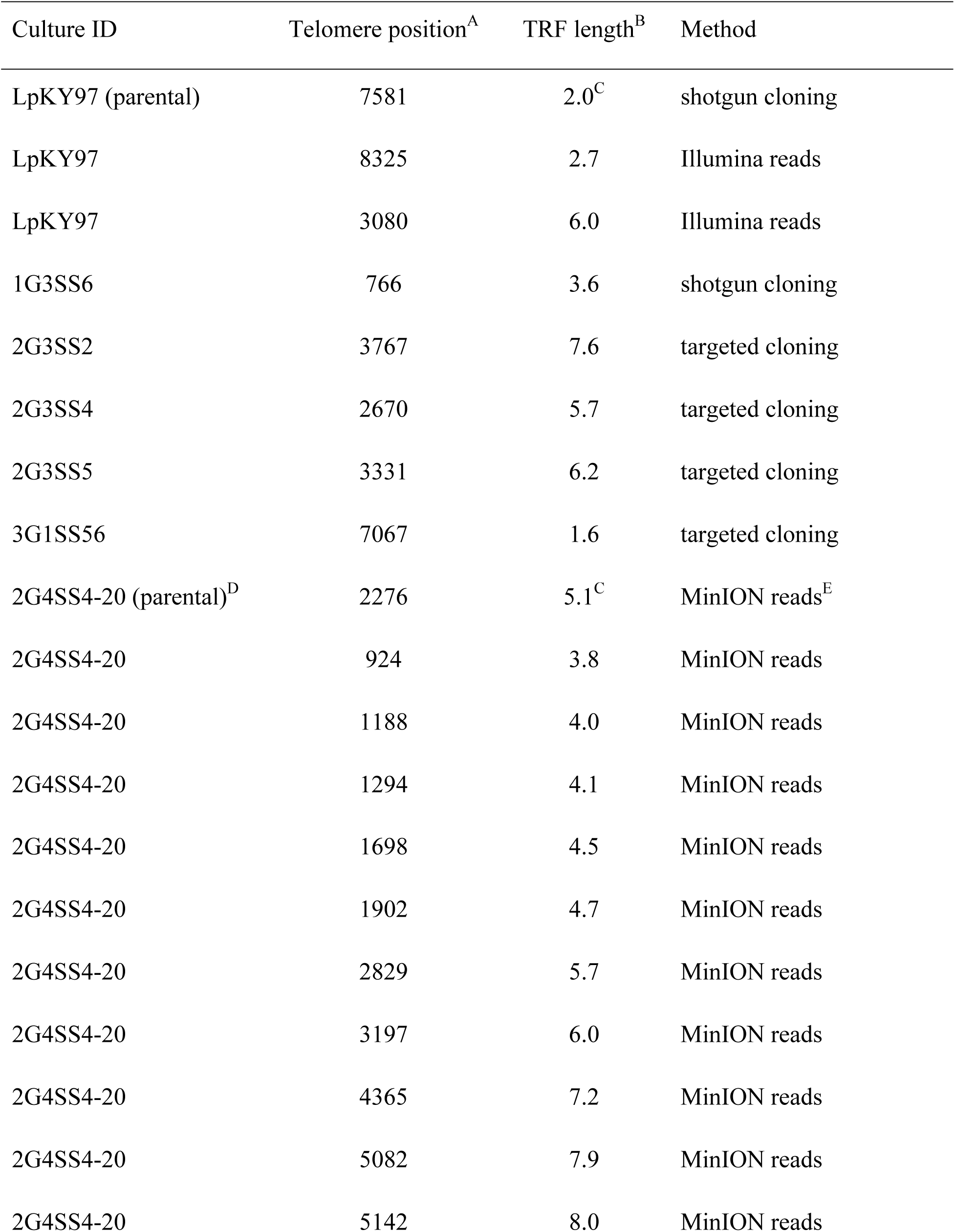
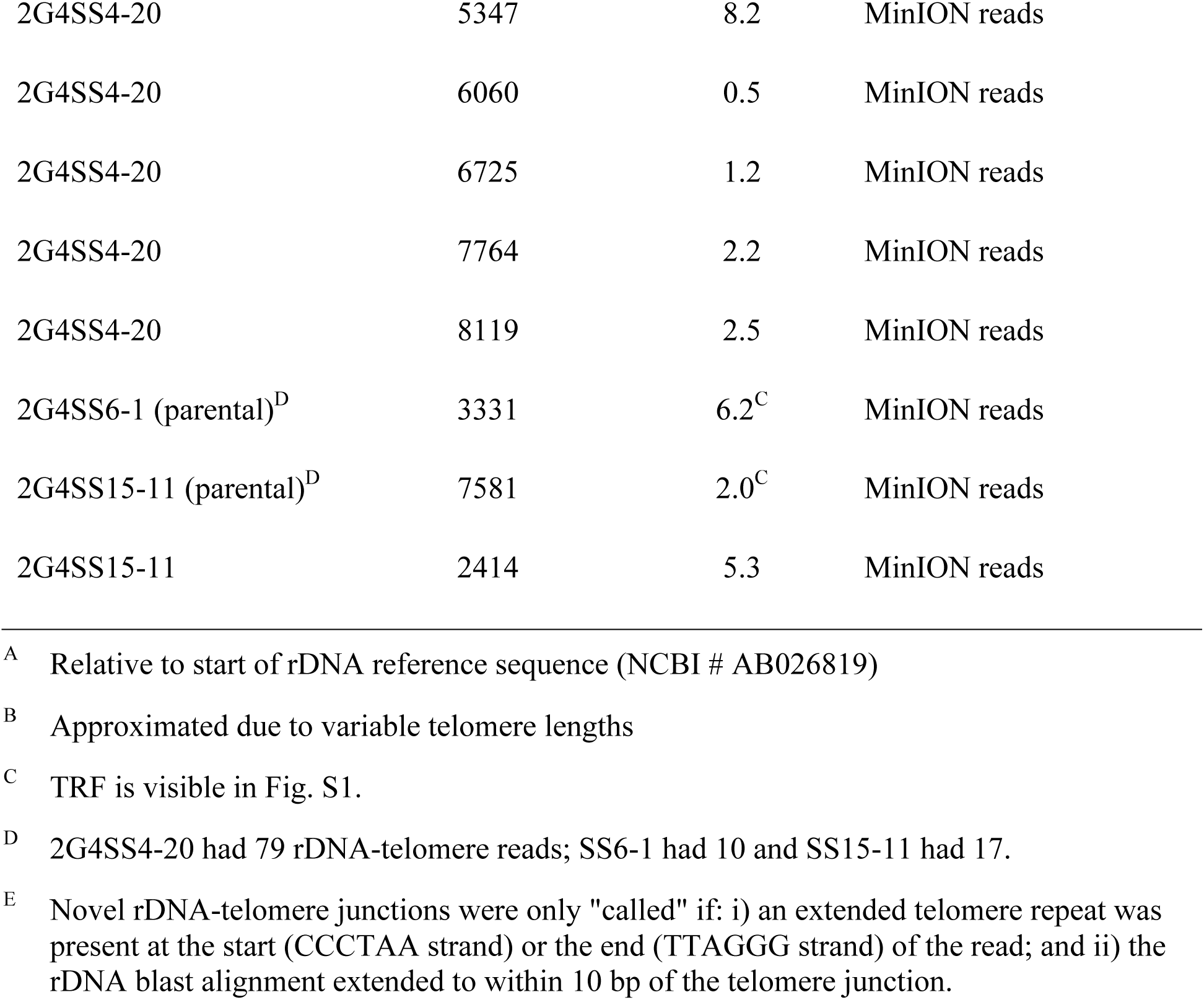
**rDNA truncations identified in this study**

| Culture ID | Telomere position<sup>A</sup> | TRF length<sup>B</sup> | Method |
| --- | --- | --- | --- |
| LpKY97 (parental) | 7581 | 2.0<sup>C</sup> | shotgun cloning |
| LpKY97 | 8325 | 2.7 | Illumina reads |
| LpKY97 | 3080 | 6.0 | Illumina reads |
| 1G3SS6 | 766 | 3.6 | shotgun cloning |
| 2G3SS2 | 3767 | 7.6 | targeted cloning |
| 2G3SS4 | 2670 | 5.7 | targeted cloning |
| 2G3SS5 | 3331 | 6.2 | targeted cloning |
| 3G1SS56 | 7067 | 1.6 | targeted cloning |
| 2G4SS4-20 (parental)<sup>D</sup> | 2276 | 5.1<sup>C</sup> | MinION reads<sup>E</sup> |
| 2G4SS4-20 | 924 | 3.8 | MinION reads |
| 2G4SS4-20 | 1188 | 4.0 | MinION reads |
| 2G4SS4-20 | 1294 | 4.1 | MinION reads |
| 2G4SS4-20 | 1698 | 4.5 | MinION reads |
| 2G4SS4-20 | 1902 | 4.7 | MinION reads |
| 2G4SS4-20 | 2829 | 5.7 | MinION reads |
| 2G4SS4-20 | 3197 | 6.0 | MinION reads |
| 2G4SS4-20 | 4365 | 7.2 | MinION reads |
| 2G4SS4-20 | 5082 | 7.9 | MinION reads |
| 2G4SS4-20 | 5142 | 8.0 | MinION reads |
| 2G4SS4-20 | 5347 | 8.2 | MinION reads |
| 2G4SS4-20 | 6060 | 0.5 | MinION reads |
| 2G4SS4-20 | 6725 | 1.2 | MinION reads |
| 2G4SS4-20 | 7764 | 2.2 | MinION reads |
| 2G4SS4-20 | 8119 | 2.5 | MinION reads |
| 2G4SS6-1 (parental)<sup>D</sup> | 3331 | 6.2<sup>C</sup> | MinION reads |
| 2G4SS15-11 (parental)<sup>D</sup> | 7581 | 2.0<sup>C</sup> | MinION reads |
| 2G4SS15-11 | 2414 | 5.3 | MinION reads |

<sup>A</sup> Relative to start of rDNA reference sequence (NCBI # AB026819)
<sup>B</sup> Approximated due to variable telomere lengths
<sup>C</sup> TRF is visible in Fig. S1.
<sup>D</sup> 2G4SS4-20 had 79 rDNA-telomere reads; SS6-1 had 10 and SS15-11 had 17.
<sup>E</sup> Novel rDNA-telomere junctions were only "called" if: i) an extended telomere repeat was present at the start (CCCTAA strand) or the end (TTAGGG strand) of the read; and ii) the rDNA blast alignment extended to within 10 bp of the telomere junction.

**Table S3.** MAGGY insertion positions

| Single Spore ## | Insertion position<br>(Relative to Telomere) | MAGGY orientation<sup>A</sup> | MoTeR target<sup>B</sup> |
| --- | --- | --- | --- |
| 0<sup>C</sup> | 3505 | + | M1 |
| 50 | 2799 | + | M1<sup>D,E</sup> |
| 56/145 | 854 | + | M1/M2<sup>D,E</sup> |
| 72 | 4205 | - | M1<sup>E</sup> |
| 61 | 2265 | - | ? |
| 80 | 2860 | + | M2<sup>F</sup> |
| 108 | 4565 | - | M2<sup>F</sup> |
| 109 | 4565 | - | M2<sup>F</sup> |
| 115(1) | 4160 | + | M2<sup>F</sup> |
| 115(2) | 5866 | + | M2<sup>F</sup> |
| 120 | 5665 | - | M2<sup>F</sup> |
| 2G4SS15-11 | 1333 | - | M1<sup>G</sup> |
| 2G4SS15-11 | ?<sup>H</sup> | + | M1<sup>G</sup> |

<sup>A</sup> orientation is defined relative to MoTeR orientation<sup>B</sup> M1; MoTeR1 and M2; MoTeR2<sup>C</sup> Insertion present in starting culture<sup>D</sup> insertion occurred in 2nd element in the tandem array with M2 in distal position (see Fig. 4A)<sup>E</sup> determined by cloning<sup>F</sup> determined by Southern hybridization<sup>G</sup> determined from MinION sequence reads
<sup>H</sup> unknown because read did not extend to telomere

**Table S4.**
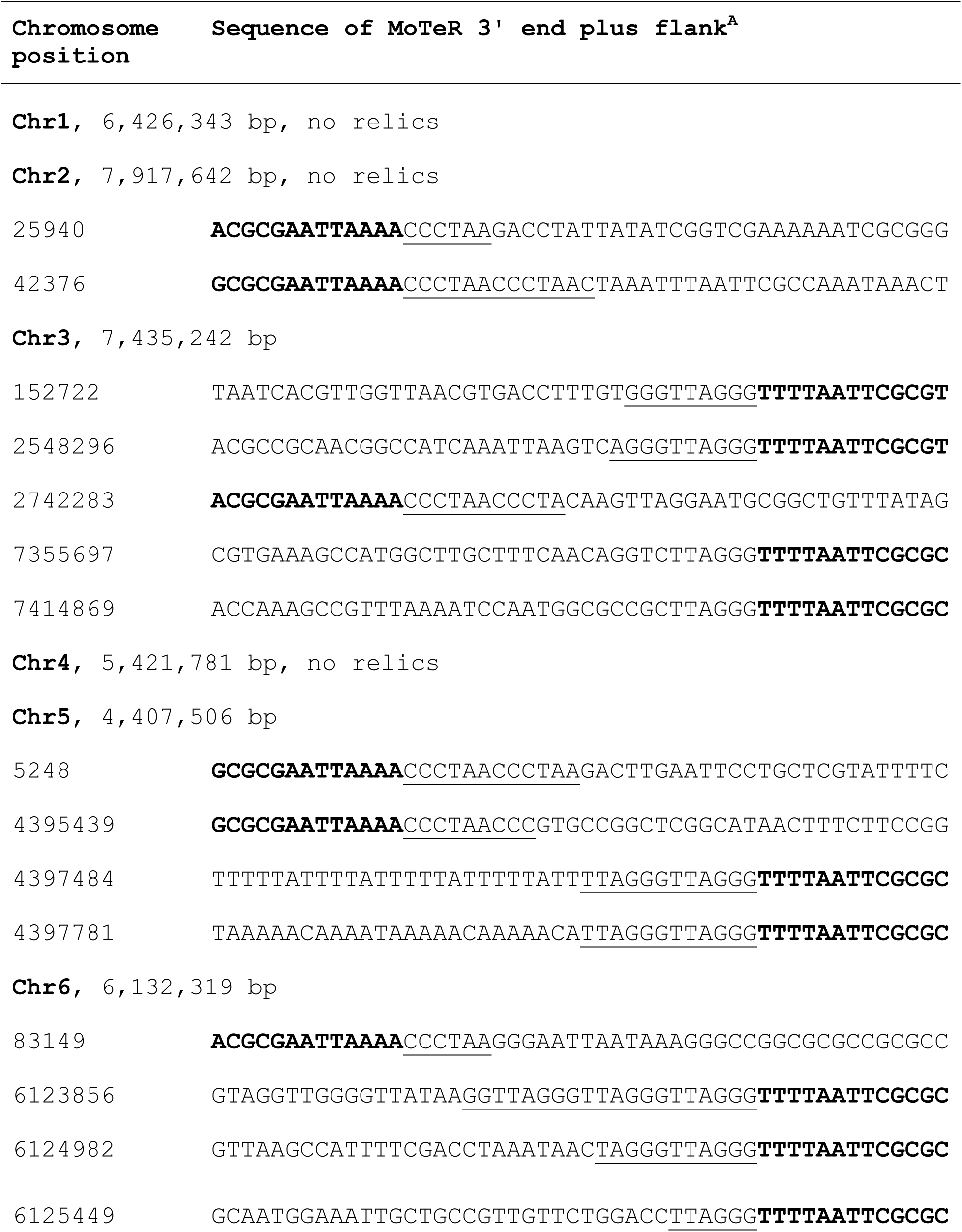
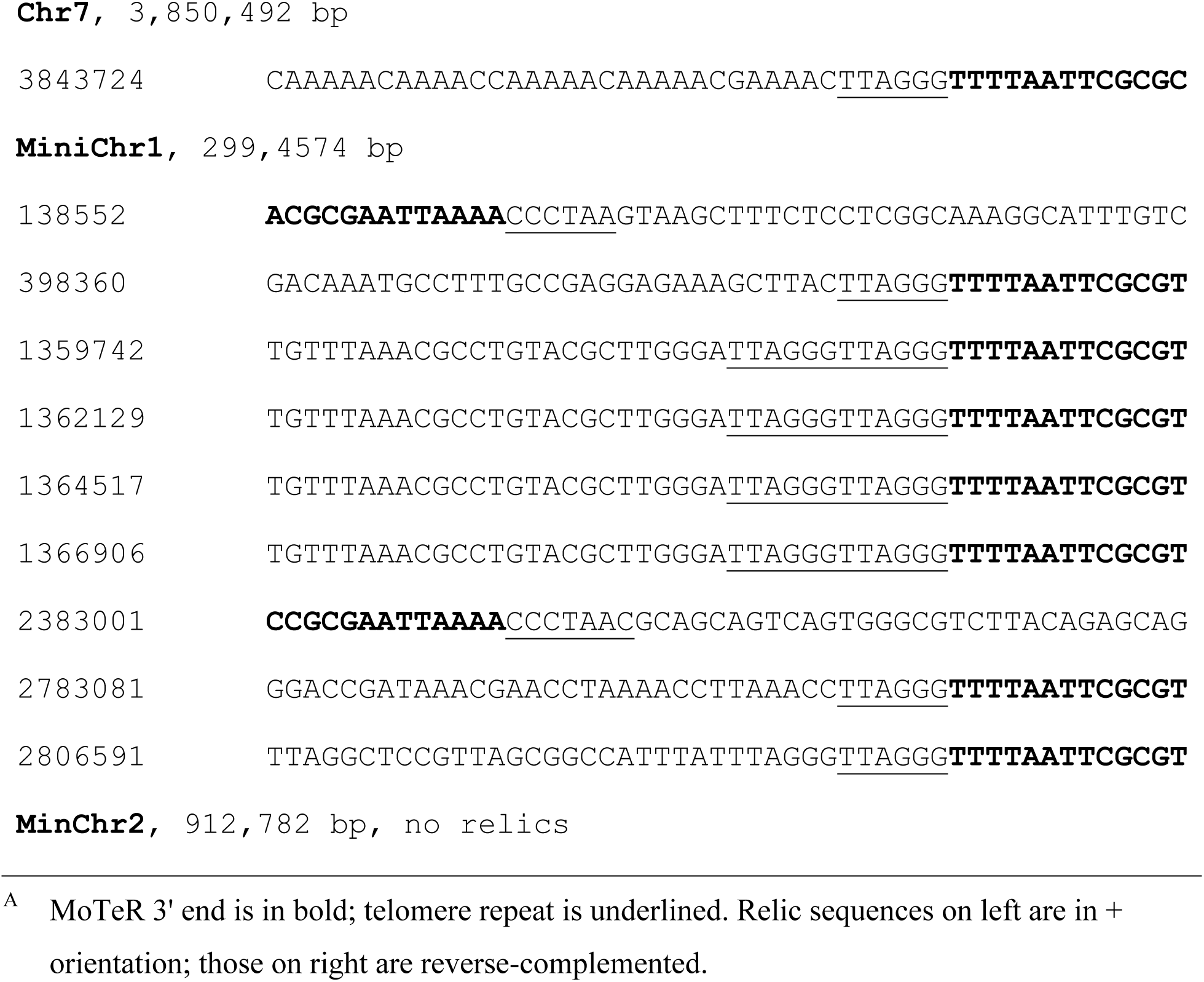
MoTeR relics in the LpKY97 genome.

**Table S5.**
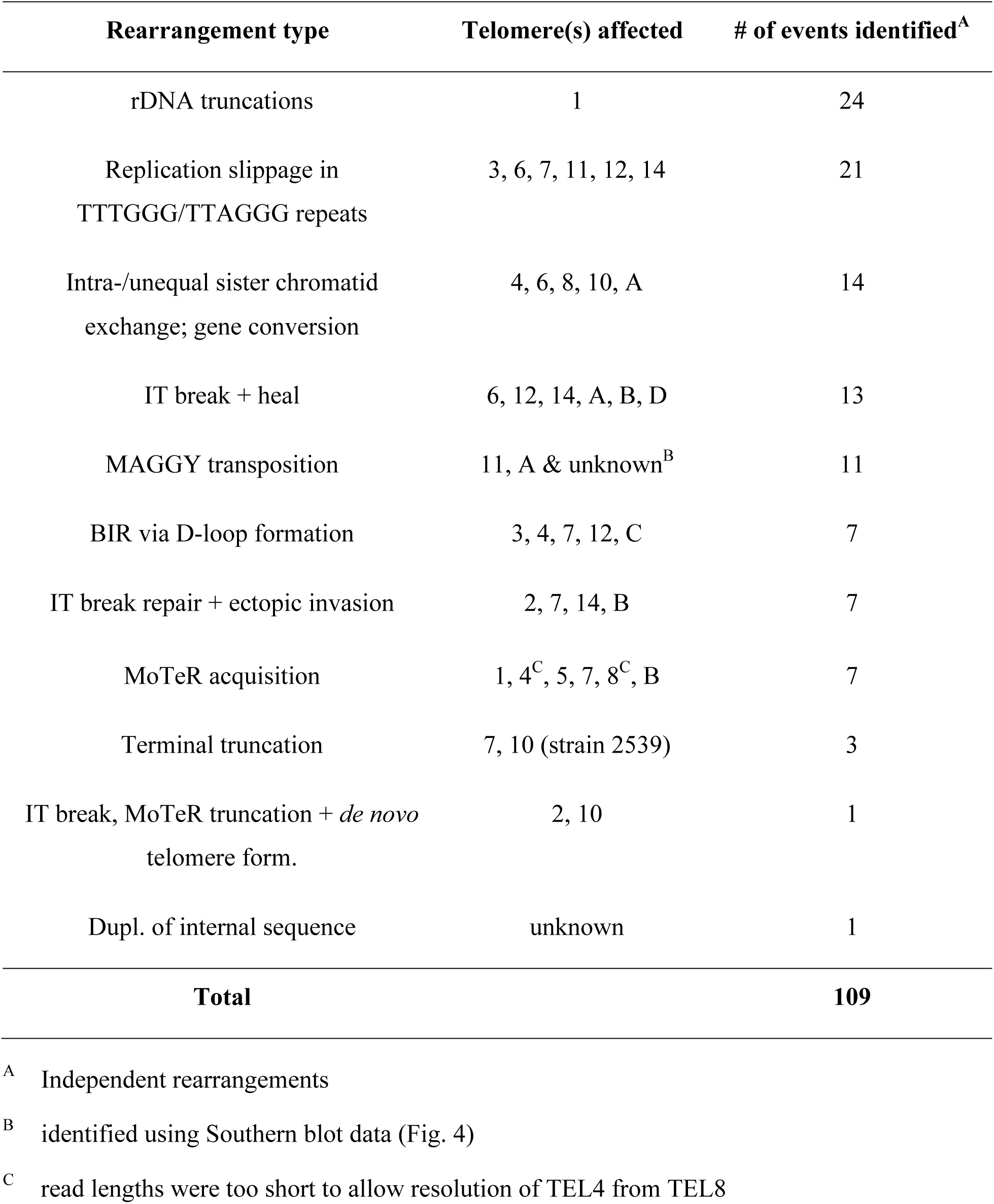
Rearrangements identified in this study

| Rearrangement type | Telomere(s) affected | # of events identified<sup>A</sup> |
| --- | --- | --- |
| rDNA truncations | 1 | 24 |
| Replication slippage in<br>TTTGGG/TTAGGG repeats | 3, 6, 7, 11, 12, 14 | 21 |
| Intra-/unequal sister chromatid<br>exchange; gene conversion | 4, 6, 8, 10, A | 14 |
| IT break + heal | 6, 12, 14, A, B, D | 13 |
| MAGGY transposition | 11, A & unknown<sup>B</sup> | 11 |
| BIR via D-loop formation | 3, 4, 7, 12, C | 7 |
| IT break repair + ectopic invasion | 2, 7, 14, B | 7 |
| MoTeR acquisition | 1, 4<sup>C</sup>, 5, 7, 8<sup>C</sup>, B | 7 |
| Terminal truncation | 7, 10 (strain 2539) | 3 |
| IT break, MoTeR truncation + *de novo*<br>telomere form. | 2, 10 | 1 |
| Dupl. of internal sequence | unknown | 1 |
| **Total** |  | **109** |

<sup>A</sup> Independent rearrangements
<sup>B</sup> identified using Southern blot data (Fig. 4)
<sup>C</sup> read lengths were too short to allow resolution of TEL4 from TEL8

## SUPPLEMENTAL FIGURE LEGENDS

**Fig. S1.**
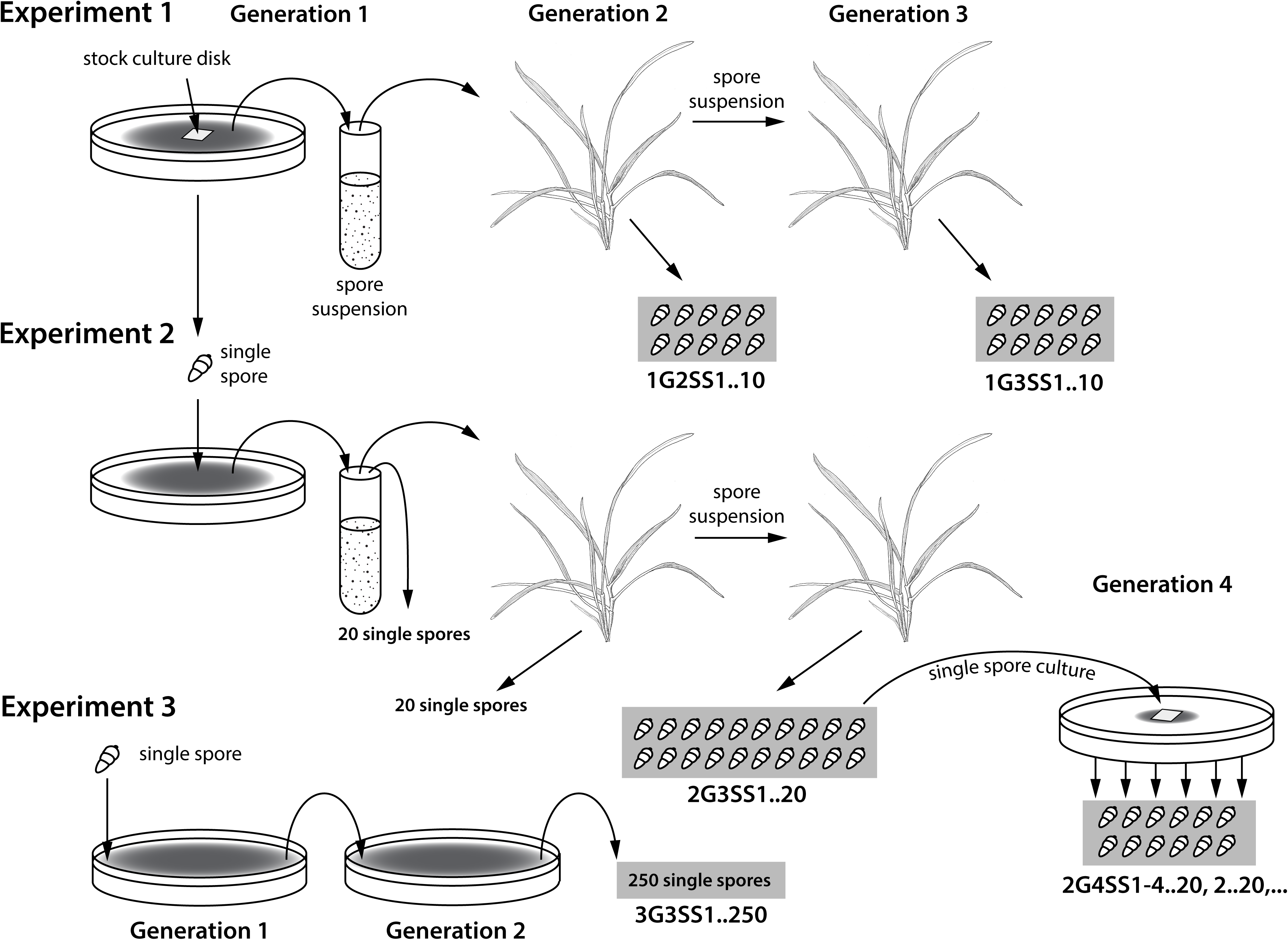
Single-spore generations used in this study. Note: populations used in this study are shown in gray boxes. *Experiment 1*. First generation spores of LpKY97 were harvested from oatmeal agar and used to inoculate plants of annual ryegrass cultivar Gulf. Second generation spores were harvested from lesions and used to inoculate a second batch of plants and a set of 10 single spore (SS) cultures was established. Third generation spores were collected from lesions and 10 third generation SS cultures were obtained. *Experiment 2*. A single spore of LpKY97 was used to inoculate an oatmeal agar plate and the resulting first spore generation was used to initiate two rounds of infection on perennial ryegrass cultivar Linn. Twenty SS isolates were established at each generation. Permanent stocks of several third generation, SS cultures were reactivated on oatmeal agar and up to 20 fourth generation SS cultures were generated. *Experiment 3.* The original LpKY97 SS culture generated in experiment 2 was serially cultured on complete medium agar by allowing it to grow across the full diameter of two 85 mm complete agar plates and 250 SS cultures were established.

**Fig. S2.**
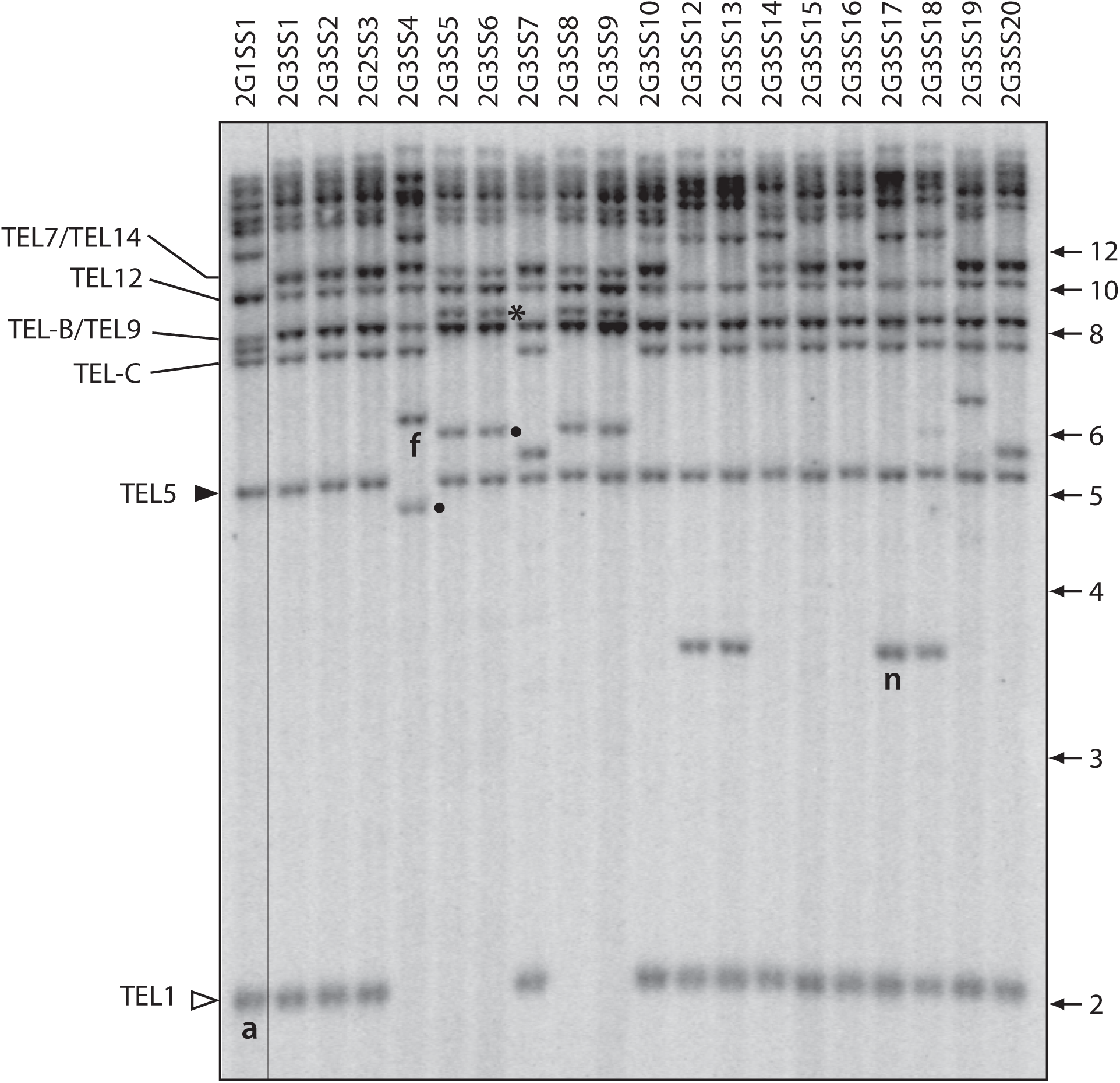
Telomere-rearrangements arising after successive generations of *in planta* growth, starting with a culture that originated from a single, genetically-purified spore. Telomeric restriction profiles of SS isolates from following successive rounds of plant infection (Fig. S1A). DNA samples were digested with *Pst*I, fractionated by gel electrophoresis, blotted to a membrane and hybridized with a telomere probe. The filled arrowhead marks the highly unstable “rDNA” telomere (TEL1), while the open arrowhead marks a stable telomere that rarely exhibited rearrangement (TEL5). Novel telomeric restriction fragments that were characterized and described here are labeled “a”, “f”, “n”, “o” and “p.” The asterisk marks a novel TRF that is shared by culture 5, 6, 8 and 9 and which arose through a MoTeR2 insertion in TEL-C - presumably via a transposition event (see Fig. S15B.iii).

**Fig. S3.**
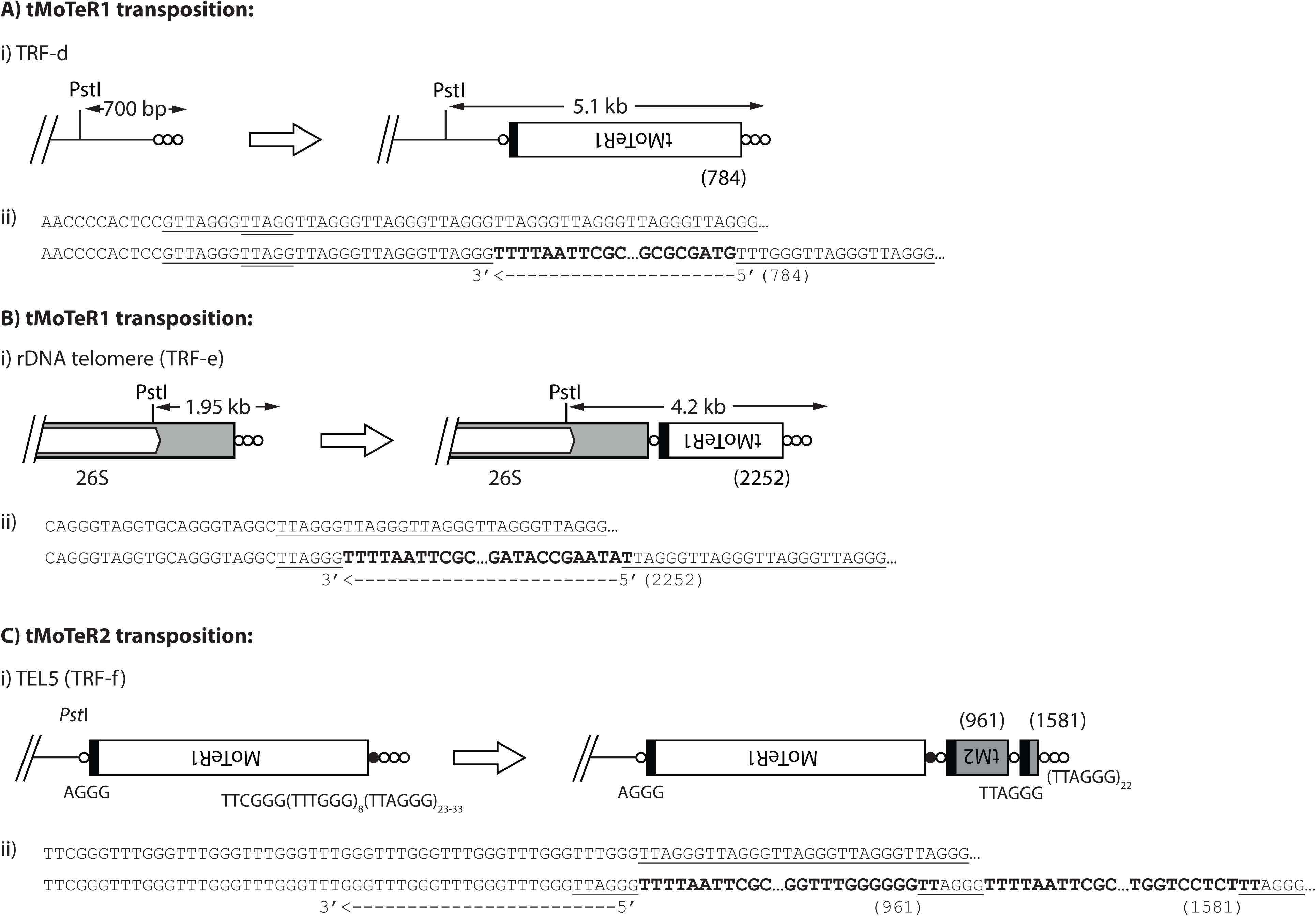
Target site analysis of putative MoTeR transposition events. A) i. Putative tMoTeR1 transposition event giving rise to TRF-d; ii. Sequence alignment showing tMoTeR insertion site pre- and post-insertion. B) i. Putative tMoTeR1 transposition event giving rise to TRF-e; ii. Sequence alignment of insertion site pre -and post-MoTeR insertion. C) i. Putative tMoTeR2 transposition events giving rise to TRF-f; ii) Sequence alignment of insertion site pre -and post-MoTeR insertion. MoTeR truncation positions are listed in parentheses.

**Fig. S4.**
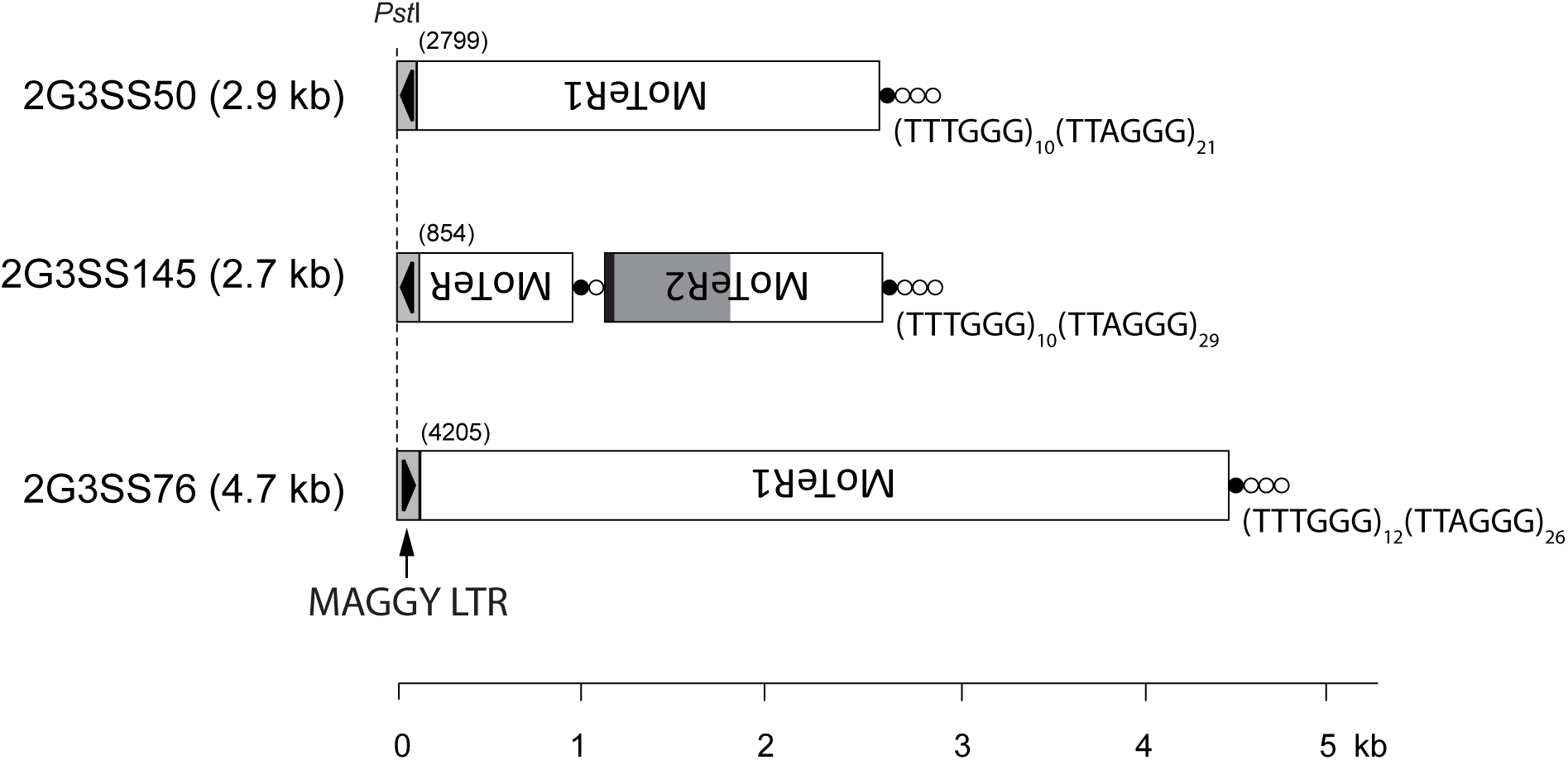
Telomeric restriction fragments representing *de novo* MAGGY insertions into MoTeR sequences. MoTeR segments are shaded as follows: white = present in MoTeR1; light gray = unique to MoTeR2; black = 3’ sequence shared by both elements. MAGGY LTR sequence is shown as a box with an arrow underneath indicating orientation, and the insertion positions are shown in parentheses.

**Fig. S5.**
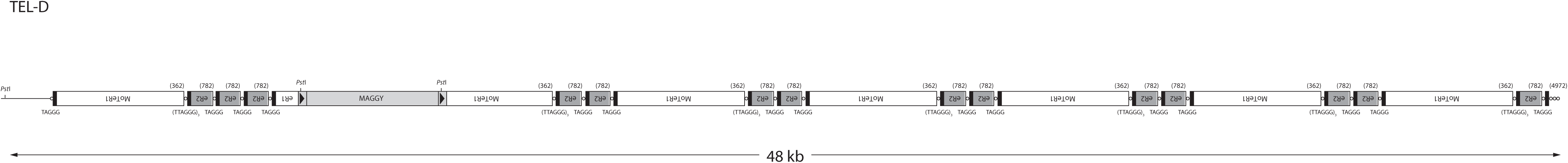
A single MoTeR array with a MAGGY insertion in the LpKY97 parental strain. The combined MinION assembly identified a single MAGGY insertion in a complex MoTeR array in TEL-D on minichromosome 2. Note: the entire array was captured in a single MinION read thereby confirming the validity of the array’s structure.

**Fig. S6.**
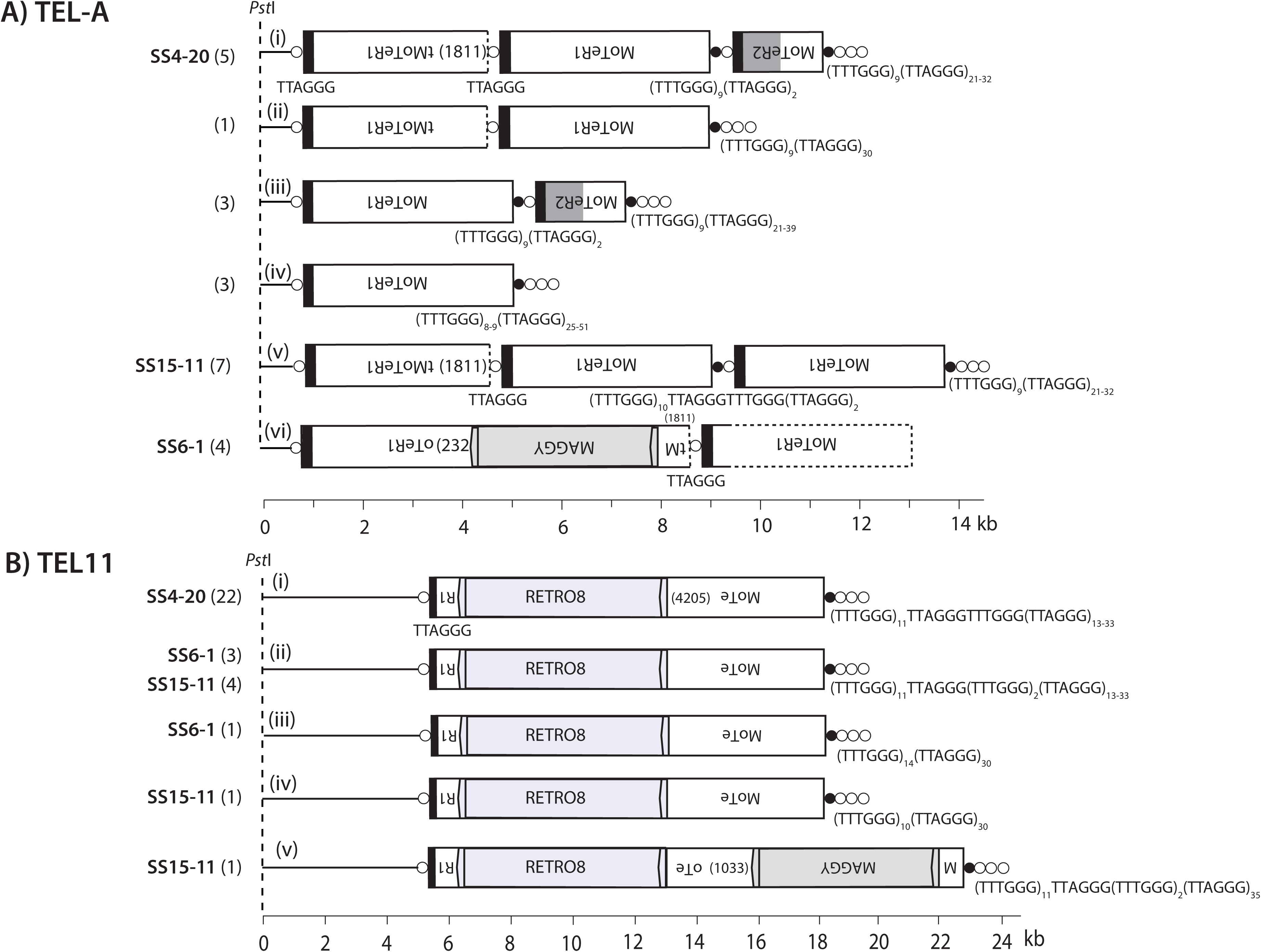
*De novo* MAGGY insertions in TEL11 and TEL-A. A) The parental form of TEL11 contained a single, full-length MoTeR1 insertion, which itself contained a super-insertion of a previously undiscovered retrotransposon, “RETRO8,” at nucleotide position 4205. A single MinION read from SS15-11 possessed a second super-insertion, consisting of a MAGGY integrated at nucleotide position 1033. LTRs are shown as block arrows at each end of RETRO8 and MAGGY, and the elements’ labels also reflect their orientations. Note the differences in variant telomere tracts among the different MinION reads. Inferred rearrangement mechanisms were as follows: length differences in variant telomere tracts in i, ii, iii, & iv) replication slippage; ii -> v) MAGGY insertion. B) Many different structural variants of TEL-A were found among the MinION read data for the three SS isolates. The original parental form had a truncated MoTeR1 in the proximal position. In SS6-1, this tMoTeR1 contained a MAGGY insertion (variant A.vi). B) TEL-A. Inferred rearrangement mechanisms: i -> ii) Interstitial telomere break & healing; i -> iii) intrachromatid recombination between MoTeR1 and tMoTeR1 copy; iii -> iv) Interstitial telomere break & healing; ii -> v) ectopic recombination; i or ii or v -> vi) MAGGY insertion

**Fig. S7.**
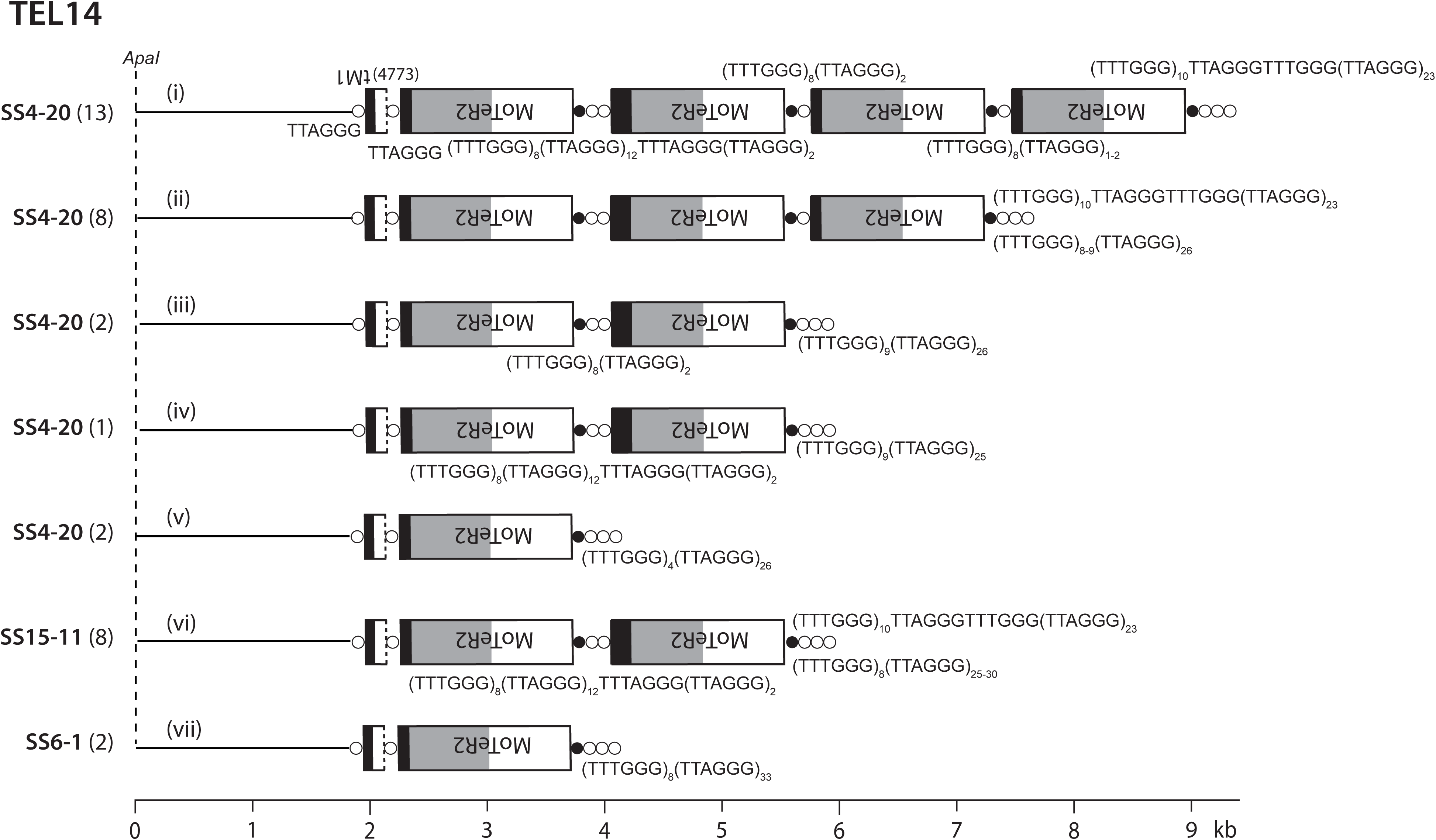
TEL14 variants identified among MinION reads. All structures were inferred from individual MinION reads that spanned the entire distance from the chromosome unique sequence to the telomere. Note the differences in variant telomere tracts at MoTeR junctions and subtending the telomeres proper. Inferred rearrangement mechanisms: i -> ii interstitial telomere break, repair via unequal sister chromatid exchange, replication slippage; i or ii -> iii) interstitial telomere break, repair via unequal sister chromatid exchange; i or ii to iv) interstitial telomere break plus slippage; i or ii or iii or iv -> v) interstitial telomere break & healing; i or ii to vi) interstitial telomere break, repair via unequal sister chromatid exchange.

**Fig. S8.**
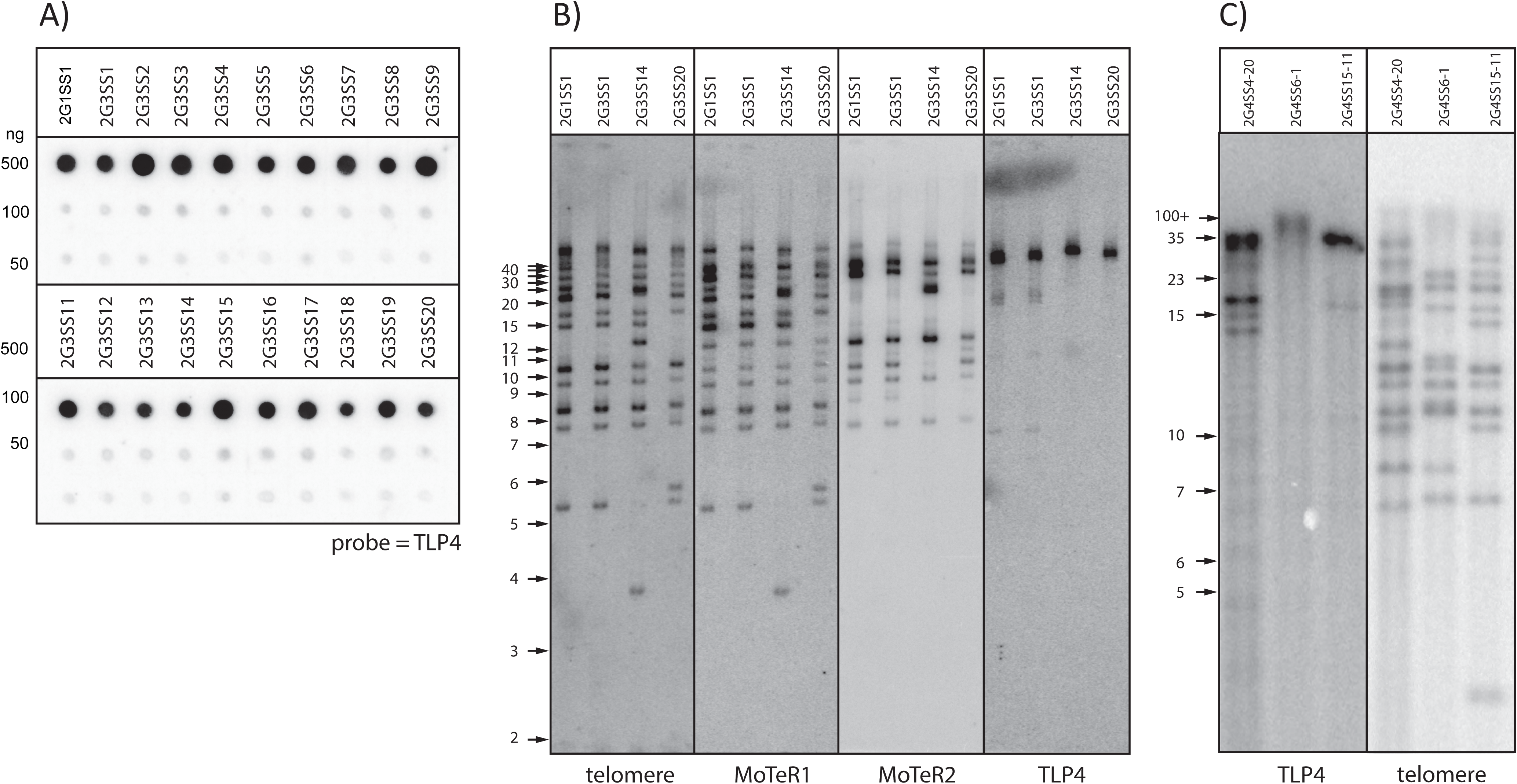
TLP4 spot blot and CHEF gel analysis of TLP4. A) Spot blot analysis of hybridization intensity in SS isolates. One microliter aliquots of genomic DNA sample from 1G1SS1 and 19 2G3SS isolates were spotted on a nylon membrane. The DNAs were denatured, neutralized and cross-linked, and the membrane was hybridized with ^32^P-labeled TLP-e. The resulting phosphorimage is shown. Note the similar hybridization intensities across spots when compared with the signals in Fig. 6. B) Analysis of TEL4 RFs in representative SS isolates. Agarose microbead-embedded genomic DNA was digested with *Pst*I, fractionated using CHEF electrophoresis, electroblotted to a nylon membrane and probed sequentially with telomere, TLP4, MoTeR1 and MoTeR2 probes. The resulting phosphorimages are shown. C). Analysis of TEL4 RFs in the DNA samples used for MinION sequencing. High molecular weight genomic DNA was digested with *Pst*I, fractionated on a 0.4% agarose gel, electroblotted to a nylon membrane and probed with the TLP4 probe. Note: equal amounts of DNA were loaded in each lane. Molecular sizes are in kilobases.

**Fig. S9.**
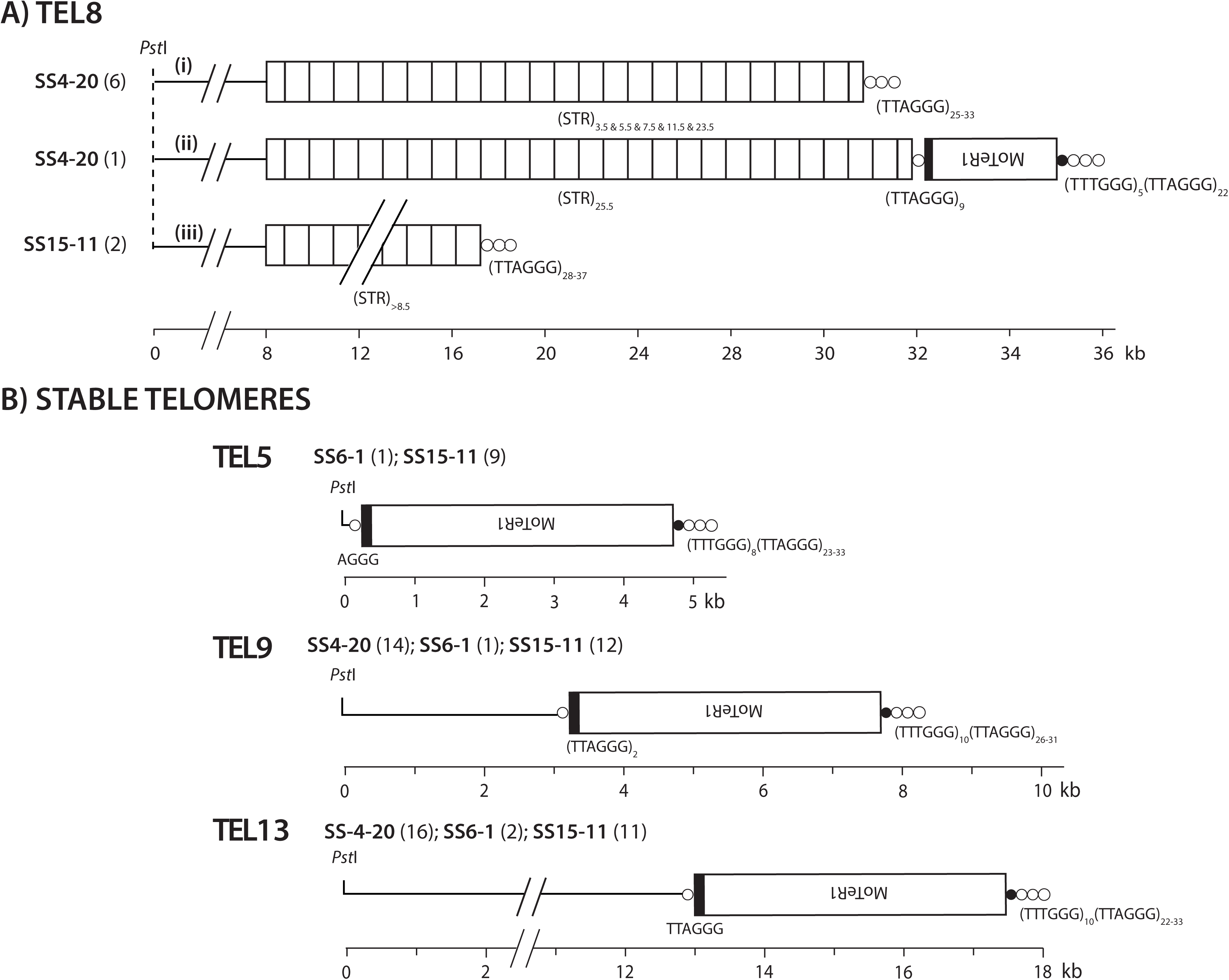
A) Alterations of the subtelomeric tandem repeat subtending TEL8; B) Structures of stable TELs 5, 9 and 14. The organization of the most terminal *Pst*I fragments was determined by subcloning and sequencing (TEL5) and by analysis of MinION reads (TELs 5, 9 and 14). No major structural variants were detected for TELs 9 and 14, although the numbers of variant TTTGGG repeats was different in some reads. All structures were inferred from individual MinION reads that spanned the entire distance from the chromosome unique sequence to the telomere. Inferred rearrangement mechanisms: STR copy number variation in i) intrachromatid unequal sister chromatid exchange; i -> ii) MoTeR1 transposition. B) Structures of stable telomeres. Note the absence of long interstitial telomeres, and the presence of TTCGGG/TTTGGG repeats subtending the telomere proper.

**Fig. S10.**
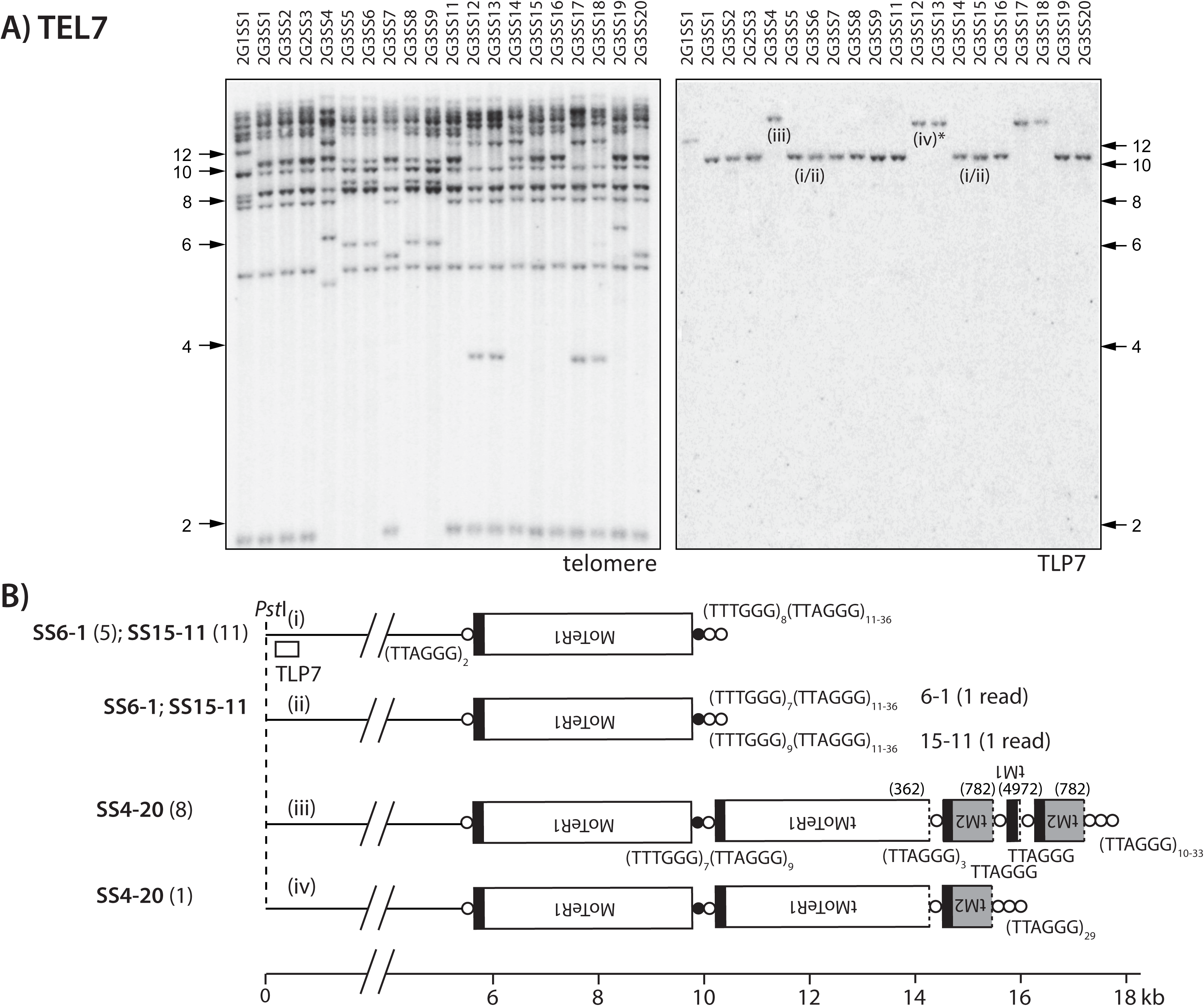
Interstitial telomere breaks at TEL7. A) Southern hybridization analysis of SS DNAs using TLP7. A Southern blot of *PstI*-digested SS DNAs was sequentially hybridized using the telomere and TLP7 probes, as shown below the respective phosphorimages. TEL7 variants are labeled i, ii, iii and iv. The asterisk is to indicate that this TRF is <u>predicted</u> to correspond to variant ii shown in B - a definitive link has not yet been established; B) Structures of the four variants as determined from MinION reads. All structures were inferred from individual MinION reads that spanned the entire distance from the chromosome unique sequence to the telomere. Inferred rearrangement mechanisms: i -> iii) transposition of a composite MoTeR cassette transcribed from the three distal elements in TEL-D (see Fig. S5), followed by transposition of a truncated MoTeR2 into the newly acquired telomere; iii -> iv) as above, but with resection/attrition and telomere healing, as an alternate pathway to transposition.

**Fig. S11.**
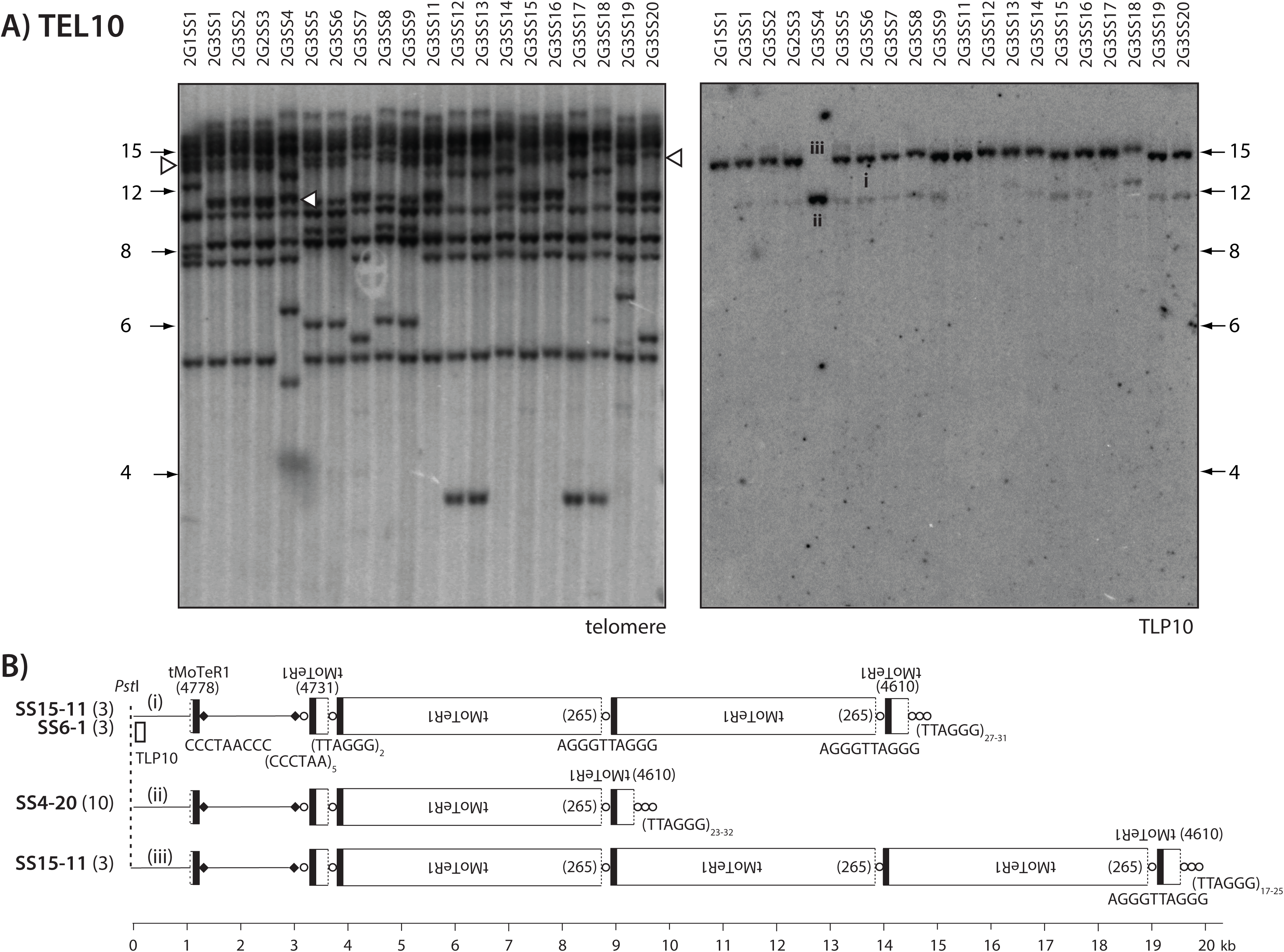
Unequal sister chromatid exchange at TEL10. A) Southern hybridization analysis of SS DNAs using TLP10. A Southern blot of *PstI*-digested SS DNAs was sequentially hybridized using the telomere and TLP10 probes, as shown below the respective phosphorimages. The TRFs that correspond to the three variants identified among MinION reads are labeled i, ii and iii. B) TEL10 structures identified among MinION reads. All structures were inferred from individual MinION reads that spanned the entire distance from the chromosome unique sequence to the telomere. Inferred rearrangement mechanism: i -> ii & iii) Unequal sister chromatid exchange.

**Fig. S12.**
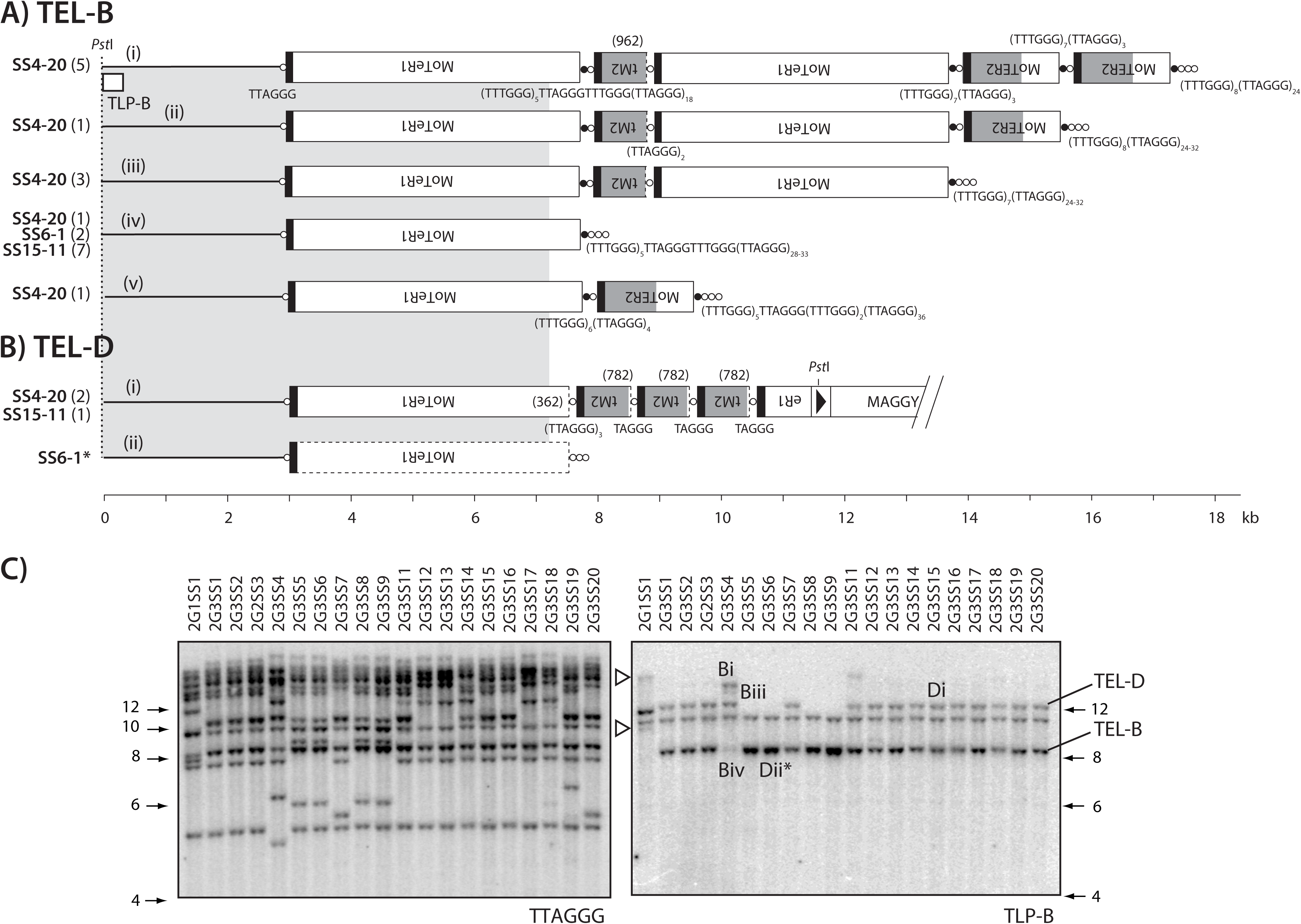
Telomere rearrangements at minichromosome TELs B and D. A) Structural organization of TRF-B variants showing the origin of TLP-B. Inferred rearrangement mechanisms: i -> ii) Interstitial telomere break & healing; i or ii -> iii) Interstitial telomere break & healing; i or ii or iii -> iv) Interstitial telomere break & healing; iv -> v) MoTeR2 transposition. B) Structural organization of TRF-D variants. The subtelomere region with sequence identity to TEL-B is highlighted using a gray-shaded background. Inferred rearrangement mechanism: i -> ii) Interstitial telomere break & healing. All structures were inferred from individual MinION reads that spanned the entire distance from the chromosome unique sequence to the telomere. C) Southern hybridization analysis of SS DNAs using TLP-B. A blot of *PstI*-digested SS DNAs was sequentially hybridized using the telomere and TLP-B probes, as shown below the respective phosphorimages. TEL-B and TEL-D variants that were detectable using the TLP-B probe are numbered i, ii, iii, and iv, with given the relevant B & D prefix. A white arrowhead marks the presumed B TRF variants in the 2G1SS1 starting culture.

**Fig. S13A.**
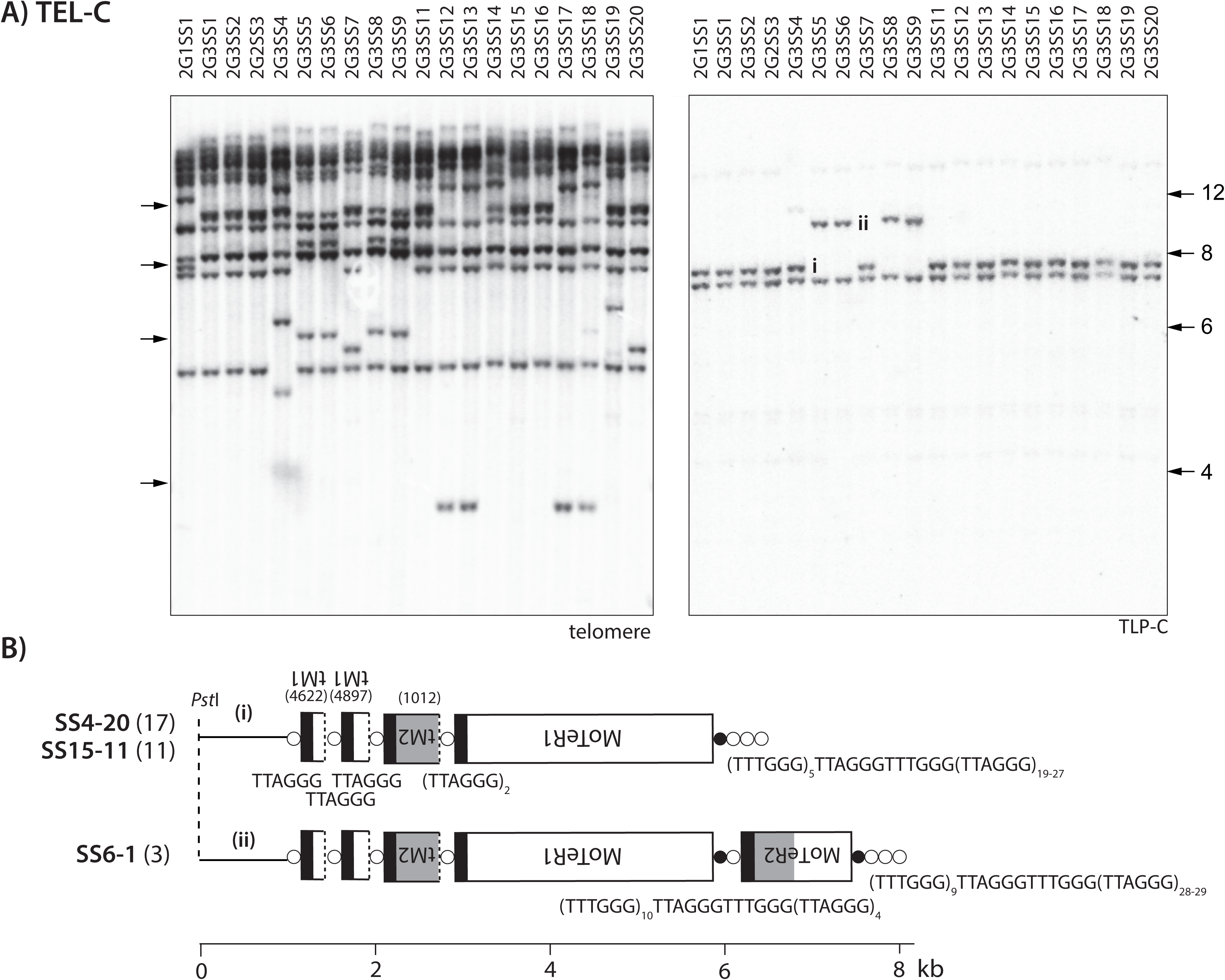
TEL-C structural variants identified using MinION reads. Inferred rearrangement mechanisms: i -> ii) Interstitial telomere break & ectopic recombination (unknown origin). With the exception of TEL-A variant v, all structures were inferred from individual MinION reads that spanned the entire distance from the chromosome unique sequence to the telomere.

**Fig. S14.**
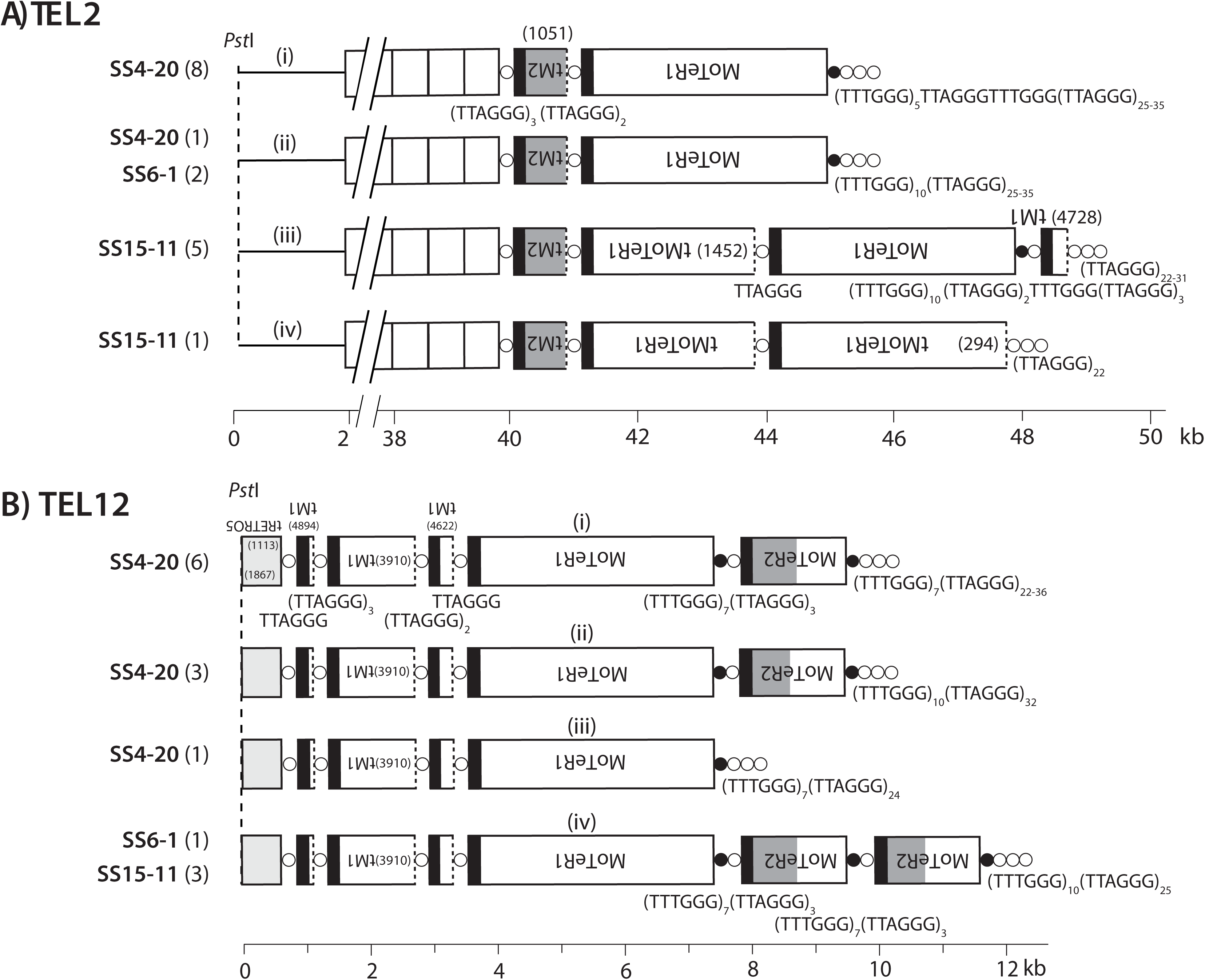
Structural variants of TEL2 and TEL12 identified using MinION reads. A) TEL2. Inferred rearrangement mechanisms: ii -> i) gene conversion with TEL-B (see Fig. S12A, variant SS4-20.iv); ii) -> iii) ectopic recombination with TEL4 (see Fig. 6B, variant SS6-1.v); i -> iv) interstitial telomere break, resection, *de novo* telomere formation. B) TEL12. Inferred rearrangement mechanisms: i or ii -> iii) interstitial telomere break & healing; i -> iv) D-loop formation and extension. All structures were inferred from individual MinION reads that spanned the entire distance from the chromosome unique sequence to the telomere.

**Fig. S15.**
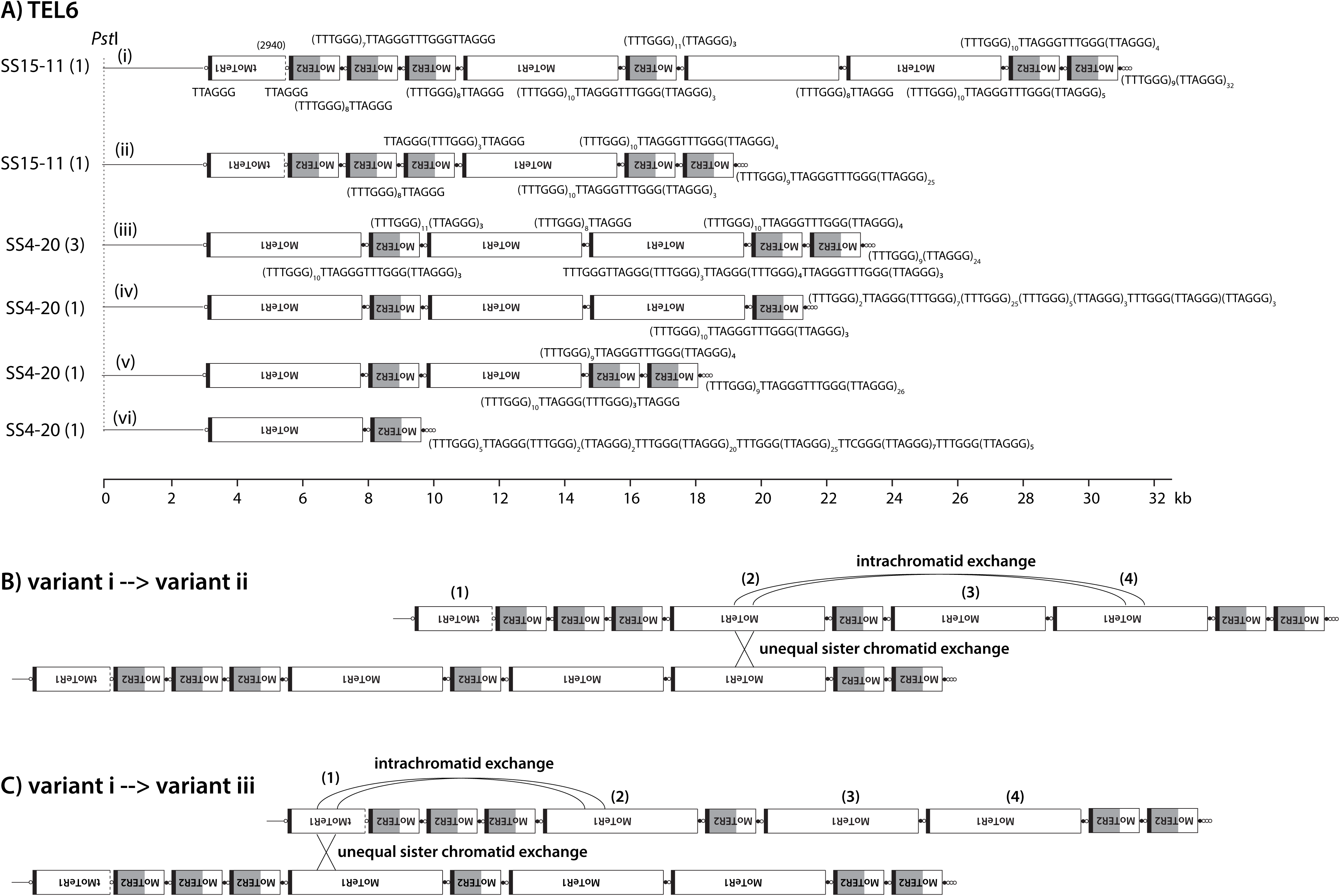
Structural variants of TEL6 identified using MinION reads. All structures were inferred from individual MinION reads that spanned the entire distance from the chromosome unique sequence to the telomere. Note the highly unusual variant telomere tracts separating the MoTeR copies and subtending the telomere proper. Inferred rearrangement mechanisms: i -> ii & i -> iii) intrachromatid/unequal sister chromatid exchange & slippage; iii - iv) interstitial telomere break and healing & slippage; iii or iv or v -> vi) interstitial telomere break and healing & slippage; i or ii or iii -> v) intrachromatid/unequal sister chromatid exchange & slippage.

## Notes

https://www.ncbi.nlm.nih.gov/bioproject/579424

